# Optimal Filters for ERP Research II: Recommended Settings for Seven Common ERP Components

**DOI:** 10.1101/2023.06.13.544794

**Authors:** Guanghui Zhang, David R. Garrett, Steven J. Luck

**Affiliations:** Center for Mind and Brain, University of California–Davis, Davis, California, 95618, USA

**Author notes:** Corresponding author Email address; Address:267 Cousteau Place, Davis, CA 95618, USA. (Guanghui Zhang).

**Keywords:** ERP, optimal filtering, noise level, SNR, wave distortion, scoring methods

## Abstract

In research with event-related potentials (ERPs), aggressive filters can substantially improve the signal-to-noise ratio and maximize statistical power, but they can also produce significant waveform distortion. Although this tradeoff has been well documented, the field lacks recommendations for filter cutoffs that quantitatively address both of these competing considerations. To fill this gap, we quantified the effects of a broad range of low-pass filter and high-pass filter cutoffs for seven common ERP components (P3b, N400, N170, N2pc, mismatch negativity, error-related negativity, and lateralized readiness potential) recorded from a set of neurotypical young adults. We also examined four common scoring methods (mean amplitude, peak amplitude, peak latency, and 50% area latency). For each combination of component and scoring method, we quantified the effects of filtering on data quality (noise level and signal-to-noise ratio) and waveform distortion. This led to recommendations for optimal low-pass and high-pass filter cutoffs. We repeated the analyses after adding artificial noise to provide recommendations for datasets with moderately greater noise levels. For researchers who are analyzing data with similar ERP components, noise levels, and participant populations, using the recommended filter settings should lead to improved data quality and statistical power without creating problematic waveform distortion.

## 1. Introduction

Almost all ERP studies involve filtering, and optimal filter settings are important to maximize statistical power and avoid waveform distortions that might lead to erroneous conclusions(Tanner et al., 2015, 2016; Acunzo et al., 2012; Rousselet, 2012; van Driel et al., 2021; VanRullen, 2011; Yeung et al., 2007). However, there is no widespread consensus about what filter settings are optimal, and the optimal settings are likely to vary across scoring methods, experimental paradigms, subject populations, and recording setups.

Filter settings vary widely across published studies, even studies within the same research domain. For example, in a review of emotional face processing studies, Schindler & Bublatzky (2020) found that studies used high-pass cutoff frequencies ranging from 0.01 Hz to 1.0 Hz and low-pass cutoff frequencies ranging from 16 to 200 Hz. Formal recommendations for filter settings also vary widely. For example, a set of recommendations for clinical ERP studies (Duncan et al., 2009) recommended a bandpass of 1–30 Hz for the mismatch negativity and 0.1–100 Hz for the P3 and N400 components (with the possibility of a low-pass cutoff of 12-15 Hz for visualization). We have previously recommended a bandpass of 0.1–30 Hz for most cognitive and affective studies (Luck, 2014). In most of these cases, little or no justification was provided for the filter cutoffs; when justifications were provided, they were usually based on informal or qualitative considerations rather than formal, quantitative considerations. Given that different filter cutoffs lead to different levels of statistical power and waveform distortion, the field would be well served by a set of careful, mathematically rigorous, and empirically justified recommendations for filter settings that take into account the details of a given study (e.g., the component being studied, the scoring method).

In a companion paper (Zhang et al., 2023), we presented a new approach for determining the optimal filter settings for a given situation. This approach involves filtering a dataset with many different filters and then assessing the resulting noise level and the signal-to-noise ratio (SNR) for each filter. Each filter is also applied to noise-free simulated ERP waveforms to quantify the amount of waveshape distortion produced by the filter. The optimal filter is chosen as the one that produces the best data quality without exceeding a preset criterion for waveshape distortion. The present paper applied this approach to an open EEG/ERP dataset — the ERP CORE (Kappenman et al., 2021) —to determine optimal filter settings for a broad set of ERP components obtained from a relatively large set of healthy young adults. These settings should also be optimal, or nearly optimal, for similar paradigms, participant populations, and recording conditions. In addition, we repeated the analyses after adding more noise to the data to determine optimal filters for modestly noisier datasets. The recommended filter cutoffs are shown in Table 1.

**Table 1:**
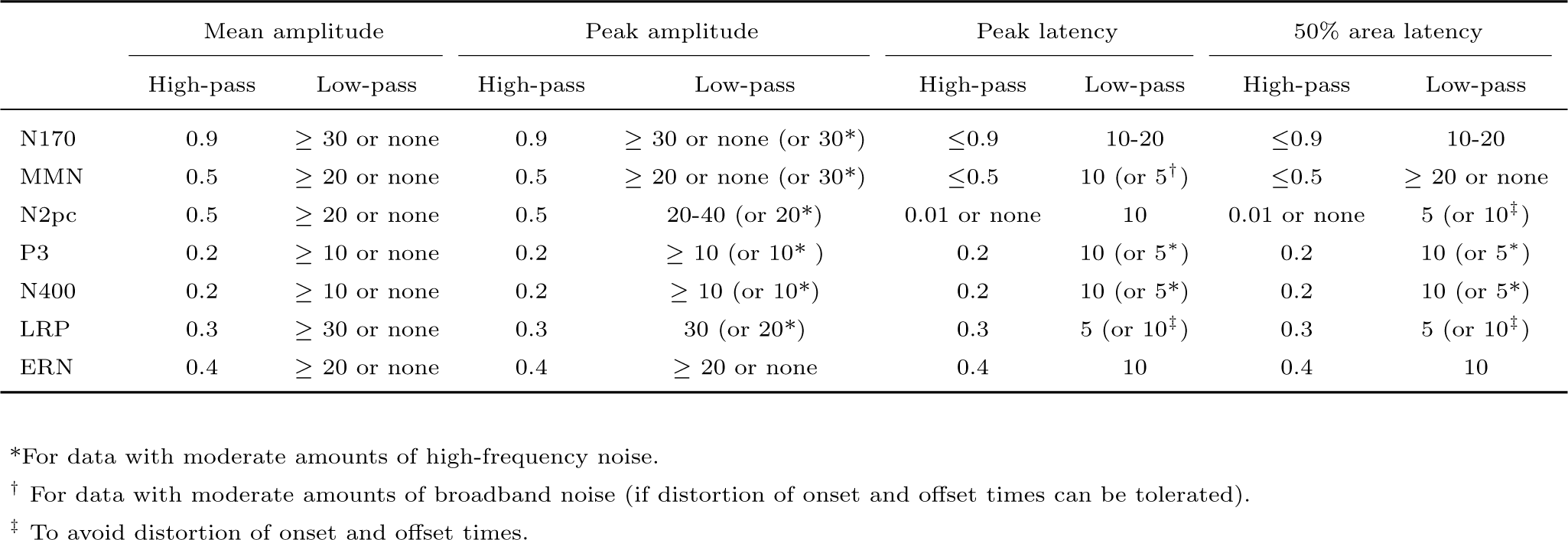
Recommended filter settings (in Hz, with a slope of 12 dB/octave) for each combination of scoring method and ERP component.

|  | Mean amplitude |  | Peak amplitude |  | Peak latency |  | 50% area latency |  |
| --- | --- | --- | --- | --- | --- | --- | --- | --- |
|  | High-pass | Low-pass | High-pass | Low-pass | High-pass | Low-pass | High-pass | Low-pass |
| N170 | 0.9 | $\geq 30$ or none | 0.9 | $\geq 30$ or none (or 30*) | $\leq 0.9$ | 10-20 | $\leq 0.9$ | 10-20 |
| MMN | 0.5 | $\geq 20$ or none | 0.5 | $\geq 20$ or none (or 30*) | $\leq 0.5$ | 10 (or 5<sup>†</sup>) | $\leq 0.5$ | $\geq 20$ or none |
| N2pc | 0.5 | $\geq 20$ or none | 0.5 | 20-40 (or 20*) | 0.01 or none | 10 | 0.01 or none | 5 (or 10<sup>‡</sup>) |
| P3 | 0.2 | $\geq 10$ or none | 0.2 | $\geq 10$ (or 10*) | 0.2 | 10 (or 5*) | 0.2 | 10 (or 5*) |
| N400 | 0.2 | $\geq 10$ or none | 0.2 | $\geq 10$ (or 10*) | 0.2 | 10 (or 5*) | 0.2 | 10 (or 5*) |
| LRP | 0.3 | $\geq 30$ or none | 0.3 | 30 (or 20*) | 0.3 | 5 (or 10<sup>‡</sup>) | 0.3 | 5 (or 10<sup>‡</sup>) |
| ERN | 0.4 | $\geq 20$ or none | 0.4 | $\geq 20$ or none | 0.4 | 10 | 0.4 | 10 |

\*For data with moderate amounts of high-frequency noise.
<sup>†</sup> For data with moderate amounts of broadband noise (if distortion of onset and offset times can be tolerated).
<sup>‡</sup> To avoid distortion of onset and offset times.

### 1.1. Overview of the approach

The companion paper (Zhang et al., 2023) provides a detailed description and justification of our approach. This section provides a brief overview.

Our approach is based on the observation that the effects of a particular type of noise depend on the method used to obtain the amplitude or latency scores that will be entered into statistical analyses and used to test the scientific hypotheses of a given study (e.g., the peak amplitude of the N170 component). For example, high-frequency noise has a large impact on peak amplitude scores but relatively little impact on mean amplitude scores. As a result, a low-pass filter might be more beneficial for peak amplitude scores than for mean amplitude scores. Therefore, it is important to assess how a filter will impact the noise level and/or SNR for a particular scoring method. In addition, costs and benefits of filtering will depend on the shape of the component being analyzed, so it is also important to assess the performance of a filter for a variety of different components.

When the SNR is computed with our approach, the specific amplitude or latency score for a specific component defines the signal, and the expected error in this score defines the noise. The signal can be quantified by obtaining the score of interest from a grand average waveform (to minimize contributions from noise). The noise that is relevant for that specific score can be assessed using a new metric of ERP data quality called the standardized measurement error (SME; Luck et al. (2021); Zhang & Luck (2023)). The SME is obtained from individual participants and then aggregated across participants by computing the root mean square (RMS) of the single-participant SME values. The SNR is then calculated as the score divided by the RMS(SME) of that score. We use the term *SNR_SME_* to indicate this specific definition of the SNR.

Filters typically reduce the amplitude of an ERP component, and it is therefore important to determine whether a given filter decreases the noise more than it decreases the amplitude. For this reason, it is essential to use the *SNR_SME_*when determining the optimal filter settings for amplitude scores. However, most filters have only a modest effect on latency values, and they do not typically reduce the difference in latency between conditions (see Zhang et al. (2023)). Thus, when determining the optimal filter settings for latency scores, we do not look at the ratio of signal to noise for each filter but instead look at the noise level directly using the RMS(SME).

Filters are a form of controlled distortion, and they temporally “smear” the ERP waveform. The smearing produced by high-pass filters is inverted, so these filters typically produce artifactual opposite-polarity peaks before and after a true peak. Low-pass filters with steep roll-offs may also produce these artifactual opposite-polarity peaks. Our approach to filter selection involves applying filters to noise-free simulated ERP waves and quantifying the size of the artifactual peaks. Simulated data must be used for this purpose because ground truth is not known for real data, making it impossible to know whether a given filter has created an artifactual peak or “revealed” a true peak that would be clearly visible in the absence of noise. The size of the artifactual peaks produced by a given filter is quantified as the amplitude of the artifactual peak relative to the amplitude of the true peak (the artifactual peak percentage). Low-pass filters also cause ERP onset latencies to become earlier and offset latencies to shift later, which can be quantified as the percentage change in the width of the filtered waveform relative to the unfiltered waveform (the temporal distortion percentage).

The amount of waveform distortion that can be tolerated depends on the scientific goals of a given study. In most cases, artifactual opposite-polarity peaks are the most significant kind of distortion, because they can cause a modulation of one component to be misinterpreted as a modulation of some other component. For example, extreme high-pass filtering can cause a true P600 effect to produce an artifactual N400-like effect (Tanner et al., 2015). In the companion paper (Zhang et al., 2023), we argued that researchers should avoid filters that produce an artifactual peak that is more than 5% of the size of the true peak, at least for studies conducted in highly cooperative participant populations using high-quality recording systems. Thus, the optimal filter in such studies is defined as the one that yields the best SME or *SNR_SME_* while producing an artifactual peak percentage that does not exceed 5%. A different threshold may be appropriate for other kinds of studies.

The effects of filtering on onset and offset latencies are not usually a major problem because all groups and conditions are typically impacted equally. Consequently, we do not ordinarily use the temporal distortion percentage in determining optimal filter parameters. However, the amount of temporal distortion would be relevant in cases where the absolute onset time of an effect is theoretically relevant.

### 1.2. Goals of the present study

Although our approach is straightforward, and we have provided tools to make it easy to implement in ERPLAB Toolbox (Lopez-Calderon & Luck, 2014), applying this approach to a broad set of data requires an investment of time. Moreover, filter settings should be set a priori, so previously collected data must be available. The present study therefore applied this approach to determine optimal filter settings for a broad set of ERP components. We used the data from the ERP CORE (Compendium of Open Resources and Experiments; Kappenman et al. (2021)), which includes data from 40 young adults who performed six standardized paradigms that yielded seven commonly-studied ERP components: P3b, N400, N170, N2pc, mismatch negativity (MMN), error-related negativity (ERN), and lateralized readiness potential (LRP). We provide recommended filter settings for each of these components, separately for four different scoring methods: mean amplitude, peak amplitude, peak latency, and 50% area latency (Clayson et al., 2013; Luck, 2014). The recommended filter settings should be useful for a broad range of researchers who record ERPs from similar subject populations using similar recording setups.

The ERP CORE data contain relatively low levels of high-frequency noise because the EEG was recorded using a high-quality recording system with active electrodes, a gel conductor, and a climate-controlled shielded chamber, and the participants were highly compliant young adults who were able to sit still. To demonstrate that our recommendations generalize to data that have modestly greater noise levels, we provide additional analyses in which different types of noise are added to the data. However, we do not expect the results to generalize to highly different subject populations (e.g., infants), very dissimilar recording setups (e.g., dry electrodes), or components with very different waveshapes (e.g., the auditory brainstem responses or contralateral delay activity). We invite other researchers to publish papers describing the optimal filter settings for these situations.

## 2. Method

All data processing was conducted in Matlab using EEGLAB Toolbox (Delorme & Makeig, 2004) and ERPLAB Toolbox (Lopez-Calderon & Luck, 2014). The data and scripts used in our analyses are available at https://osf.io/z3hfp/. If desired, the processing steps can be carried out using the graphical user interface rather than using scripts (using version 9.20 or higher of ERPLAB Toolbox).

### 2.1. ERP CORE Data and Preprocessing

The ERP CORE (Kappenman et al., 2021) data were downloaded from the online repository (https://doi.org/10.18115/D5JW4R). Details of the participants, paradigms, recording methods, and analysis procedures can be found in the original paper. Here, we provide a brief overview of the participants, recording methods, and preprocessing procedures. A brief description of each individual ERP paradigm is provided in the corresponding portion of the Results section.

The ERP CORE contains data from 40 neurotypical college students (25 female) who were recruited from University of California, Davis community. We used all 40 participants in the present analysis, irrespective of the number of artifacts or behavioral errors. The one exception is that one participant had zero usable trials for one condition of the N2pc experiment and was excluded from the analyses of that experiment.

The EEG was recorded using a DC-coupled Biosemi ActiveTwo recording system (Biosemi B.V., Amsterdam, the Netherlands) with active electrodes, an antialiasing filter (fifth order sinc filter, half-power cutoff at 204.8 Hz) and a 1024 Hz sampling rate. Although data are available for 30 scalp sites, along with horizontal and vertical electrooculogram electrodes, the present analyses focused on the single site where a given component was largest (see Table 2).

**Table 2:**
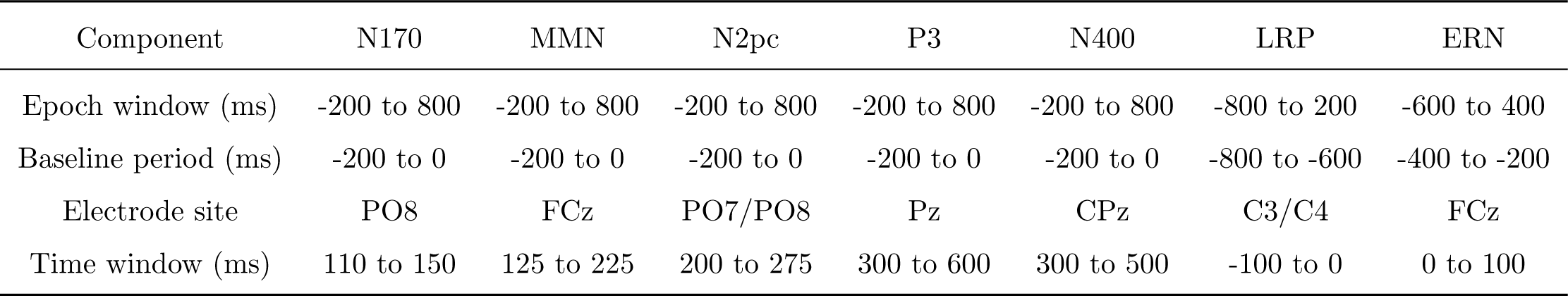
Epoch window, baseline period, electrode site, and time window used for each ERP component.

| Component | N170 | MMN | N2pc | P3 | N400 | LRP | ERN |
| --- | --- | --- | --- | --- | --- | --- | --- |
| Epoch window (ms) | -200 to 800 | -200 to 800 | -200 to 800 | -200 to 800 | -200 to 800 | -800 to 200 | -600 to 400 |
| Baseline period (ms) | -200 to 0 | -200 to 0 | -200 to 0 | -200 to 0 | -200 to 0 | -800 to -600 | -400 to -200 |
| Electrode site | PO8 | FCz | PO7/PO8 | Pz | CPz | C3/C4 | FCz |
| Time window (ms) | 110 to 150 | 125 to 225 | 200 to 275 | 300 to 600 | 300 to 500 | -100 to 0 | 0 to 100 |

The data used here had already passed through several steps of the ERP CORE preprocessing pipeline. This pipeline began by shifting the event codes to reflect the intrinsic delay of the video monitor and resampling the data at 256 Hz with an antialiasing filter at 115 Hz. The data were then referenced to the average of the P9 and P10 electrodes, except that the average of all scalp sites was used as the reference for the N170 paradigm. Independent component analysis (ICA) was used to correct the data for eye blinks and eye movements. A noncausal Butterworth high-pass filter (half-amplitude cutoff 0.1 Hz, 12dB/oct roll-off) was applied to optimize the ICA decomposition, and the component weights were then transferred back to the original data so that the artifacts could be corrected in the unfiltered data (see Chapter 9 in Luck (2022)).

### 2.2. Filtering and additional preprocessing

To assess the effects of filtering on the SME and *SNR_SME_*, we separately filtered the continuous EEG data resulting from the preceding steps using a variety of different combinations of low-pass and high-pass cutoffs. All cutoff frequencies listed in this paper indicate the half-amplitude point in the frequency response function. We used noncausal Butterworth filters (Hamming, 1998) because they are efficient, flexible, well-behaved, and widely used for EEG and ERP signals. The companion paper (Zhang et al., 2023) indicated that a relatively gentle slope of 12 dB/octave was optimal for minimizing waveform distortions, so we used this roll-off for all analyses. However, it would be straightforward to repeat the analyses with a different slope using the data and scripts we have provided at https://osf.io/z3hfp/. It would also be straightforward to repeat the analyses with other families of filters (e.g., finite impulse response filters).

After the data were filtered, we implemented the remaining preprocessing steps used in the original ERP CORE paper. First, we epoched and baseline-corrected the EEG using the time windows shown in Table 2. Then, we averaged the EEG epochs, excluding trials with artifacts or incorrect behavioral responses.

It is possible that filtering could influence the SNR of a given dataset indirectly by impacting the artifact rejection and correction processes. However, it is possible and even advisable to use different filtering parameters for determining which trials to reject, for performing the ICA decomposition, and for averaging and scoring the ERP amplitudes and latencies (Luck, 2014). Thus, the question of what filter is optimal for the primary ERP analyses is independent of the question of what filter is optimal for artifact rejection or correction. To keep these issues separate, and to focus on the effects of filtering on the primary ERP scores, we performed artifact rejection and correction in a manner that was not influenced by the different filter settings being examined in the main analyses. Specifically, we applied the rejection flags and ICA decompositions from the original ERP CORE data to the present data, independent of the filter settings being tested. The ERP CORE repository (https://doi.org/10.18115/D5JW4R) provides the artifact rejection and correction parameters, along with the number of included and excluded trials.

### 2.3. Computing the signal, noise, and signal-to-noise ratio

We computed the signal, noise, and signal-to-noise ratio separately for each combination of the seven ERP components and four scoring methods: mean amplitude, peak amplitude, peak latency, and 50% area latency. As justified in the companion paper (Zhang et al., 2023), the signal, noise, and signal-to-noise ratio were computed from the difference waves for a given component (e.g., the faces-minus-cars difference wave for the N170 component). However, only the noise values were relevant for the latency measures (see Section 1.1).

Table 2 shows the time windows and electrode sites that were used for each component. Mean amplitude was defined as the mean voltage during the time window. Peak amplitude was defined as the maximal positive voltage (for the P3 component) or the maximal negative voltage (for the other components) in that time window, and peak latency was defined as the latency at which that peak occurred. The 50% area amplitude was computed by taking the integral of the voltages during that time window and finding the point that bisected that integral into equal halves. To increase temporal precision, spline interpolation was used to upsample the waveforms by a factor of 10 before the latencies were scored (see Luck (2014) for the rationale). The signal was estimated by obtaining a given score from the grand average difference waveform for a given component.

We used the RMS(SME) for a given score to estimate the noise. The SME was obtained separately from each participant, and values were aggregated across participants by computing the root mean square (RMS) of the single-participant values (see the companion paper for the rationale). The aggregate *SNR_SME_*was then computed as the score from the grand average divided by the RMS(SME) of the single-participant scores.

When the amplitude of a component is scored as the mean voltage within a given time range, there is a simple analytic equation that can be used to estimate the SME (which is then called the analytic SEM or aSME). Moreover, ERPLAB Toolbox can directly compute aSME values at the time of averaging. To obtain the mean amplitude SME values for difference scores (e.g., faces minus cars), the aSME values were computed directly by ERPLAB for each parent waveform (e.g., separately for faces and cars), and the SME of the difference was calculated as *SME_A−B_* = ^✓^*SME* ^2^ + *SME_B_*^2^. In this equation, *SME_A−B_* is the SME of the difference between conditions A and B, and *SME_A_* and *SME_B_* are the SMEs of the two individual conditions (see the companion paper for details).

This approach to estimating the SME for a difference between conditions is not valid for differences between electrode sites, such as the contralateral-minus-ipsilateral difference waves used to isolate the N2pc component and the lateralized readiness potential. For these components, we used bootstrapping to estimate the SME (which is then called the bootstrapped SME or bSME) directly from the difference wave. The bootstrapping process is described in detail by Luck et al. (2021) and currently requires simple scripting (see example bSME scripts at https://doi.org/10.18115/D58G91). The scripts used to compute bSME values and the associated *SNR_SME_*values for the present study are provided at https://osf.io/z3hfp/. We used 1,000 bootstrap iterations for each bSME value.

The analytic SME is always inappropriate for scoring methods other than mean amplitude, so we also used bootstrapping to compute bSME values for peak amplitude, peak latency, and 50% area latency.

### 2.4. Effects of increased noise

The ERP CORE data are quite clean, especially with respect to line noise. To determine whether the results of the present study generalize to noisier data, we repeated all the analyses after adding a random-phase 60 Hz sinusoidal oscillation (to simulate line noise), white noise (to simulate muscle activity and other kinds of spiky noise), and pink noise (to simulate movement artifacts and other kinds of nonstationary noise). The noise was added to the continuous EEG prior to filtering. The 60 Hz oscillating noise had a peak-to-peak amplitude of 20 µV. The white noise and pink noise had a mean of zero and a standard deviation of 7.07 µV (which was the same as the standard deviation of the line noise). Supplementary Figure S1 provides an example of the EEG after the addition of these types of noise, showing that the noise was moderate relative to the original EEG. Researchers can modify the scripts at https://osf.io/z3hfp/ if they wish to see the effects of more extreme noise.

### 2.5. Quantifying waveform distortion

To quantify the distortion produced by a given filter, we applied the filter to an artificial waveform that was designed to simulate the grand average difference wave for a given component (e.g., the faces-minus-cars difference wave for the N170). As described in the companion paper (Zhang et al., 2023), the artificial waveforms were Gaussian or ex-Gaussian functions. The grand average difference waves are only an approximation of the true shape of the single-participant ERP components, so we did not perform a formal curve-fitting process to determine the parameters for the artificial waveforms. Instead, we matched them by eye to the grand average difference waveforms using the *Create Artificial ERP Waveform* tool in ERPLAB Toolbox. The parameters used for the simulated version of each component are shown in Table 3.

**Table 3.** Parameters used to create the artificial waveforms.

| Component | N170 | MMN | N2pc | P3 | N400 | LRP | ERN |
| --- | --- | --- | --- | --- | --- | --- | --- |
| Function | Gaussian | Gaussian | Gaussian | Ex-Gaussian | Ex-Gaussian | Gaussian | Ex-Gaussian |
| Peak Amplitude (μV) | -4.60 | -2.82 | -1.62 | 8.60 | -9.65 | -3.20 | -12.50 |
| Gaussian Mean (ms) | 129 | 190 | 257 | 310 | 280 | -47 | 76 |
| Gaussian SD (ms) | 14 | 33 | 28 | 58 | 65 | 36 | 26 |
| Exponential Tau (ms) | 0 | 0 | 0 | 2000 | 1400 | 0 | -300 |

We used these simulated ERP components to quantify the size of the artifactual opposite-polarity peaks produced by a given filter. This was calculated as the artifactual peak percentage, defined as 100 times the amplitude at the time of the largest artifactual peak divided by the amplitude at the time of the true peak (with the amplitudes obtained from the filtered waveform).

## 3. Results

Each of the following sections provides the details of the filter optimization process for each of the seven ERP components. These sections are written so that readers can focus on the section for a single component rather than reading the sections for all seven components. However, we recommend that all readers start with the section for the N170 component, which provides a more detailed description of how the optimal filter settings are obtained. The sections are ordered according to ascending latency for the stimulus-locked components (N170, MMN, N2pc, P3, N400), followed by the two response-locked components (LRP and ERN).

The optimal filter settings resulting from this process are listed for all seven components in Table 1. Note that these filter settings are somewhat more aggressive than the generic bandpass of 0.1–30 Hz that we have recommended previously for cognitive and affective research (Luck, 2014) and should lead to improved statistical power. This shows the value of the present quantitative approach for identifying optimal filter parameters.

### 3.1. The N170 component

As illustrated in Figure 1 a, the stimuli in the N170 paradigm consisted of a randomized sequence of faces, cars, scrambled faces, and scrambled cars (see Kappenman et al. (2021) for details). Participants were instructed to press one button for intact stimuli (faces or cars) and a different button for scrambled stimuli (scrambled faces or scrambled cars). Here, we focus solely on the ERPs elicited by faces and cars. Figure 1 b shows the grand average ERPs elicited by the faces and the cars, and Figure 1 c shows the faces-minus-cars difference wave overlaid with the artificial N170 waveform.

**Figure 1:**
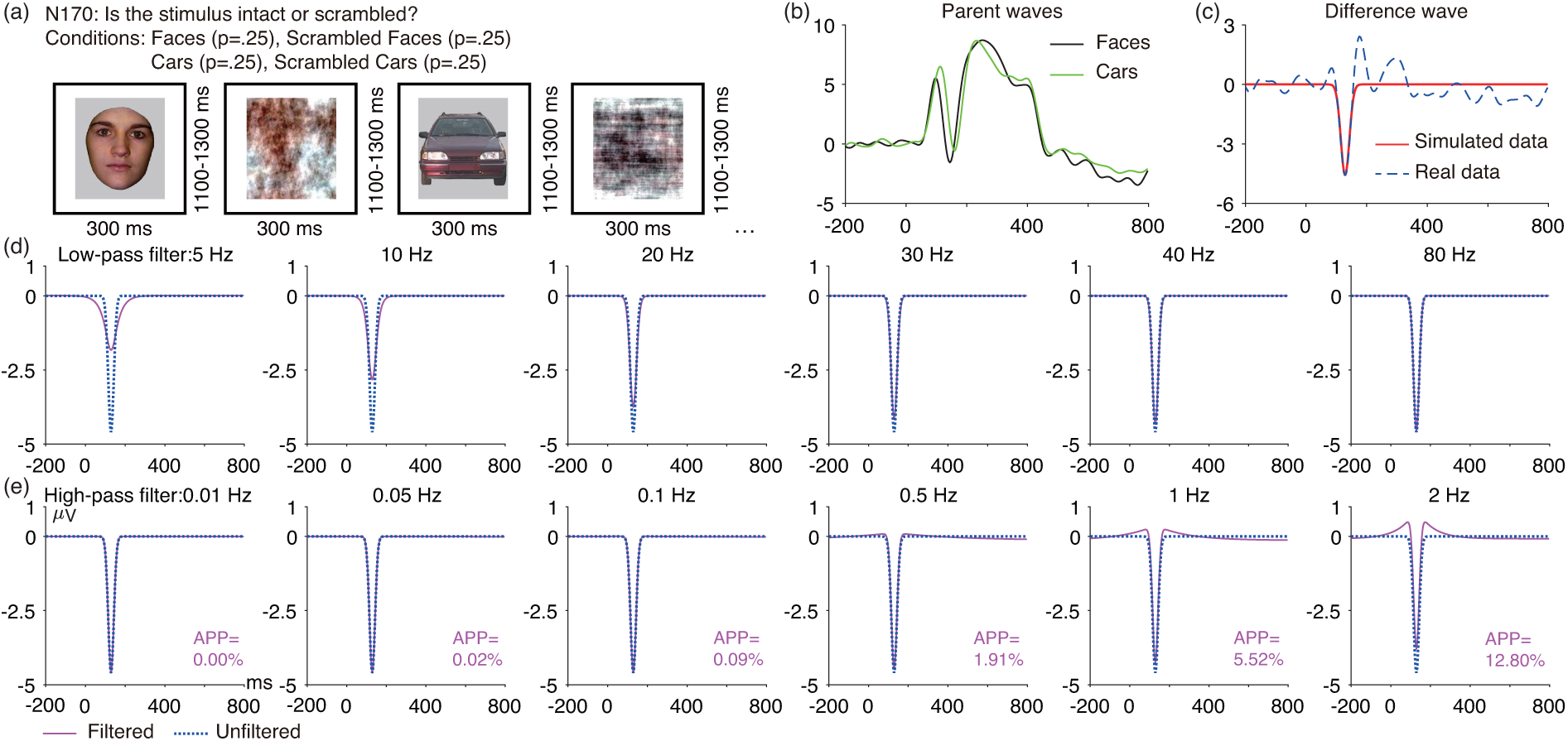
(a) N170 face perception paradigm. Only the data from the face and car trials were used in the present analyses. (b) Grand average ERP waveforms from the PO8 electrode site for the face and car trials. (c) Grand average face-minus-car difference wave at PO8, along with the simulated N170 difference wave (Gaussian function, mean = 129 ms, SD = 14 ms, peak amplitude = -4.6 µV). (d) Artificial waveform overlaid with the low-pass filtered version of that waveform for several different filter cutoffs. (e) Artificial waveform overlaid with the high-pass filtered version of that waveform for several different filter cutoffs. The number next to each high-pass filtered waveform is the artifactual peak percentage (APP). Note that the simulated waveforms were preceded and followed by 1000 ms of zero values to avoid edge artifacts. All the filters used here were noncausal Butterworth filters with a slope of 12 dB/octave, and cutoff frequencies indicate the half-amplitude point. The figure was adopted from Kappenman et al. (2021).

#### 3.1.1. N170 waveform distortion

Figure 1 d shows the effects of several low-pass filters on the simulated N170 waveform. As the low-pass cutoff frequency was reduced, the amplitude of the N170 was reduced, and the N170 became broader. With a 5 Hz cutoff frequency, the amplitude was reduced by more than 50%. Note that all cutoff frequencies listed in this paper indicate the half-amplitude point in the frequency response function, and the results would be different for half-power cutoffs at these frequencies. In addition, all filters examined here have a roll-off of 12 dB/octave, and the filter distortion would be greater for steeper roll-offs.

Figure 1 e shows the effects of several high-pass filters on the simulated N170 waveform. Even with a 2 Hz cutoff, high-pass filtering produced very little amplitude reduction. However, higher cutoff frequencies led to clear artifactual peaks before and after the N170 peak. If these peaks were large enough to be statistically significant, they could lead to invalid conclusions (e.g., that P1 amplitude is greater for faces than for cars). However, this is unlikely to occur if the artifactual peak is less than 5% of the size of the true peak (Zhang et al., 2023). This 5% threshold for artifactual peak percentage was exceeded for the 1 Hz and 2 Hz cutoffs. Supplementary Figure S2 provides artifactual peak percentage values for a denser sampling of cutoff frequencies. The 5% threshold for the artifactual peak percentage was exceeded for high-pass cutoff frequencies that exceeded 0.9 Hz. Thus, 0.9 Hz (with a 12 dB/octave roll-off) is the highest high-pass cutoff frequency that we would recommend for datasets like the ERP CORE N170 experiment.

#### 3.1.2. Optimal filters for N170 mean amplitude and peak amplitude

Figure 2 shows the signal, noise, and signal-to-noise ratio for several combinations of low-pass and high-pass cutoffs. Supplementary Figure S2 provides these values for a denser sampling of cutoff frequencies.

**Figure 2:**
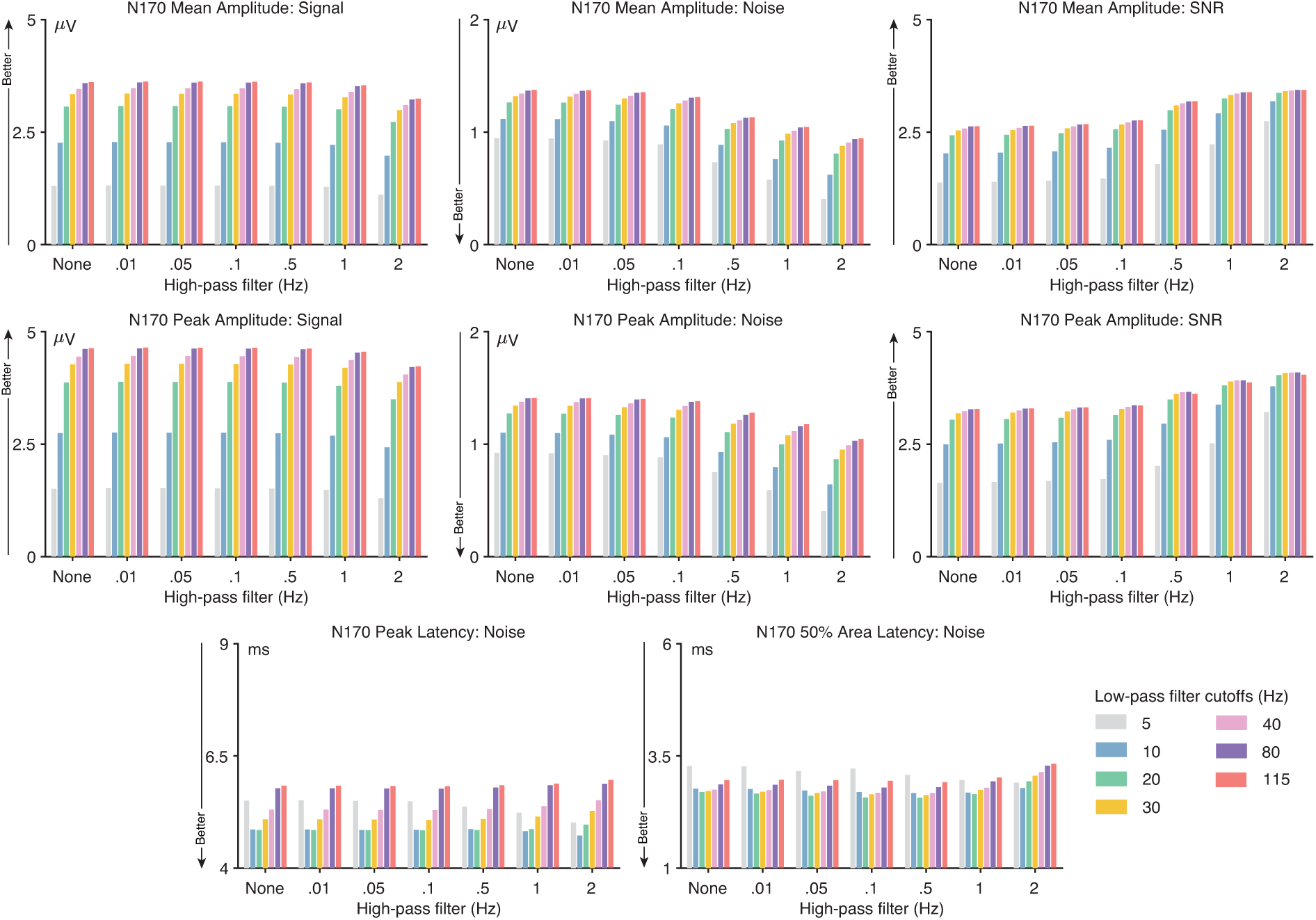
N170 data quality metrics for four different scoring methods and several combinations of high-pass filter cutoffs (0, 0.01, 0.05, 0.1, 0.5, 1, and 2 Hz) and low-pass filter cutoffs (5, 10, 20, 30, 40, 80, and 115 Hz). The signal was defined as the score (e.g., peak amplitude) obtained from the grand average ERP difference wave (faces minus cars). The noise was defined as the root mean square (RMS) of the single-participant standardized measurement error (SME) for that score. The signal-to-noise ratio (SNR) was computed as the signal divided by the noise. SNR is unitless. For latency scores, the signal is not consistently reduced by filtering, so only the RMS(SME) value is provided for the peak latency and 50% area latency scores.

These figures show that the N170 mean and peak amplitude scores were dramatically reduced when the low-pass cutoff was less than 20 Hz, but with little impact of the high-pass cutoff. These effects are consistent with the effects of filtering on the simulated N170 shown in Figure 1. The noise—quantified as RMS(SME)—was also progressively reduced as the low-pass cutoff frequency decreased and as the high-pass cutoff increased. Decreasing the low-pass cutoff frequency reduced the signal more than it reduced the noise, so decreasing the low-pass cutoff reduced the signal-to-noise ratio (*SNR_SME_*), especially for cutoffs of less than 30 Hz. This indicates that low-pass filtering is not helpful and can actually be harmful for N170 amplitude scores. For datasets like the ERP CORE N170 experiment, we therefore recommend no low-pass filtering, or a low-pass filter with a cutoff of 30 Hz or higher if desired to minimize the “fuzz” in the waveforms so that differences between groups or conditions are easier to visualize (see Table 1).

By contrast, increasing the high-pass cutoff frequency reduced the noise more than it reduced the signal, resulting in a progressive increase in the *SNR_SME_* value as the cutoff frequency increased. However, high-pass cutoffs of greater than 0.9 Hz produced artifactual peaks that exceeded our 5% threshold (see Figure S2), so we recommend a high-pass cutoff of 0.9 Hz.

The ERP CORE dataset contains relatively low levels of high-frequency noise. Low-pass filtering may become more valuable for datasets with greater high-frequency noise, especially for peak amplitude scores. Consistent with this, supplementary Figures S3 - S5 show that the *SNR_SME_*for peak amplitude was maximal for a low-pass filter cutoff of 30 Hz when high-frequency noise was added to the continuous EEG prior to filtering. Thus, we recommend a low-pass cutoff at 30 Hz for N170 peak amplitude scores when the EEG contains moderate levels of high-frequency noise. However, the *SNR_SME_* for mean amplitude was not improved by low-pass filtering in the noisier data. In addition, the increased noise did not change the recommended high-pass filter cutoffs for either mean amplitude or peak amplitude.

#### 3.1.3. Optimal filters for N170 peak latency and 50% area latency

The bottom row of Figure 2 shows how the noise level varied across filter settings for the N170 peak latency and 50% area latency scores. As noted in Section 1.1, filters can substantially reduce amplitude values but typically produce minimal bias for latency scores, so it is not necessary to compute the ratio of signal to noise when determining optimal filter settings for latency scores. Instead, the goal is to minimize the noise—quantified with the RMS(SME)— without creating problematic artifactual peaks.

For peak latency, the RMS(SME) was best with a low-pass cutoff of 10-20 Hz. With higher cutoff frequencies (e.g., 30 Hz), high-frequency noise in the data was not attenuated and added variability to the latency of the peak. With lower cutoff frequencies (e.g., 5 Hz), the amplitude of the N170 was substantially reduced by the low-pass filter, making it more difficult to reliably determine the peak latency. The same pattern was observed for the 50% area latency score, but with a smaller impact of the filtering. Consequently, we recommend a low-pass cutoff frequency of 10-20 Hz for N170 peak latency and 50% area latency scores^1^.

With a low-pass cutoff at 20 Hz, there was very little impact of the high-pass filter on the noise for either peak latency or 50% area latency, except that the noise started to rise when the cutoff exceeded 0.9 Hz. Because the noise was largely the same for high-pass cutoffs between 0 and 0.9 Hz (see supplementary Figure S2), any high-pass cutoff below 1 Hz would be justified for N170 latency scores.

### 3.2. The mismatch negativity (MMN)

As shown in Figure 3 a, the mismatch negativity (MMN) was elicited using a passive auditory oddball task. Participants watched a silent video while they were presented with a task-irrelevant sequence of standard tones (1000 Hz, 80 dB, p = .8) and deviant tones of a lower intensity (1000 Hz, 70 dB, p = .2). Each participant was presented with 800 standards and 200 deviants. Figure 3 b shows the grand average waveforms for standard and deviant stimuli at the FCz electrode site, and Figure 3 c shows the deviant-minus-standard difference wave overlaid with the simulated MMN.

**Figure 3:**
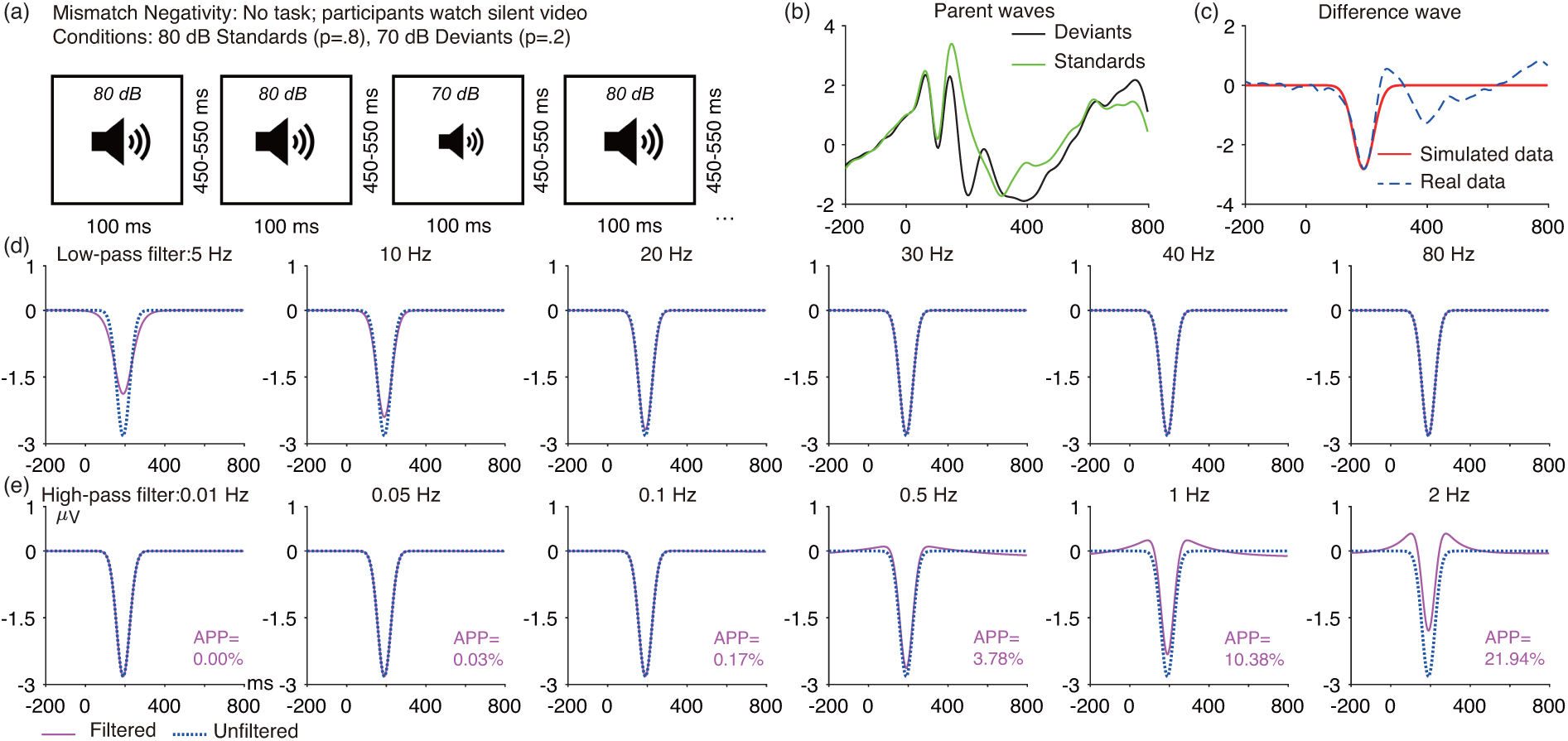
(a) MMN passive auditory oddball task. (b) Grand average ERP waveforms at FCz electrode site for deviant and standard trials. (c) Grand average deviant-minus-standard wave at FCz along with its simulated MMN difference wave (Gaussian function, mean = 190 ms, SD = 33 ms, peak amplitude = -2.82 *µV*). (d) Artificial waveform overlaid with the low-pass filtered version of that waveform for several different filter cutoffs. (e) Artificial waveform overlaid with the high-pass filtered version of that waveform for several different filter cutoffs. The number next to each high-pass filtered waveform is the artifactual peak percentage (APP). Note that the simulated waveforms were preceded and followed by 1000 ms of zero values to avoid edge artifacts. All the filters used here were noncausal Butterworth filters with a slope of 12 dB/octave, and cutoff frequencies indicate the half-amplitude poin

#### 3.2.1. MMN waveform distortion

Figure 3 shows the effects of several low-pass and high-pass filters on the simulated MMN waveform. For low-pass filters, the peak amplitude of the MMN was progressively reduced as the low-pass cutoff decreased, and the MMN became progressively broader. For high-pass filters, cutoffs above 0.5 Hz strongly distorted the MMN waveform, producing large opposite-polarity peaks before and after the MMN peak. Figure S6 provides results for a denser sampling of high-pass cutoff frequencies. The 5% threshold for the artifactual peak percentage was exceeded for high-pass cutoff frequencies greater than 0.5 Hz, so this is the highest high-pass cutoff frequency that we would recommend for datasets like the ERP CORE MMN experiment.

#### 3.2.2. Optimal filters for MMN mean amplitude and peak amplitude

As shown in Figure 4, the *SNR_SME_* for low-pass filters was largely constant when the cutoff frequency was 20 Hz and higher. For datasets like the ERP CORE MMN experiment, we therefore recommend no low-pass filtering, or a low-pass filter with a cutoff of 20 Hz or higher if desired to make it easier to visualize the ERP waveforms (see Table 1). However, for data with greater levels of high-frequency noise, a low-pass cutoff of 30 Hz yields the highest *SNR_SME_* for peak amplitude scores (see supplementary Figures S7 - S9); this cutoff is recommended when peak amplitude is being measured in moderately noisy data.

**Figure 4:**
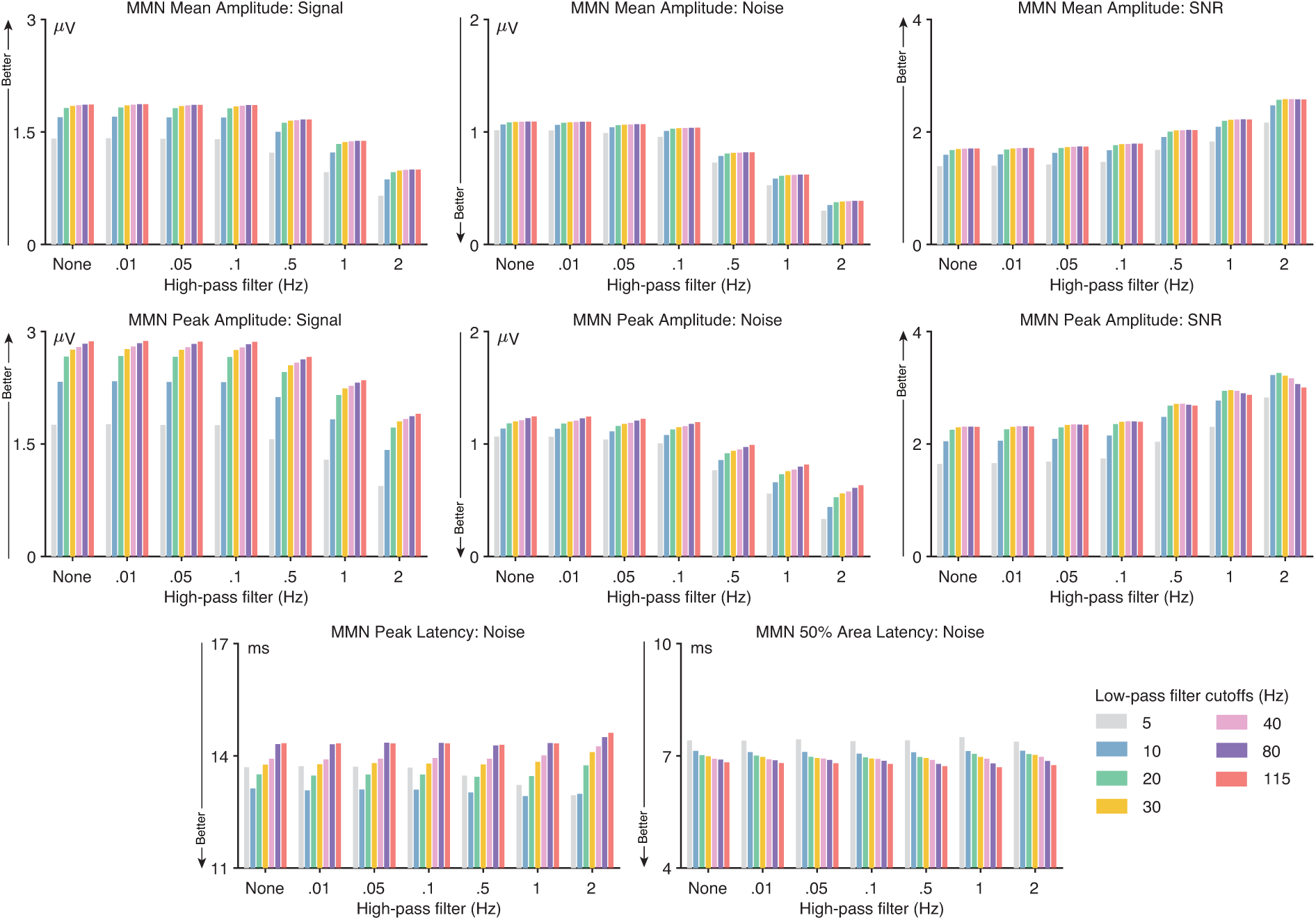
MMN data quality metrics for four different scoring methods and several combinations of high-pass filter cutoffs (0, 0.01, 0.05, 0.1, 0.5, 1, and 2 Hz) and low-pass filter cutoffs (5, 10, 20, 30, 40, 80, and 115 Hz). The signal was defined as the score (e.g., peak amplitude) obtained from the grand average ERP difference wave (deviants minus standards). The noise was defined as the root mean square (RMS) of the single-participant standardized measurement error (SME) for that score. The signal-to-noise ratio (SNR) was computed as the signal divided by the noise. SNR is unitless. For latency scores, the signal is not consistently reduced by filtering, so only the RMS(SME) value is provided for the peak latency and 50% area latency scores.

For high-pass filtering, the *SNR_SME_*increased as the high-pass cutoff increased, particularly at cutoffs of 1 Hz and higher. However, waveform distortion also increased as the high-pass cutoff increased. A high-pass cutoff of 0.5 Hz yielded the highest *SNR_SME_* while remaining below our 5% threshold for artifactual peak distortion, so this is our recommended high-pass cutoff frequency for MMN amplitude scores. This recommendation does not change for datasets with moderately greater noise levels (see supplementary Figures S7 - S9).

#### 3.2.3. Optimal filters for MMN peak latency and 50% area latency

The bottom row of Figure 4 shows how the RMS(SME) values varied across filter settings for the MMN peak latency and 50% area latency scores. For peak latency, the data quality was largely unaffected by high-pass filtering, but low-pass filtering had a substantial impact on the RMS(SME) values. The combination of a 0.5 Hz high-pass filter and a 10 Hz low-pass filter produced the lowest noise without exceeding our 5% threshold for artifactual peak percentage (see also supplementary Figure S6), so this is our recommendation for MMN peak latency. When white noise or pink noise was added to the data (supplementary Figures S8 and S9), decreasing the low-pass cutoff to 5 Hz slightly improved the noise level, so a 5 Hz cutoff may be more appropriate for data containing moderate amounts of broadband noise. However, a 5 Hz low-pass filter will spread the MMN considerably, making it appear to onset quite a bit earlier than the true onset time (see Figure 3), so a 10 Hz low-pass cutoff may be preferable (see Table 1).

The results for 50% area latency were quite different. High-pass filtering again had little effect, but low-pass filtering actually increased the noise levels slightly (see Figure 4 and supplementary Figure S6). We therefore recommend a high-pass cutoff of 0.5 Hz or lower and no low-pass filtering (or a low-pass filter with a cutoff of 20 Hz or higher to make it easier to visualize the ERP waveforms). These recommendations do not change for data with moderately greater noise levels (see supplementary Figures S7 - S9).

### 3.3. The N2pc component

As illustrated in Figure 5 a, the N2pc component was obtained using a simple visual search task. For a given block, either pink or blue was designated the target color, with the other color being the non-target. Each stimulus array contained one pink square, one blue square, and 22 black squares. The side containing the target color was unpredictable, but the target and nontarget color were always on opposite sites. For each array, participants pressed one of two buttons to indicate the location (top or bottom) of a gap in the attended-color square (while maintaining central fixation). Each participant completed a total of 320 trials (160 trials with the target on each side). Figure 5 b shows the grand average ERPs at the PO7/PO8 electrode site of the hemisphere contralateral or ipsilateral to the target. Figure 5 c shows the grand average contralateral-minus-ipsilateral difference wave overlaid with the simulated N2pc.

**Figure 5:**
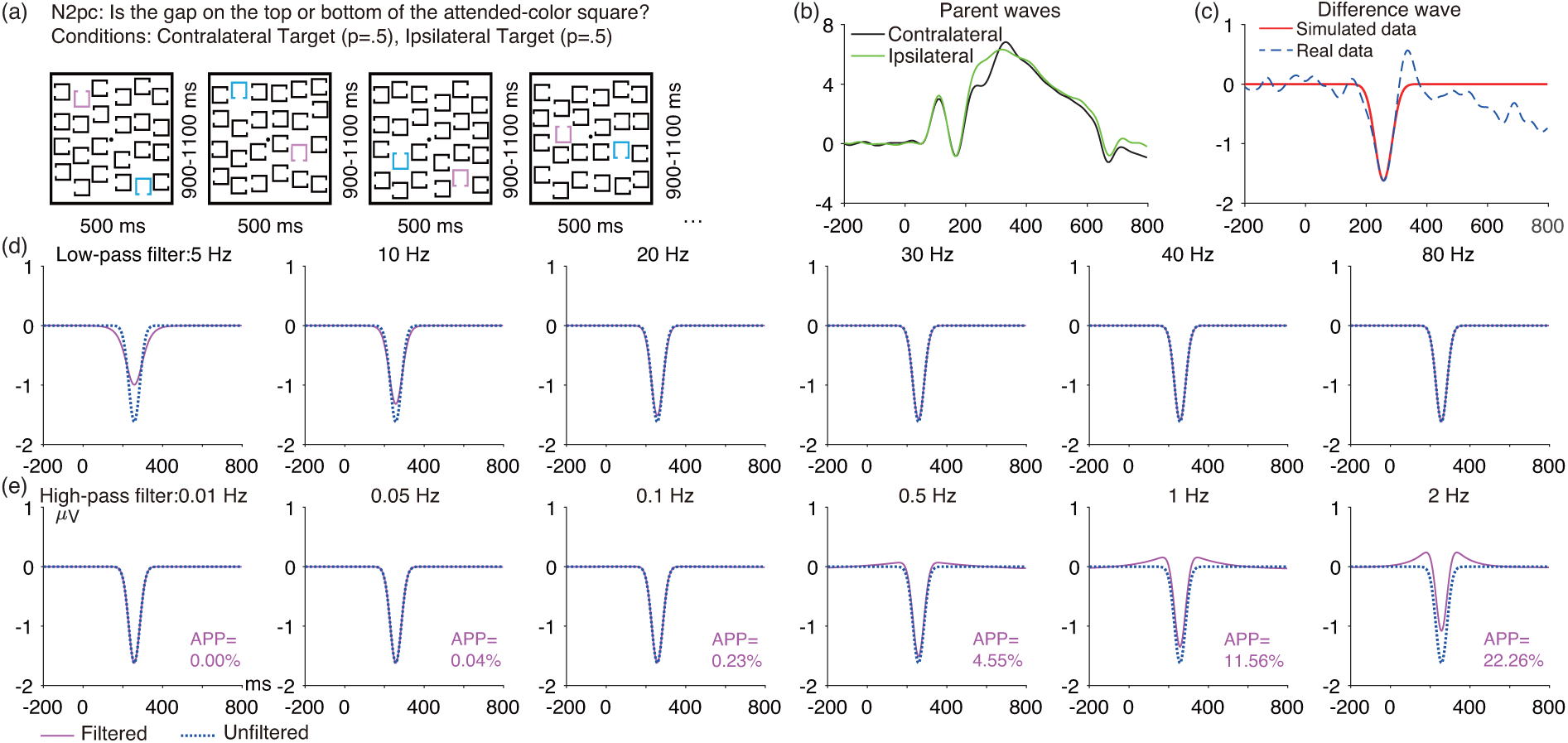
(a) N2pc simple visual search task. (b) Grand average waves from the PO7/PO8 electrode sites for contralateral and ipsilateral conditions. (c) Grand average contralateral-minus-ipsilateral difference wave at PO7/PO8, along with the simulated N2pc difference wave (Gaussian function, mean = 257 ms, SD = 28 ms, peak amplitude = -1.62 *µV*). (d) Artificial waveform overlaid with the low-pass filtered version of that waveform for several different filter cutoffs. (e) Artificial waveform overlaid with the high-pass filtered version of that waveform for several different filter cutoffs. The number next to each high-pass filtered waveform is the artifactual peak percentage (APP). Note that the simulated waveforms were preceded and followed by 1000 ms of zero values to avoid edge artifacts. All the filters used here were noncausal Butterworth filters with a slope of 12 dB/octave, and cutoff frequencies indicate the half-amplitude point.

#### 3.3.1. N2pc waveform distortion

Figure 5 shows the effects of several low-pass and high-pass filters on the simulated N2pc waveform. For low-pass filters, the N2pc became substantially smaller and broader when the cutoff was below 20 Hz. For high-pass filters, cutoffs above 0.5 Hz strongly distorted the N2pc waveform, producing large opposite-polarity peaks before and after the N2pc peak. Figure S10 provides results for a denser sampling of high-pass cutoff frequencies. The 5% threshold for the artifactual peak percentage was exceeded for high-pass cutoff frequencies greater than 0.5 Hz, so this is the highest high-pass cutoff frequency that we would recommend for datasets like the ERP CORE N2pc experiment.

#### 3.3.2. Optimal filters for N2pc mean amplitude and peak amplitude

As shown in Figure 6, low-pass filters had little impact on the N2pc except when the cutoffs were less than 20 Hz. Peak amplitude was more impacted by filtering, with reductions in *SNR_SME_* for frequencies less than 20 Hz and also for frequencies greater than 40 Hz. For datasets like the ERP CORE N2pc experiment, we therefore recommend a low-pass cutoff of *≥*20 Hz (or no low-pass filtering) for mean amplitude, and 20-40 Hz for peak amplitude. For datasets with more high-frequency noise, however, a low-pass cutoff of 20 Hz is optimal for peak amplitude scores (see supplementary Figures S11-S13; this assumes a high-pass cutoff of 0.5 Hz).

**Figure 6:**
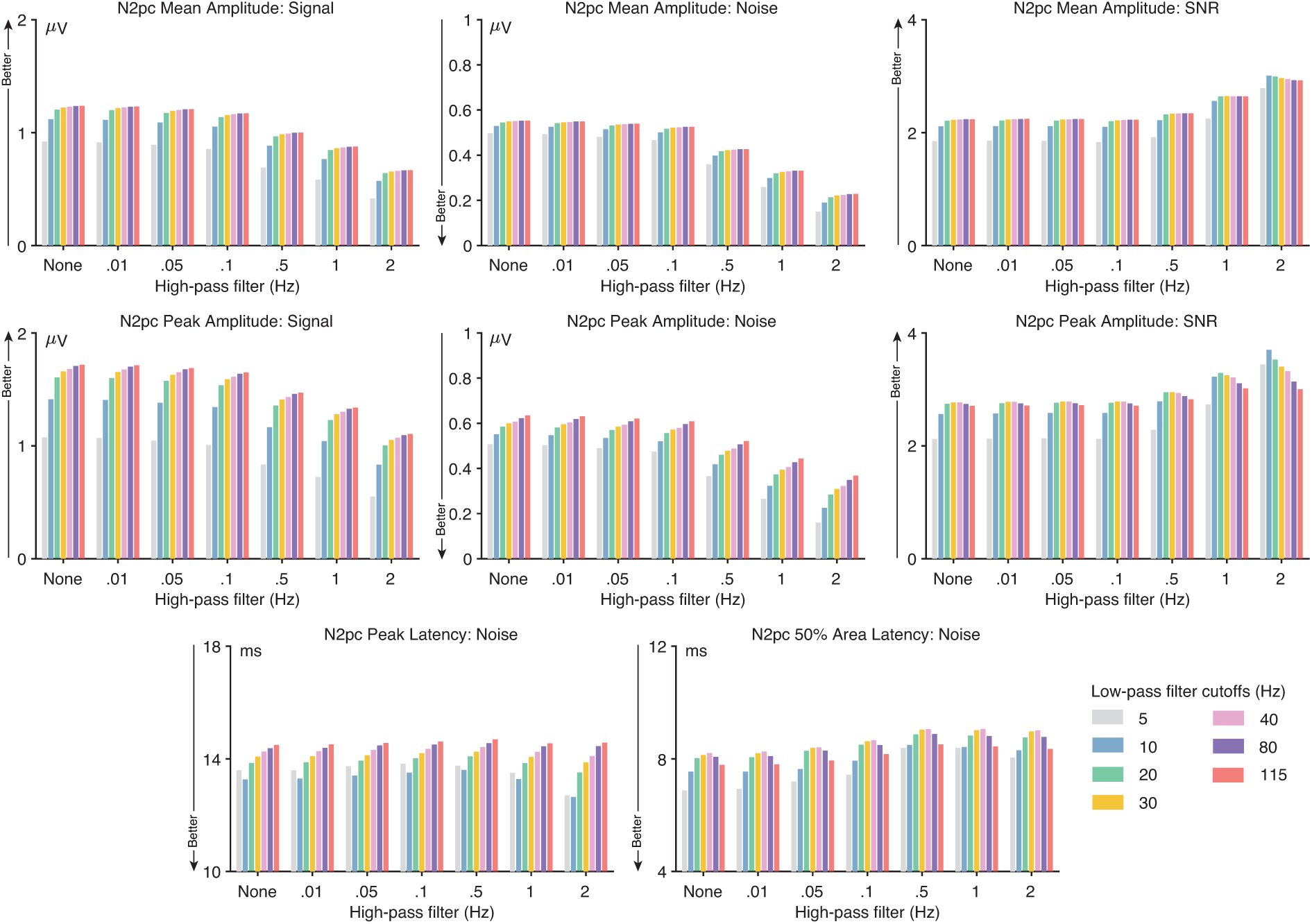
N2pc data quality metrics for four different scoring methods and several combinations of high-pass filter cutoffs (0, 0.01, 0.05, 0.1, 0.5, 1, and 2 Hz) and low-pass filter cutoffs (5, 10, 20, 30, 40, 80, and 115 Hz). The signal was defined as the score (e.g., peak amplitude) obtained from the grand average ERP difference wave (contralateral minus ipsilateral). The noise was defined as the root mean square (RMS) of the single-participant standardized measurement error (SME) for that score. The signal-to-noise ratio (SNR) was computed as the signal divided by the noise. SNR is unitless. For latency scores, the signal is not consistently reduced by filtering, so only the RMS(SME) value is provided for the peak latency and 50% area latency scores.

For high-pass filtering, the *SNR_SME_*increased as the high-pass cutoff increased, particularly at cutoffs of 0.5 Hz and higher (see Figure 6 and supplementary Figure S10). However, waveform distortion also increased as the high-pass cutoff increased. A high-pass cutoff of 0.5 Hz yielded the highest *SNR_SME_*while remaining below our 5% threshold for artifactual peak distortion, so this is our recommended cutoff. This recommendation does not change for datasets with moderately greater noise levels (see supplementary Figures S11 - S13).

#### 3.3.3. Optimal filters for N2pc peak latency and 50% area latency

The bottom row of Figure 6 shows how the RMS(SME) values varied across filter settings for N2pc peak latency and 50% area latency scores. For peak latency, low-pass filtering had a substantial impact on the RMS(SME) values but high-pass filtering had very little impact. The combination of a 10 Hz low-pass filter with either no high-pass filter or a .01 Hz high-pass cutoff produced the lowest noise without exceeding our 5% threshold for artifactual peak percentage (see also supplementary Figure S10), so this is our recommendation for N2pc peak latency. This recommendation does not change for datasets with moderately greater noise levels (see supplementary Figures S11 - S13).

For 50% area latency, a 5 Hz low-pass filter produced the lowest RMS(SME) values, combined with either no high-pass filter or a .01 Hz high-pass cutoff, so this is our recommendation. Note, however, that a 5 Hz low-pass filter will spread the N2pc considerably, making it appear to onset quite a bit earlier than the true onset time (see Figure 5), so some researchers may prefer a 10 Hz low-pass cutoff. These recommendations do not change for datasets with moderately greater noise levels (see supplementary Figures S11 - S13).

### 3.4. The P3 component

As illustrated in Figure 7 a, an active visual oddball task was used to elicit the P3 component. Participants were shown a randomized sequence of five letters (A, B, C, D, and E), with a probability of p = .2 for each letter. One of the five letters was designed as the target for a given block, and participants were instructed to press one button when the target was presented and a different button for any non-target. For example, if D was the target, participants pressed the target button for D and the non-target button for A, B, C, or E. Each participant received 40 target trials (referred to as rare trials) and 160 non-target trials (referred to as frequent trials). Figure 7 b shows the grand average ERP waveforms at the Pz electrode site for the rare and frequent conditions, and Figure 7 c shows the rare-minus-frequent difference wave overlaid with the simulated P3.

**Figure 7:**
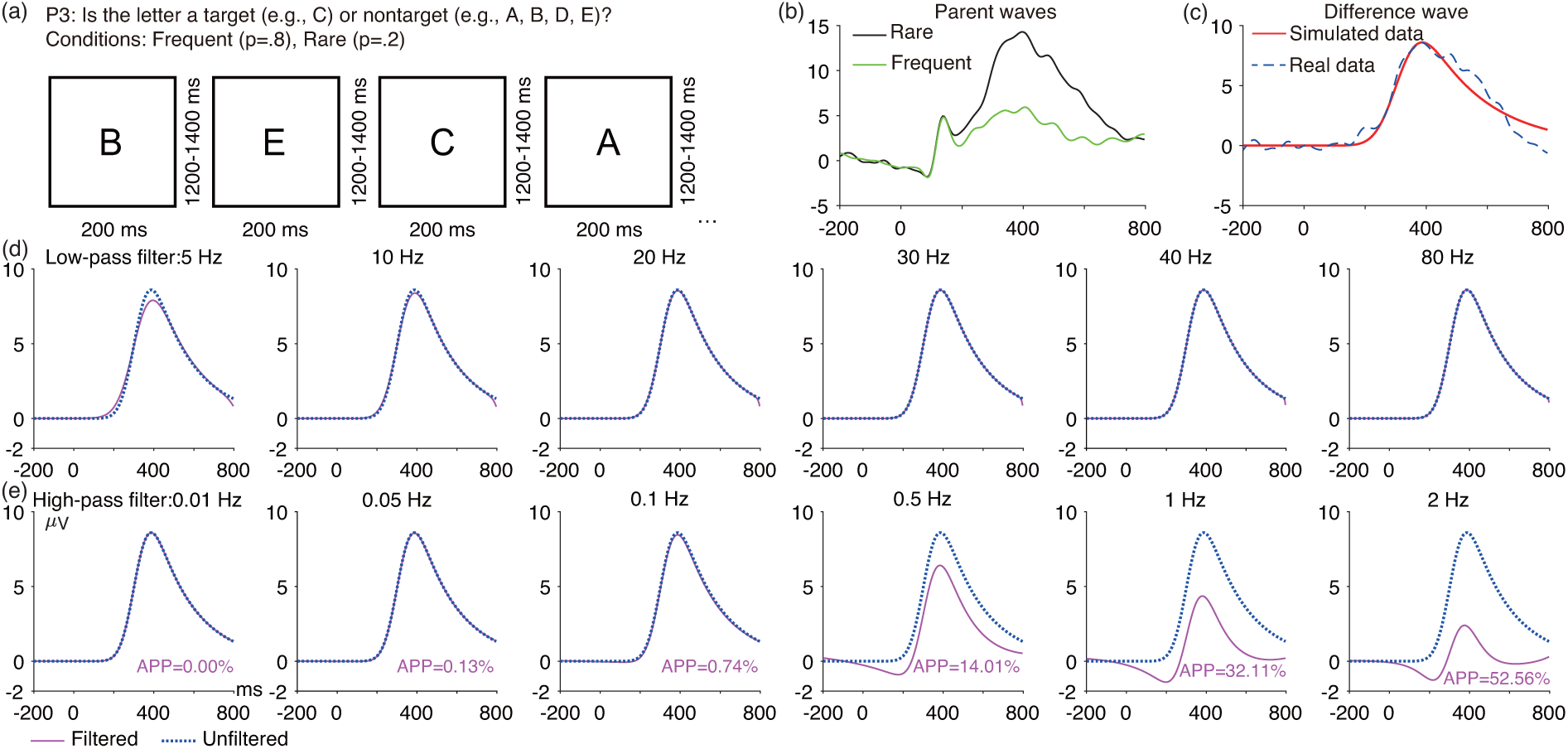
(a) P3 active visual oddball paradigm. (b) Grand average ERP waveforms from the Pz electrode site for the frequent and rare trials. (c) Grand average rare-minus-frequent difference wave at Pz, along with the simulated P3 difference wave (Ex-Gaussian function, mean = 310 ms, SD = 58 ms, *λ* = 2000 ms, peak amplitude = 8.6 *µV*). (d) Artificial waveform overlaid with the low-pass filtered version of that waveform for several different filter cutoffs. (e) Artificial waveform overlaid with the high-pass filtered version of that waveform for several different filter cutoffs. The number next to each high-pass filtered waveform is the artifactual peak percentage (APP). Note that the artificial waveforms were preceded and followed by 1000 ms of zero values to avoid edge artifacts. All the filters used here were noncausal Butterworth filters with a slope of 12 dB/octave, and cutoff frequencies indicate the half-amplitude point.

#### 3.4.1. P3 waveform distortion

Figure 7 shows the effects of several low-pass and high-pass filters on the simulated P3 waveform. Low-pass filters produced minimal distortion of this waveform for cutoffs of 10 Hz or higher. However, high-pass filters with cutoffs of 0.5 Hz or higher created substantial distortion, with a large negative artifactual peak preceding the true P3 wave that could easily be mistaken for an N2 effect. Figure S14 provides results for a denser sampling of high-pass cutoff frequencies. The 5% threshold for the artifactual peak percentage was exceeded for high-pass cutoff frequencies greater than 0.2 Hz, so this is the highest high-pass cutoff frequency that we would recommend for datasets like the ERP CORE P3 experiment.

#### 3.4.2. Optimal filters for P3 mean amplitude and peak amplitude

As shown in Figure 8, low-pass filters had minimal impact on the *SNR_SME_* for mean and peak P3 amplitude scores (unless an inadvisably strong high-pass filter was also used). For datasets like the ERP CORE P3 experiment, we therefore recommend no low-pass filtering, or a low-pass filter with a cutoff of 10 Hz or higher if desired to make it easier to visualize the ERP waveforms. For datasets with more noise, there is a small benefit to applying a 10 Hz low-pass filter for peak amplitude scores (see supplementary Figures S15-S17; this assumes a high-pass cutoff of *<* 0.5 Hz).

**Figure 8:**
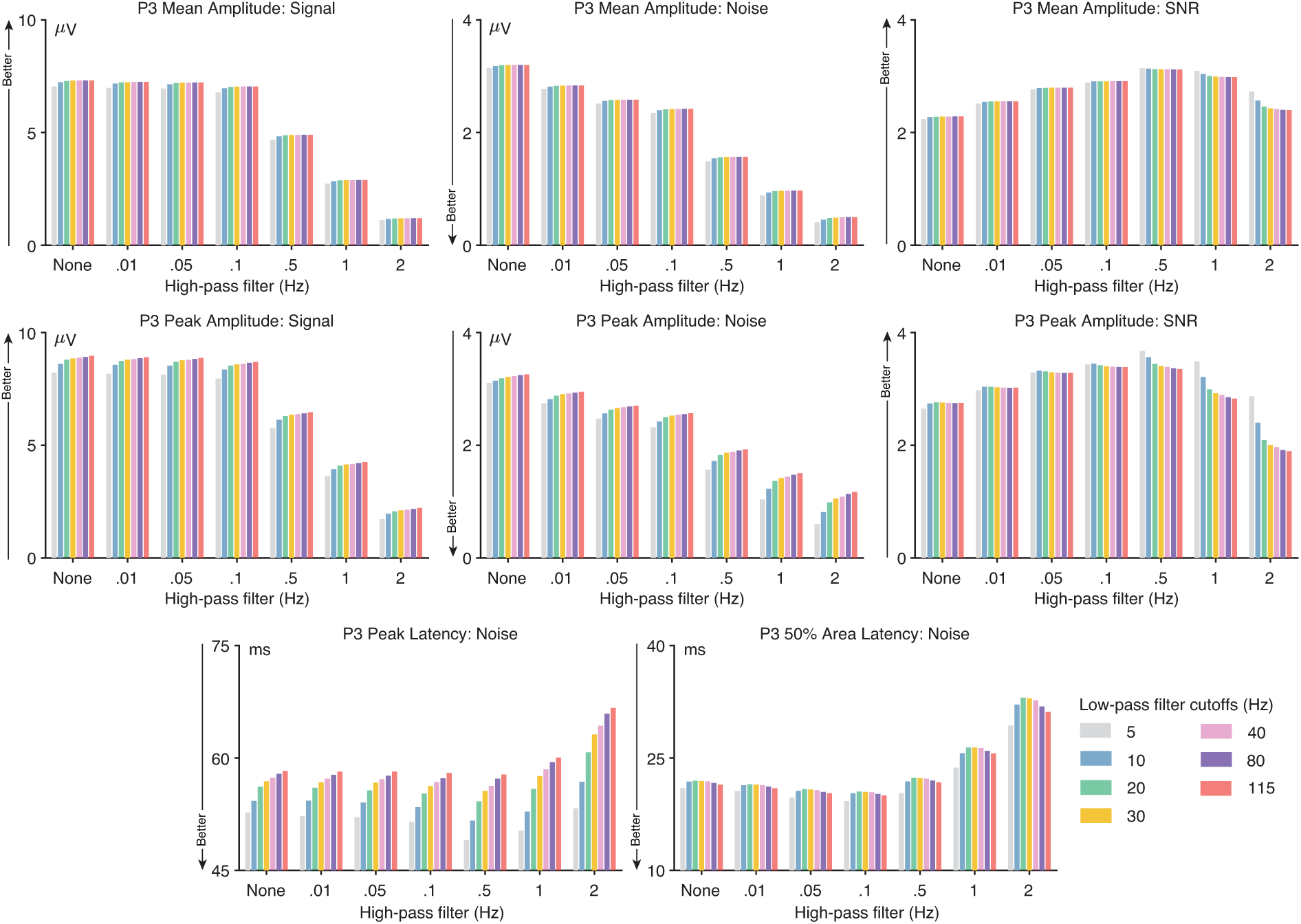
P3 data quality metrics for four different scoring methods and several combinations of high-pass filter cutoffs (0, 0.01, 0.05, 0.1, 0.5, 1, and 2 Hz) and low-pass filter cutoffs (5, 10, 20, 30, 40, 80, and 115 Hz). The signal was defined as the score (e.g., peak amplitude) obtained from the grand average ERP difference wave (frequent minus rare). The noise was defined as the root mean square (RMS) of the single-participant standardized measurement error (SME) for that score. The signal-to-noise ratio (SNR) was computed as the signal divided by the noise. SNR is unitless. For latency scores, the signal is not consistently reduced by filtering, so only the RMS(SME) value is provided for the peak latency and 50% area latency scores.

For high-pass filtering, the *SNR_SME_* increased as the high-pass cutoff increased up to 0.5 Hz (because the reduction in noise outweighed the small reduction in signal) but then decreased for higher cutoffs (because the large reduction in signal outweighed the reduction in noise; see Figure 8 and supplementary Figure S14). A high-pass cutoff of 0.2 Hz yielded the highest *SNR_SME_* while remaining below our 5% threshold for artifactual peak distortion, so this is our recommended cutoff. This recommendation does not change for datasets with moderately greater noise levels (see supplementary Figures S15 - S17).

#### 3.4.3. Optimal filters for P3 peak latency and 50% area latency

The bottom row of Figure 8 shows how the RMS(SME) values varied across filter settings for P3 peak latency and 50% area latency scores. For both peak latency and 50% area latency, a low-pass cutoff of 5 Hz produced the lowest noise, so that is our recommendation. However, this cutoff also produces a modest decrease in onset latency (Figure 7), so some researchers may prefer a low-pass cutoff of 10 Hz. A high-pass cutoff of 0.2 Hz yielded the lowest noise while remaining below our 5% threshold for artifactual peak distortion (see supplementary Figure S14), so this is our recommended high-pass cutoff. These recommendations do not change for datasets with moderately greater noise levels (see supplementary Figures S15 - S17).

### 3.5. The N400 component

As depicted in Figure 9 a, the N400 component was elicited using a word pair judgment paradigm. Each trial consisted of a red prime word followed by a green target word. Participants were required to report whether the green word was semantically related (p = .5) or unrelated (p = .5) to the preceding red word by pressing one of two buttons. A total of 120 trials were presented to each participant (60 trials for related word pairs and 60 trials for unrelated word pairs). Figure 9 b displays the grand average waveforms for the unrelated and related words at the CPz electrode site, and Figure 9 c shows the unrelated-minus-related difference wave overlaid with the simulated N400.

**Figure 9:**
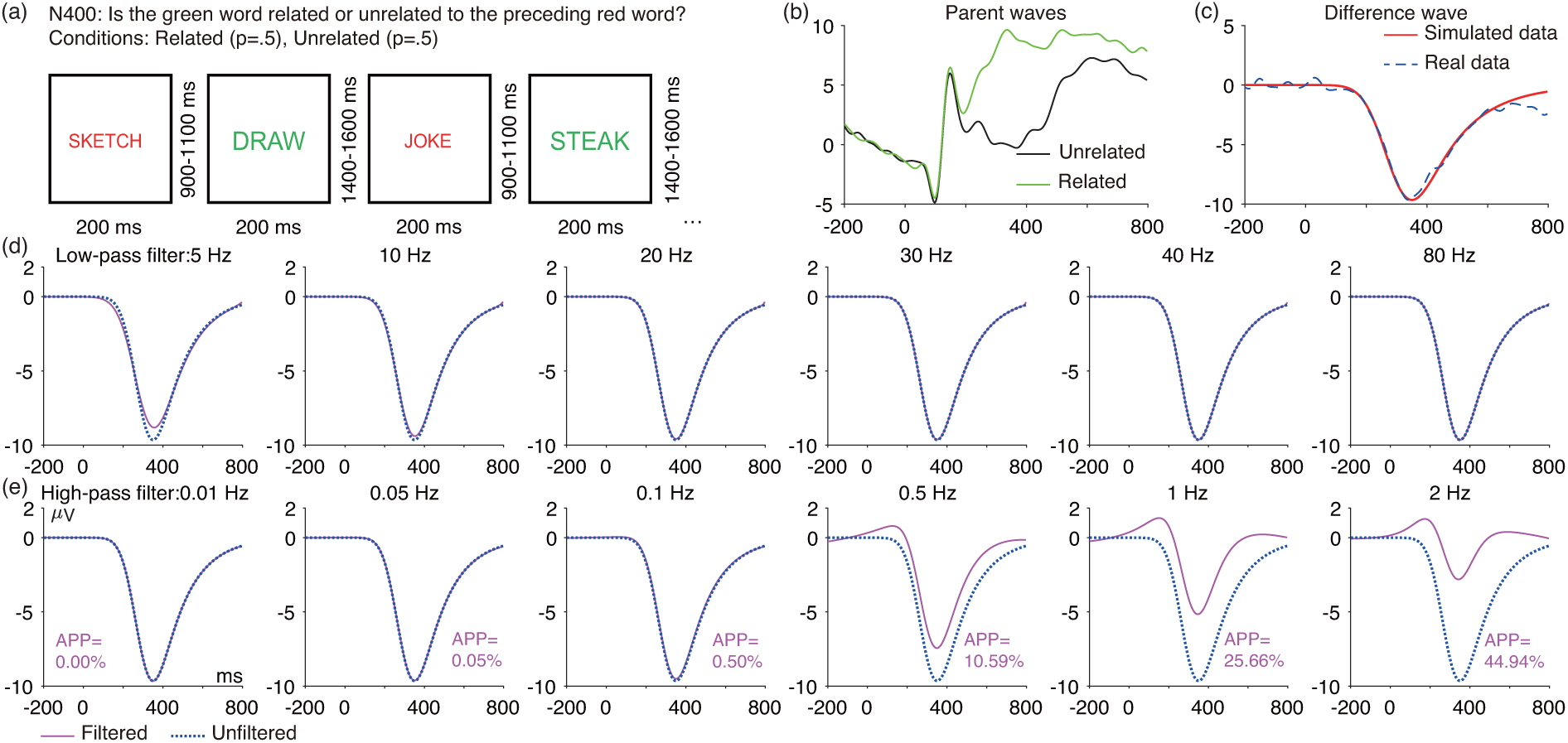
(a) N400 word pair judgment paradigm. (b) Grand average ERP waveforms at CPz electrode site for unrelated and related trials. (c) Grand average unrelated-minus-related difference wave along with its simulated N400 difference wave (Ex-Gaussian function, mean = 280 ms, SD = 65 ms, *λ* = 1400 ms, peak amplitude = -9.65 *µV*) (d) Artificial waveform overlaid with the low-pass filtered version of that waveform for several different filter cutoffs. (e) Artificial waveform overlaid with the high-pass filtered version of that waveform for several different filter cutoffs. The number next to each high-pass filtered waveform is the artifactual peak percentage (APP). Note that the artificial waveforms were preceded and followed by 1000 ms of zero values to avoid edge artifacts. All the filters used here were noncausal Butterworth filters with a slope of 12 dB/octave, and cutoff frequencies indicate the half-amplitude point.

#### 3.5.1. N400 waveform distortion

Figure 9 shows the effects of several low-pass and high-pass filters on the simulated N400 waveform. As was observed for the P3, low-pass filters produced minimal distortion of the N400 waveform as long as the cutoff was 10 Hz or higher. However, high-pass filters with cutoffs of 0.5 Hz or higher created substantial distortion, with a large positive artifactual peak preceding the true N400 wave that could easily be mistaken for a P2 effect. Figure S18 provides results for a denser sampling of high-pass cutoff frequencies. The 5% threshold for the artifactual peak percentage was exceeded for high-pass cutoff frequencies greater than 0.3 Hz, so this is the highest high-pass cutoff frequency that we would recommend for datasets like the ERP CORE N400 experiment.

#### 3.5.2. Optimal filters for N400 mean amplitude and peak amplitude

As shown in Figure 10, low-pass filters had minimal impact on the *SNR_SME_* for mean and peak N400 amplitude scores (unless an inadvisably strong high-pass filter was also used). For datasets like the ERP CORE N400 experiment, we therefore recommend no low-pass filtering, or a low-pass filter with a cutoff of 10 Hz or higher if desired to make it easier to visualize the ERP waveforms. For datasets with more noise, there is a small benefit to applying a 5 or 10 Hz low-pass filter (see supplementary Figures S19 - S21). For high-pass filtering, the *SNR_SME_* increased as the high-pass cutoff increased up to 0.2 Hz (presumably reflecting noise reduction) but then decreased for higher cutoffs (presumably because the reduction in signal outweighed the reduction in noise; see Figure 10 and supplementary Figure S18). A high-pass cutoff of 0.2 Hz was also below our 5% threshold for artifactual peak distortion, so this is our recommended high-pass cutoff. This recommendation does not change for datasets with moderately greater noise levels (see supplementary Figures S19 - S21).

**Figure 10:**
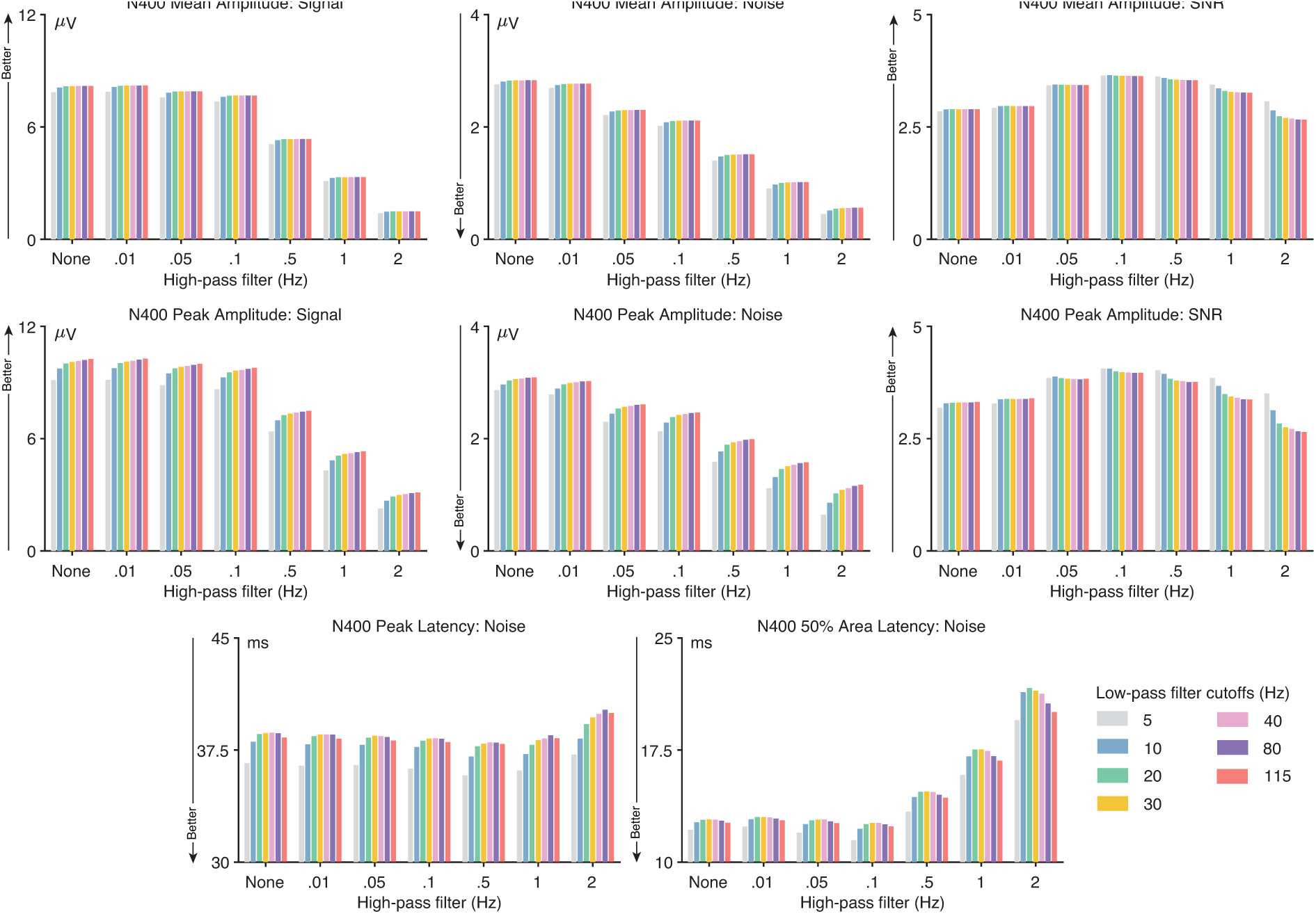
N400 data quality metrics for four different scoring methods and several combinations of high-pass filter cutoffs (0, 0.01, 0.05, 0.1, 0.5, 1, and 2 Hz) and low-pass filter cutoffs (5, 10, 20, 30, 40, 80, and 115 Hz). The signal was defined as the score (e.g., peak amplitude) obtained from the grand average ERP difference wave (unrelated minus related). The noise was defined as the root mean square (RMS) of the single-participant standardized measurement error (SME) for that score. The signal-to-noise ratio (SNR) was computed as the signal divided by the noise. SNR is unitless. For latency scores, the signal is not consistently reduced by filtering, so only the RMS(SME) value is provided for the peak latency and 50% area latency scores.

#### 3.5.3. Optimal filters for N400 peak latency and 50% area latency

The bottom row of Figure 10 shows how the RMS(SME) values varied across filter settings for the N400 peak latency and 50% area latency scores. For both peak latency and 50% area latency, a low-pass cutoff of 5 Hz produced the lowest noise, so that is our recommendation. However, this cutoff also produces a modest decrease in onset latency (see Figure 9), so some researchers may prefer a low-pass cutoff of 10 Hz. A high-pass cutoff of 0.2 Hz yielded the lowest noise while remaining below our 5% threshold for artifactual peak distortion (see supplementary Figure S18), so this is our recommended high-pass cutoff. These recommendations do not change for datasets with moderately greater noise levels (see supplementary S19 - S21).

### 3.6. The lateralized readiness potential (LRP)

As depicted in Figure 11 a, the LRP was elicited using a flankers paradigm. In this paradigm, each stimulus consisted of a central arrow surrounded by flanking arrows, which could either point in the same direction as the central arrow or in the opposite direction. On each trial, participants indicated whether the central arrow was pointing leftward (p = .5, 200 trials) or rightward (p = .5, 200 trials) by pressing a button with the corresponding hand. Figure 11 b shows the grand average ERPs at the C3/C4 electrode site of the hemisphere contralateral or ipsilateral to the responding hand (on trials with correct responses). Figure 11 c shows the grand average contralateral-minus-ipsilateral difference wave overlaid with the simulated LRP.

**Figure 11:**
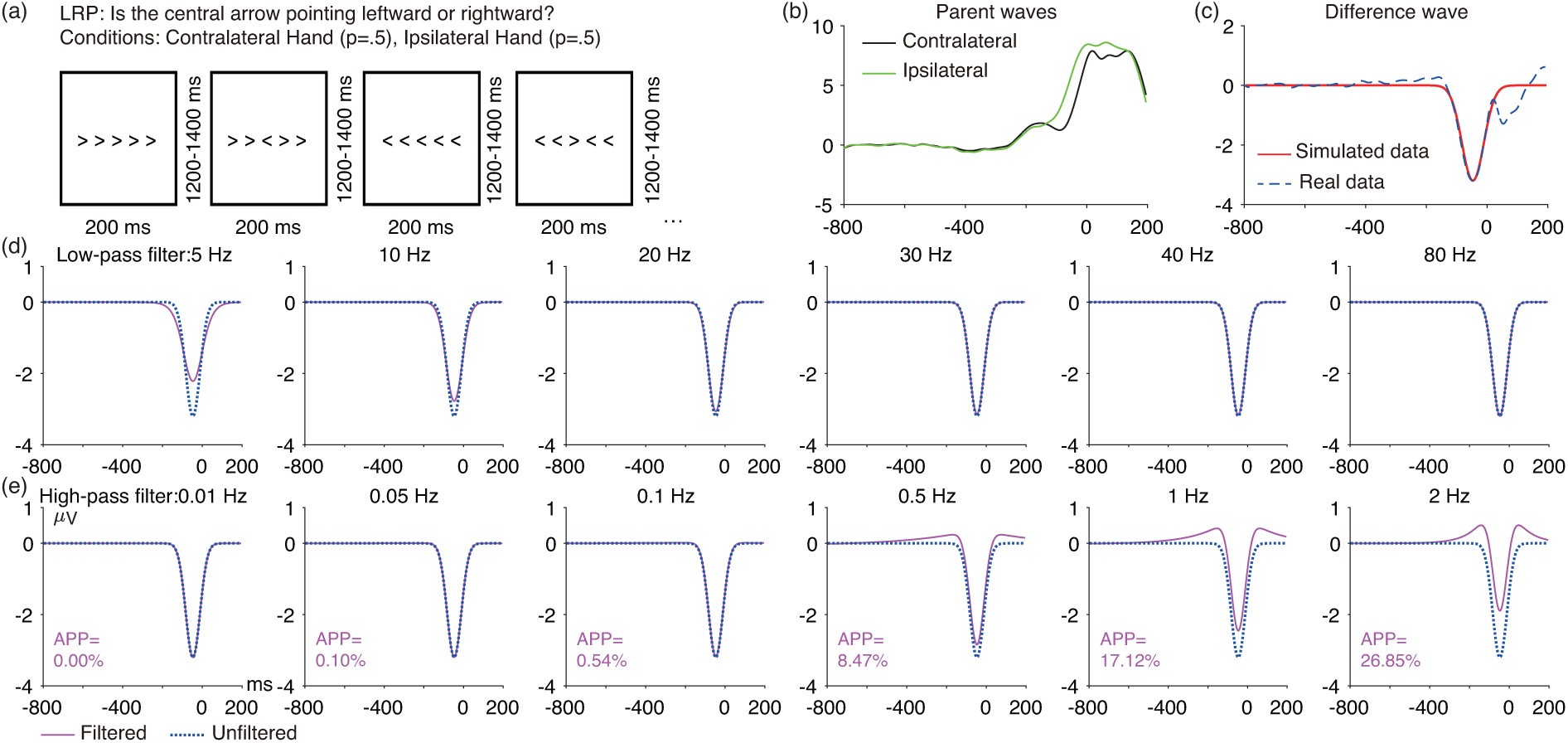
(a) LRP flankers task. (b) Grand average ERP waveforms from the C3/C4 electrode sites for the contralateral and ipsilateral trials. (c) Grand average contralateral-minus-ipsilateral difference wave at C3/C4, along with the simulated LRP difference wave (Gaussian function, mean = -47 ms, SD = 36 ms, peak amplitude = -3.2 *µV*). (d) Artificial waveform overlaid with the low-pass filtered version of that waveform for several different filter cutoffs. (e) Artificial waveform overlaid with the high-pass filtered version of that waveform for several different filter cutoffs. The number next to each high-pass filtered waveform is the artifactual peak percentage (APP). Note that the artificial waveforms were preceded and followed by 1000 ms of zero values to avoid edge artifacts. All the filters used here were noncausal Butterworth filters with a slope of 12 dB/octave, and cutoff frequencies indicate the half-amplitude point.

The LRP can be analyzed either time-locked to the stimulus or time-locked to the response. Here, we used response-locked averages. The LRP is typically broader in stimulus-locked averages than in response-locked averages, so different filter parameters may be appropriate for stimulus-locked averages. It would be straightforward for researchers to take the LRP scripts at https://osf.io/z3hfp/ and modify them to analyze stimulus-locked averages.

#### 3.6.1. LRP waveform distortion

Figure 11 shows the effects of several low-pass and high-pass filters on the simulated LRP waveform. For low-pass filters, the LRP became noticeably smaller and broader for cutoffs below 20 Hz. For high-pass filters, cutoffs above 0.1 Hz clearly distorted the LRP waveform, producing large opposite-polarity peaks before and after the LRP peak. Figure S22 provides results for a denser sampling of high-pass cutoff frequencies. The 5% threshold for the artifactual peak percentage was exceeded for high-pass cutoff frequencies greater than 0.3 Hz, so this is the highest high-pass cutoff frequency that we would recommend for datasets like the ERP CORE LRP experiment.

#### 3.6.2. Optimal filters for LRP mean amplitude and peak amplitude

As shown in Figure 12, low-pass filters had little impact on the *SNR_SME_* for LRP mean amplitude scores except that the *SNR_SME_*was reduced for cutoffs less than 30 Hz. For peak amplitude scores, the *SNR_SME_*was also reduced for low-pass cutoff frequencies greater than 30 Hz (except when combined with an inadvisably high high-pass cutoff).

**Figure 12:**
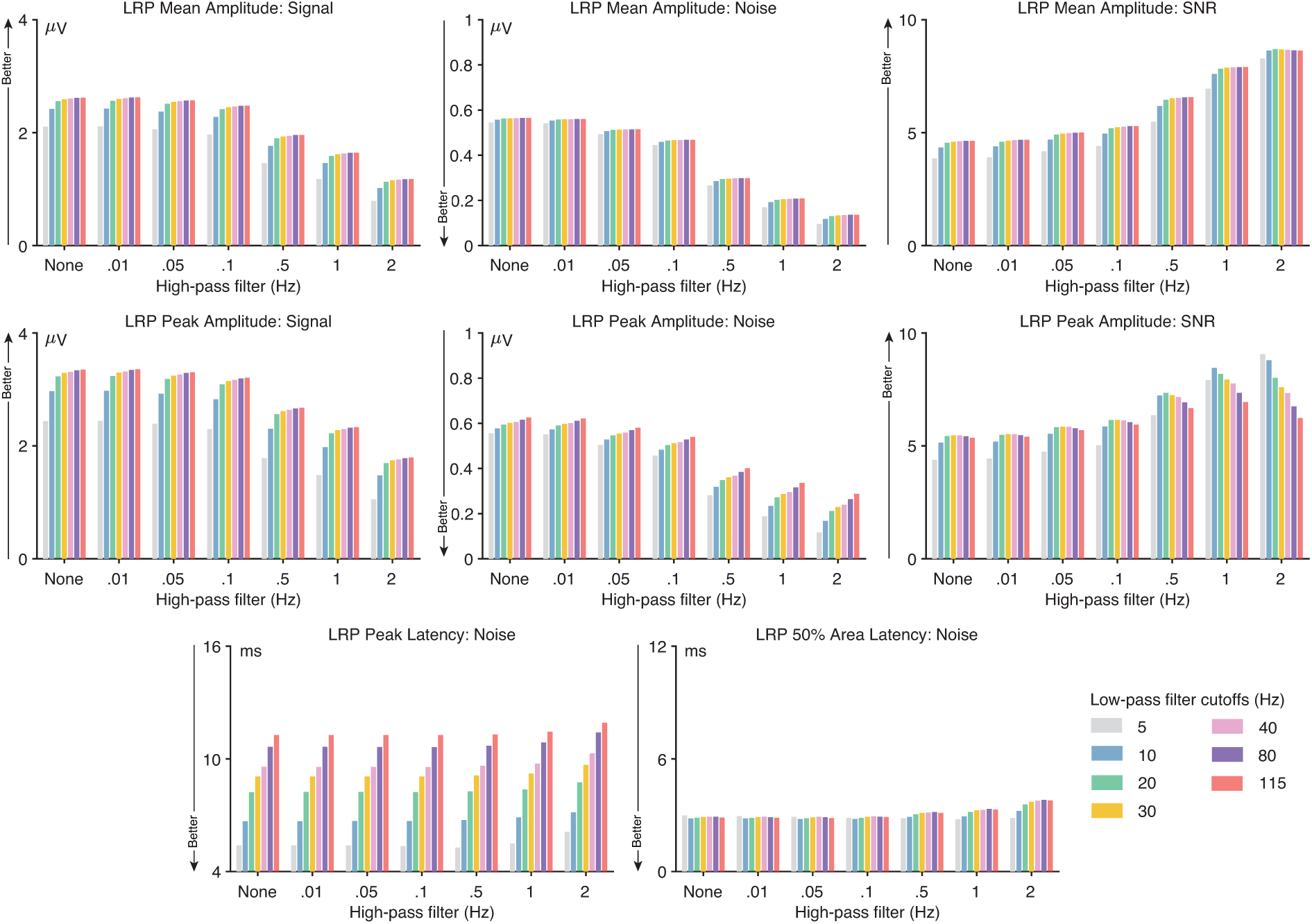
LRP data quality metrics for four different scoring methods and several combinations of high-pass filter cutoffs (0, 0.01, 0.05, 0.1, 0.5, 1, and 2 Hz) and low-pass filter cutoffs (5, 10, 20, 30, 40, 80, and 115 Hz). The signal was defined as the score (e.g., peak amplitude) obtained from the grand average ERP difference wave (contralateral minus ipsilateral). The noise was defined as the root mean square (RMS) of the single-participant standardized measurement error (SME) for that score. The signal-to-noise ratio (SNR) was computed as the signal divided by the noise. SNR is unitless. For latency scores, the signal is not consistently reduced by filtering, so only the RMS(SME) value is provided for the peak latency and 50% area latency scores.

For high-pass filtering, the *SNR_SME_*increased as the high-pass cutoff increased for both mean and peak amplitude (see also supplementary Figure S22). However, waveform distortion also increased as the high-pass cutoff increased. A high-pass cutoff of 0.3 Hz yielded the highest *SNR_SME_*while remaining below our 5% threshold for artifactual peak distortion. At this high-pass cutoff frequency, a low-pass cutoff of 30 Hz yielded the highest *SNR_SME_*for peak amplitude. Thus, for datasets like the ERP CORE LRP experiment, we recommend a high-pass cutoff of 0.3 Hz combined with a low-pass cutoff of 30 Hz for peak amplitude or *≥* 30 Hz for mean amplitude. For datasets with more high-frequency noise, decreasing the low-pass cutoff to 20 Hz would slightly improve the *SNR_SME_*for peak amplitude scores (see supplementary Figures S23 - S25).

#### 3.6.3. Optimal filters for LRP peak latency and 50% area latency

The bottom row of Figure 12 shows how the RMS(SME) values varied across filter settings for LRP peak latency and 50% area latency scores. For peak latency, low-pass filtering had a substantial impact on the RMS(SME) values but high-pass filtering had very little impact. The combination of a 5 Hz low-pass cutoff and a high-pass cutoff of 0.3 Hz produced the lowest noise without exceeding our 5% threshold for artifactual peak percentage (see also supplementary Figure S22), so this is our recommendation for LRP peak latency. However, a 5 Hz low-pass filter leads to a noticeably earlier LRP onset and later LRP offset, so researchers may want to use a 10 Hz high-pass cutoff instead. These recommendations do not change for datasets with moderately greater noise levels (see supplementary Figures S23 - S25).

For 50% area latency, the filter cutoffs had remarkably little effect on the RMS(SME) metric of noise (except when the high-pass cutoff exceeded the 5% artifactual peak threshold). The lowest noise value was obtained for a .3 Hz high-pass cutoff combined with a 5 Hz low-pass cutoff, so that is our recommendation. However, researchers may want to use a 10 Hz high-pass cutoff to minimize distortion of the LRP onset and offset times.

### 3.7. The error-related negativity (ERN)

As shown in Figure 13 a, the ERN was obtained from the same flankers paradigm used for the LRP (as described in Section 3.6). Whereas the LRP was defined as the difference between the contralateral and ipsilateral hemispheres (relative to the response hand), the ERN was defined as the difference between correct trials and error trials at a single midline electrode site (FCz). Each participant made correct responses on approximately 320-360 trials and incorrect responses on approximately 40-80 trials. Response-locked averages were used for the ERN. Figure 13 b presents the grand average ERP waveforms for trials with correct versus incorrect responses, and Figure 13 c shows the incorrect-minus-correct difference wave overlaid with the simulated ERN.

**Figure 13:**
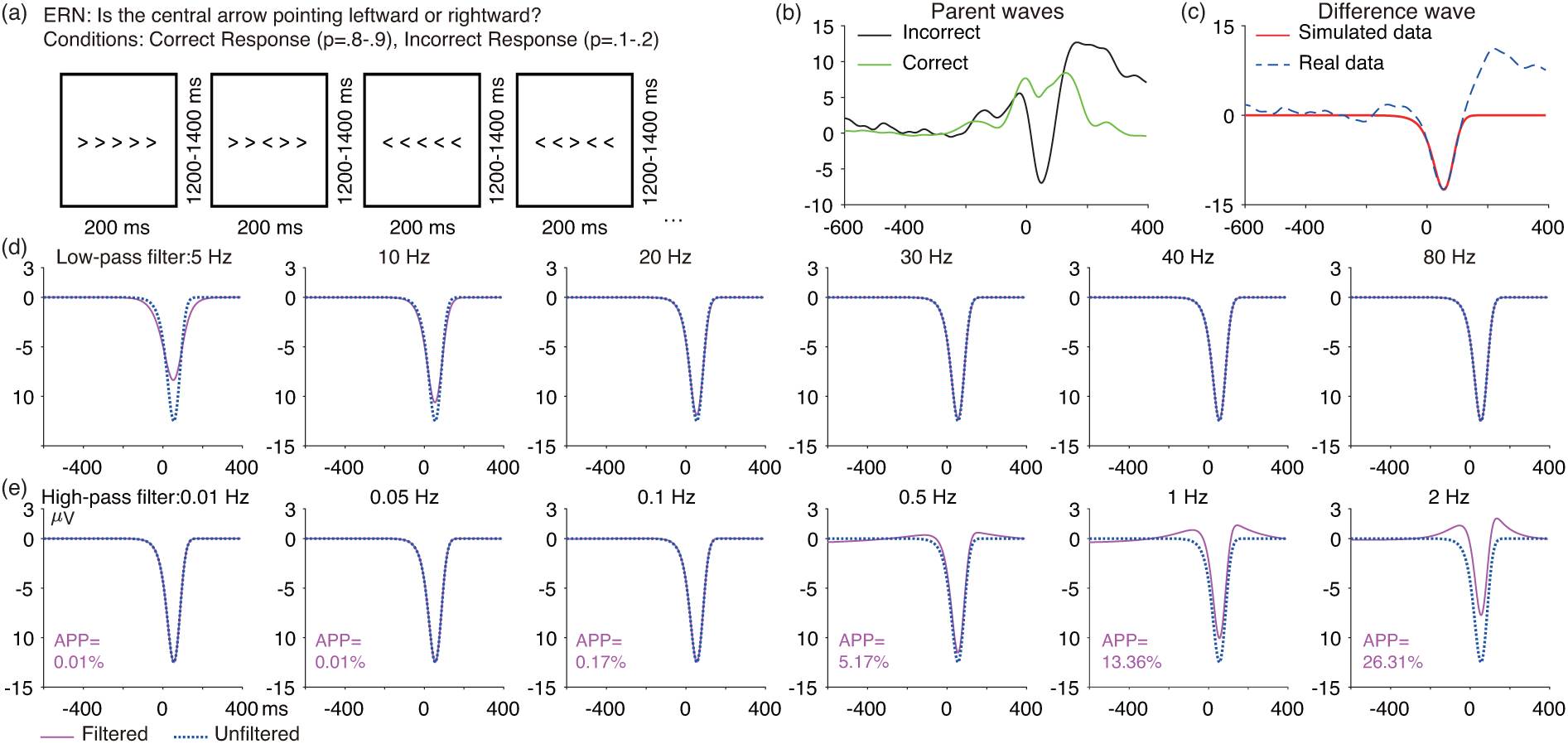
(a) ERN flankers task. (b) Grand average ERP waveforms from the FCz electrode site for correct and incorrect responses. (c) Grand average incorrect-minus-correct difference wave at FCz, along with its simulated ERP difference wave (Ex-Gaussian function, mean = 76 ms, SD = 26 ms, *λ* = -300 ms, peak amplitude = -12.5 *µV*). (d) Artificial waveform overlaid with the low-pass filtered version of that waveform for several different filter cutoffs. (e) Artificial waveform overlaid with the high-pass filtered version of that waveform for several different filter cutoffs. The number next to each high-pass filtered waveform is the artifactual peak percentage (APP). Note that the artificial waveforms were preceded and followed by 1000 ms of zero values to avoid edge artifacts. All the filters used here were noncausal Butterworth filters with a slope of 12 dB/octave, and cutoff frequencies indicate the half-amplitude point.

#### 3.7.1. ERN waveform distortion

Figure 13 shows the effects of several low-pass and high-pass filters on the simulated ERN waveform. For low-pass filters, the ERN became noticeably smaller and broader for cutoffs below 10 Hz. For high-pass filters, cutoffs of 0.5 Hz and higher clearly distorted the LRP waveform, producing distinct positive peaks before and after the ERN peak. Figure S26 provides results for a denser sampling of high-pass cutoff frequencies. The 5% threshold for the artifactual peak percentage was exceeded for high-pass cutoff frequencies greater than 0.4 Hz (when combined with most low-pass cutoffs), so this is the highest high-pass cutoff frequency that we would recommend for datasets like the ERP CORE ERN experiment.

#### 3.7.2. Optimal filters for ERN mean amplitude and peak amplitude

As shown in Figure 14, low-pass filters had relatively little impact on the *SNR_SME_* for ERN mean amplitude scores, except that the *SNR_SME_*was reduced for cutoffs less than 20 Hz. For peak amplitude scores, the *SNR_SME_* was maximal at 20 Hz (except when combined with an inadvisably high high-pass cutoff). For high-pass filtering, the *SNR_SME_*increased as the high-pass cutoff increased for both mean and peak amplitude (see also supplementary Figure S26). However, waveform distortion also increased as the high-pass cutoff increased. A high-pass cutoff of 0.4 Hz yielded the highest *SNR_SME_*for both mean and peak amplitude while remaining below our 5% threshold for artifactual peak distortion. Thus, for datasets like the ERP CORE LRP experiment, we recommend a high-pass cutoff of 0.4 Hz combined with a low-pass cutoff of 20 Hz for peak amplitude or *≥* 20 Hz for mean amplitude. These recommendations are also appropriate for data with modestly higher levels of noise (see supplementary Figures S27 - S29).

**Figure 14:**
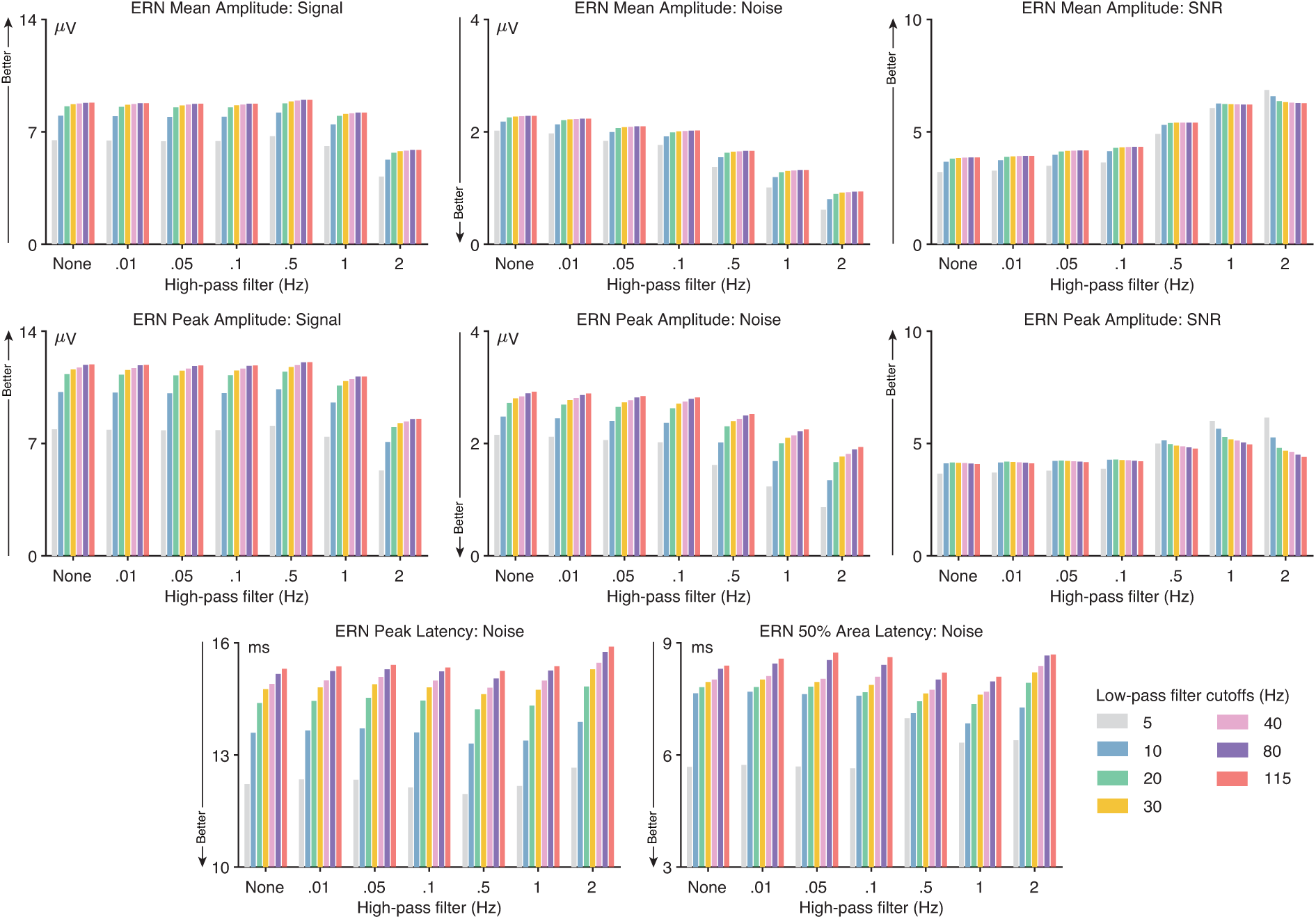
ERN data quality metrics for four different scoring methods and several combinations of high-pass filter cutoffs (0, 0.01, 0.05, 0.1, 0.5, 1, and 2 Hz) and low-pass filter cutoffs (5, 10, 20, 30, 40, 80, and 115 Hz). The signal was defined as the score (e.g., peak amplitude) obtained from the grand average ERP difference wave (correct minus incorrect). The noise was defined as the root mean square (RMS) of the single-participant standardized measurement error (SME) for that score. The signal-to-noise ratio (SNR) was computed as the signal divided by the noise. SNR is unitless. For latency scores, the signal is not consistently reduced by filtering, so only the RMS(SME) value is provided for the peak latency and 50% area latency scores.

#### 3.7.3. Optimal filters for ERN peak latency and 50% area latency

The bottom row of Figure 14 shows how the RMS(SME) values varied across filter settings for the ERN peak latency and 50% area latency scores. For both peak latency and 50% area latency, the lowest noise levels were produced by a low-pass cutoff of 5 Hz combined with no high-pass filtering (see also supplementary Figure S26), so this is our recommendation. However, a 5 Hz low-pass filter produces a noticeably earlier LRP onset and later LRP offset, so researchers may want to use a 10 Hz high-pass cutoff instead. These recommendations do not change for datasets with moderately greater noise levels (see supplementary Figures S27 - S29).

## 4. Discussion

The present study examined the impact of a broad range of low-pass and high-pass filter cutoffs on seven common ERP components recorded from a set of neurotypical young adults. We quantitatively assessed the effects of a broad range of filter cutoffs on waveform distortion and data quality (signal, noise, and signal-to-noise ratio). The observed effects led to recommendations for optimal filter cutoffs that maximize data quality while producing minimal waveform distortion. By repeating the analyses after adding artificial noise to the data, we were also able to provide recommendations for data with moderately greater noise levels (Table 1). These recommendations were based on objective, data-driven criteria, and similar studies that follow these recommendations should have improved data quality and therefore greater statistical power without inducing problematic waveform distortion. The data and the analysis scripts are available online at https://osf.io/z3hfp/, making it straightforward for researchers to repeat these analyses with different filters, different noise levels, or different thresholds for waveform distortion. In addition, the companion paper (Zhang et al., 2023) describes how the present approach can be applied to other datasets using tools available in version 9.2 and higher of ERPLAB Toolbox.

We would like to emphasize that the present results demonstrate that our previous recommendation (Luck, 2014) of a bandpass of 0.1–30 Hz was somewhat less than optimal, especially for relatively narrow components such as N170 and MMN. Some of the recommendations from other researchers were even farther from optimal, at least with respect to our criteria for optimality (e.g., 0.01-100 Hz for P3 and N400; Duncan et al. (2009)). This shows the value of using a quantitative, data-driven approach for selecting filter settings.

### 4.1. Effects of filter cutoffs on data quality and waveform distortion

In some cases, we found fairly large differences in noise levels and SNR for different filter settings. For example, the SNR for N170 peak amplitude was reduced by 50% when a 5 Hz low-pass filter was applied relative to when the cutoff was 30 Hz. Similarly, the SNR for P3 mean amplitude was 20% greater for the high-pass cutoff of 0.2 Hz recommended here than for the cutoff of 0.01 Hz recommended by Duncan et al. (2009) and Luck (2005). Moreover, increasing the high-pass cutoff from 0.2 Hz to 1 Hz for the P3 increased the size of the artifactual peak by a factor of almost 10 (see supplementary Figure S14). These results provide quantitative evidence that some filter cutoffs are much better than others.

On the other hand, small changes in cutoff frequencies typically produced only small changes in data quality and waveform distortion, so slight deviations from the recommendations shown in Table 1 would be expected to have only a minor impact. For example, our recommended high-pass filter cutoff for P3 mean amplitude was 0.2 Hz because the SNR was higher at 0.2 Hz than at 0.1 Hz and because cutoffs above 0.2 Hz led to an artifactual peak percentage that exceeded our 5% cutoff (see supplemental Figure S14). However, the SNR was only slightly better at 0.2 Hz than at 0.1 Hz (3.0 vs. 2.9), which is unlikely to be a meaningful difference in most studies. Moreover, it is possible that the SNR would be slightly better at 0.1 Hz than at 0.2 Hz in a different sample of participants. Similarly, a cutoff of 0.2 Hz might exceed our 5% threshold for the artifactual peak percentage in a study with a broader P3 waveform. However, the 0.2 Hz cutoff for P3 mean amplitude and the other values shown in Table 1 represent the current best guesses for optimal filter cutoffs for studies like the ERP CORE, and it would be sensible to use these cutoffs unless there is a good reason to use something else or until quantitative recommendations become available from a larger dataset.

### 4.2. Deviating from the present recommendations

One such reason for deviating from these recommendations would be the presence of very different noise levels. By adding noise to the ERP CORE data, we were able to demonstrate that more aggressive filtering of high frequencies (i.e., lower low-pass cutoffs) becomes optimal when there is more high-frequency or broadband noise in the data. However, the addition of moderately large noise did not impact our recommendations for high-pass filtering. This reflects the fact that our high-pass cutoff recommendations were mainly driven by the artifactual peak percentage, not by the SNR. For example, the best SNR for P3 mean amplitude was produced by a 0.5 Hz high-pass cutoff, but we recommended a 0.2 Hz cutoff to avoid exceeding the 5% threshold for artifactual peak percentage. For studies with very high levels of low-frequency noise and/or small numbers of trials, the increased SNR resulting from a 0.5 Hz high-pass cutoff might make it worthwhile to increase the threshold for artifactual peak percentage. In these cases, an artifactual peak of even 10% would be unlikely to be statistically significant and lead to incorrect conclusions.

Researchers might also choose to use the same filter cutoffs for different scoring methods, opting for simplicity over optimal data quality. For example, a researcher who is measuring both the mean amplitude and the peak latency of the ERN might use a bandpass of 0.4 – 10 Hz for both scores, even though a bandpass of 0.4–20 Hz is recommended for ERN mean amplitude. This would produce only a 2% drop in SNR (see supplementary Figure S26), which is unlikely to change the results of a single study. By contrast, if the optimal LRP peak latency bandpass of 0.3 – 5 Hz is also applied to LRP mean amplitude data, this would produce a 17% reduction in SNR relative to the optimal mean amplitude bandpass of 0.3 – 30 Hz (see supplementary Figure S22). This is more likely to be problematic.

In the end, decisions about filter cutoffs should reflect the scientific goals of a given study along with the nature of the data being filtered. The present approach to filter selection allows researchers to make these decisions in an informed manner.

### 4.3. General patterns

Several general patterns can be seen in the optimal filter values shown in Table 1. First, the optimal filter settings depend on the specific scoring method. Mean amplitude scores are largely insensitive to high-frequency noise, whereas high-frequency noise can distort peak amplitude and peak latency (seeClayson et al. (2013); Luck (2014)). Consequently, a low-pass filter is ordinarily unnecessary for mean amplitude, but it can improve data quality for peak amplitude and peak latency. For example, whereas low-pass filtering does not improve the SNR for N2pc mean amplitude, applying a 10 Hz low-pass filter reduces the noise level of N2pc peak latency scores by approximately 20% when the data are contaminated by substantial broadband noise (see supplementary Figure S12). However, low-pass filtering had much less effect on N2pc 50% area latency scores.

Our results also revealed that the optimal filter settings depend on the shape of the ERP component that is being analyzed. As a component becomes broader, it suffers less amplitude reduction from low-pass filtering but more amplitude reduction from high-pass filtering. In addition, high-pass filters produce larger artifactual peaks in broader components than in narrower components. Consequently, relatively high low-pass and high-pass cutoffs are optimal for narrower components (e.g., N170) and relatively low low-pass and high-pass cutoffs are optimal for broader components (e.g., P3). Researchers can use this finding to make informed guesses about optimal cutoffs for components that were not examined in the present study.

### 4.4. Limitations and future directions

The present study has some important limitations. First, our recommendations may not be applicable for recording setups that yield very different types or levels of noise. For example, data collected with dry electrodes often contain substantially more noise (Kam et al., 2019; Li et al., 2020; Shad et al., 2020). Similarly, our recommendations may not generalize to mobile experiments, for which a previous study has shown that higher high-pass filter cutoffs are beneficial (Klug & Gramann, 2021). However, it would be important for future research on optimal filter values for these systems to use an approach like we have used, which quantifies the data quality for the specific score of interest and also quantifies the amount of waveform distortion.

Another limitation is that the data used in the present study were obtained from highly cooperative college students, and the optimal filter settings may differ considerably in other populations, such as infants or individuals with clinical disorders. For example, in infant EEG recordings, higher high-pass filter cutoffs (e.g., 1 Hz) are currently viewed as appropriate (Gabard-Durnam et al., 2018; Debnath et al., 2020). However, such high cutoffs may induce problematic waveform distortion and may attenuate the signal more than the noise. Consequently, a careful quantitative analysis of a broad set of paradigms and scoring methods would be needed to determine the optimal filter settings for infants, specific clinical populations, and other groups that are likely to be quite different from the college students tested in the ERP CORE.

Finally, the present study assessed only four scoring methods, and did not consider onset latencies or mass univariate approaches. Moreover, it focused only on the Butterworth family of infinite impulse response filters, and it is possible that the results would be different for other filter families (e.g., finite impulse response filters). It would be valuable for future research to determine the optimal filter settings for other families of filters and other ERP scoring approaches.

## Funding Information

This study was made possible by grants R01MH087450 and R25MH080794 from the National Institute of Mental Health.

## Appendix A. Supplementary materials

**Figure S1:**
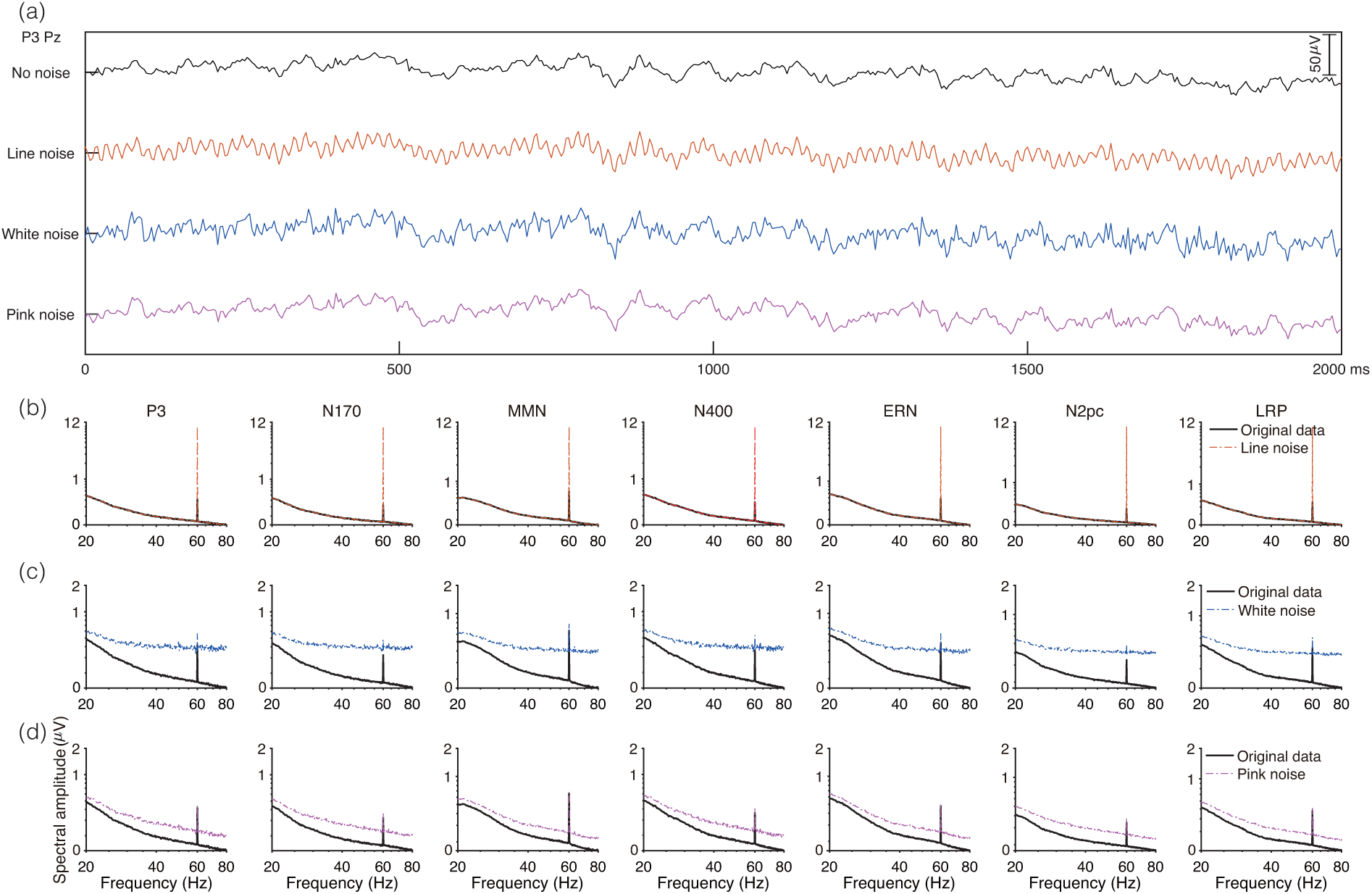
(a) Example of an EEG data segment before and after the addition of three different types of noise (60 Hz line noise, pink noise, and white noise). The data were obtained from the first 2000 ms of the P3 paradigm in Participant 1 at the Pz electrode site. (b, c, d) Grand average EEG amplitude density as a function of frequency for each component in the original data and in the data with added line noise (b), white noise (c), and pink noise (d). Amplitude density was calculated from each individual participant at the electrodes of interest for a given component (see Table 2) and then averaged across participants. Both the amplitude and frequency are on logarithmic scales. The frequency range shown here is limited to 20-80 Hz to make it easier to see the amplitudes at the frequencies where adding noise led to a substantial increase in amplitude. Full amplitude spectra for the original data are shown in Kappenman et al. (2021).

**Figure S2:**
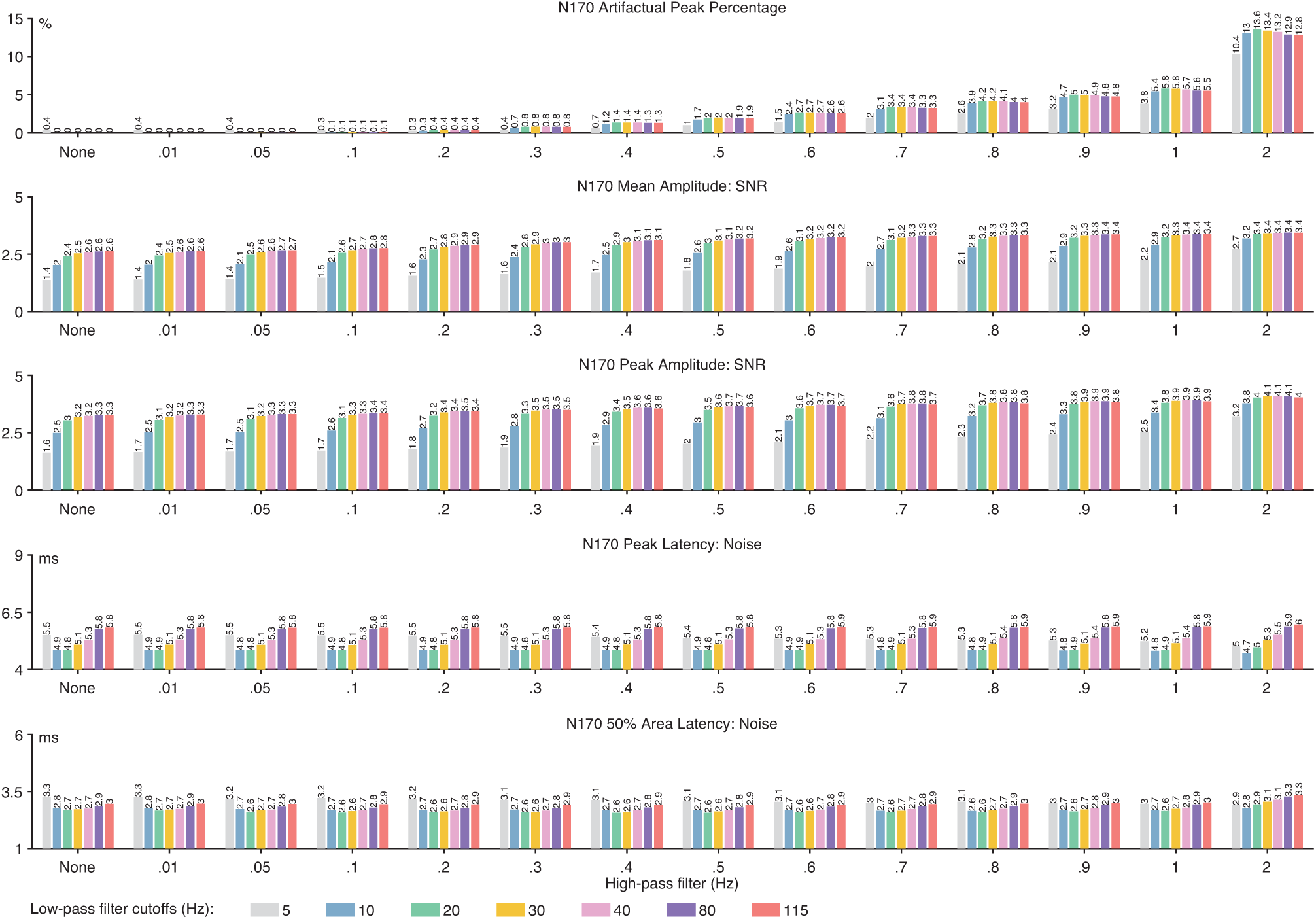
N170 artifactual peak percentage and data quality metrics. Data quality metrics are provided for each of the four scoring methods (mean amplitude, peak amplitude, peak latency and 50% area latency) and for each combination of high-pass filter cutoff frequency (0, 0.01, 0.05, 0.1, 0.2, 0.3, 0.4, 0.5, 0.6, 0.7, 0.8, 0.9, 1, and 2 Hz) and low-pass filter cutoff frequency (5, 10, 20, 30, 40, 80, and 115 Hz). The SNR values are unitless.

**Figure S3:**
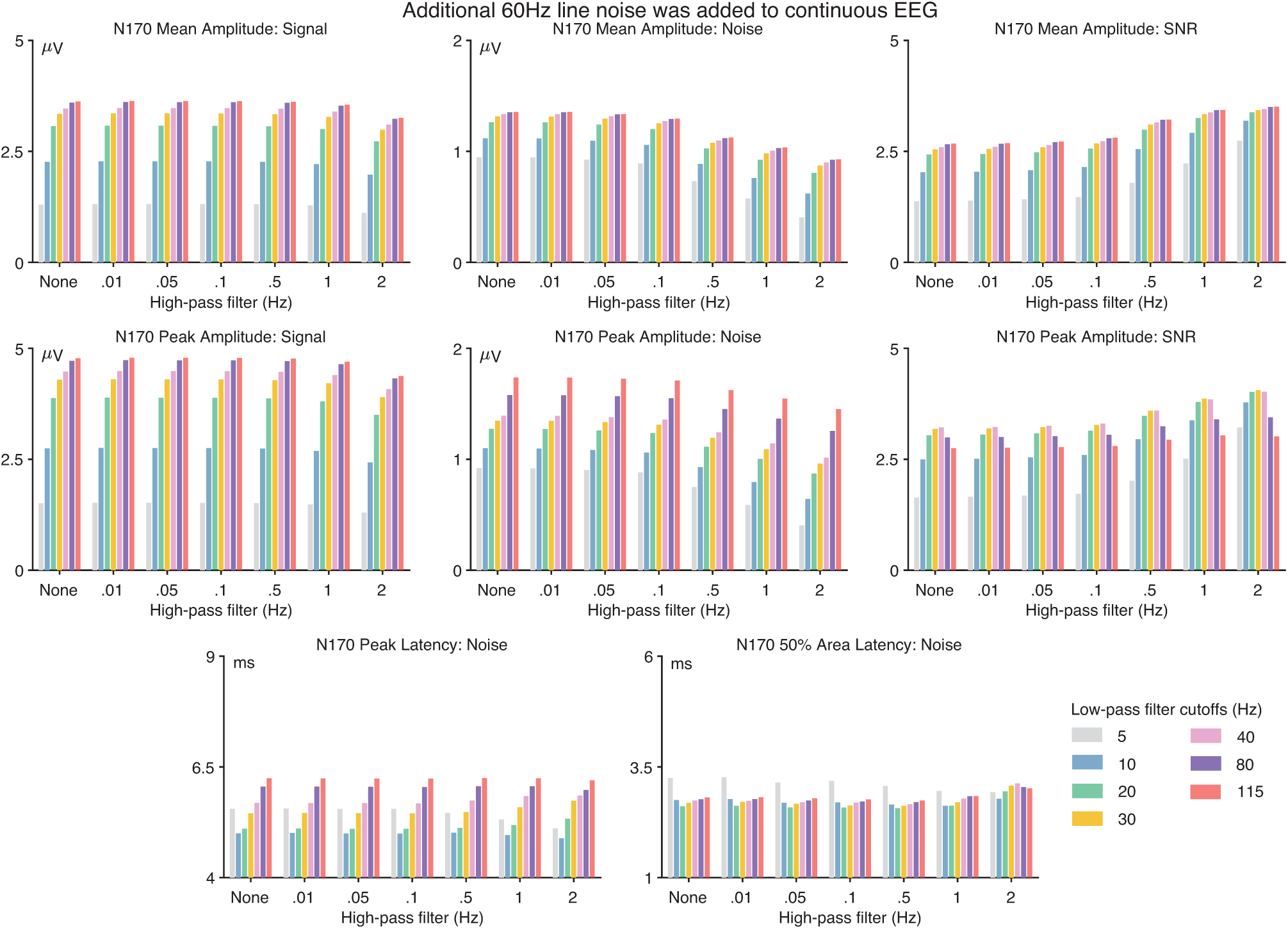
Data quality metrics for the N170 component when 60 Hz line noise (20 µV peak-to-peak amplitude) had been added to the continuous EEG prior to filtering. The format is identical to that of Figure 2 in the main document.

**Figure S4:**
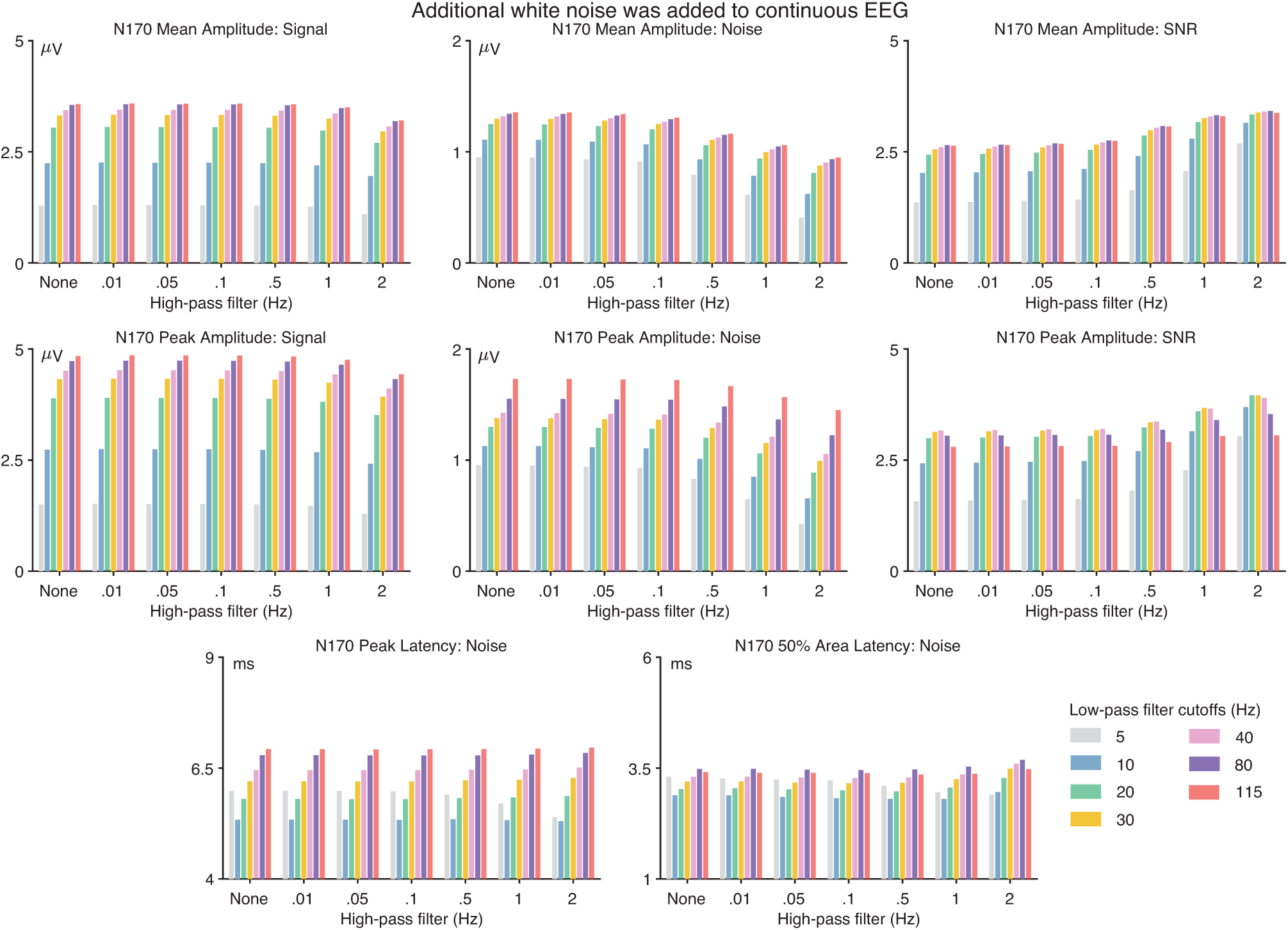
Data quality metrics for the N170 component when white noise (standard deviation = 7.07 µV) had been added to the continuous EEG prior to filtering. The format is identical to that of Figure 2 in the main document.

**Figure S5:**
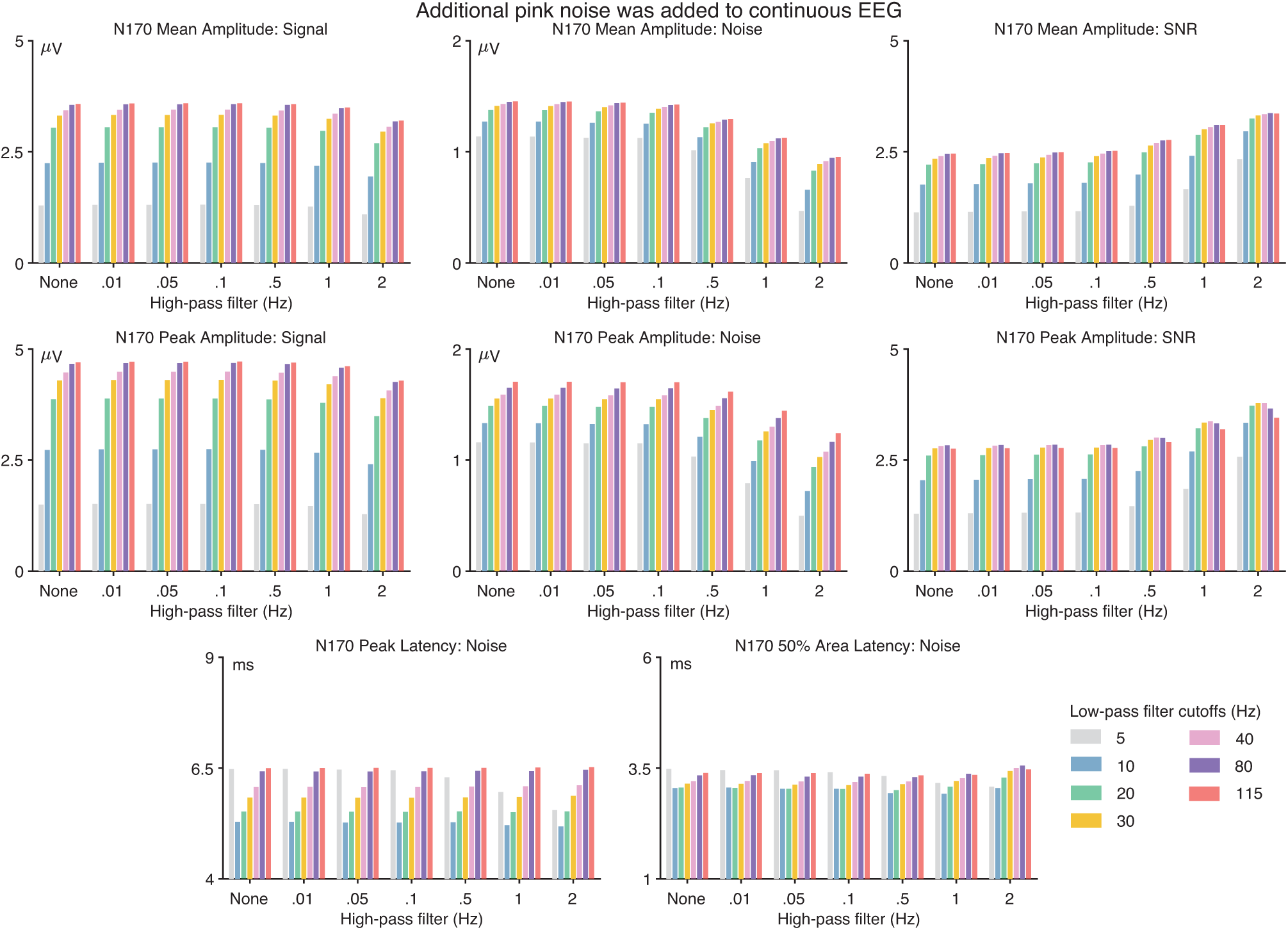
Data quality metrics for the N170 component when pink noise (standard deviation = 7.07 µV) had been added to the continuous EEG prior to filtering. The format is identical to that of Figure 2 in the main document.

**Figure S6:**
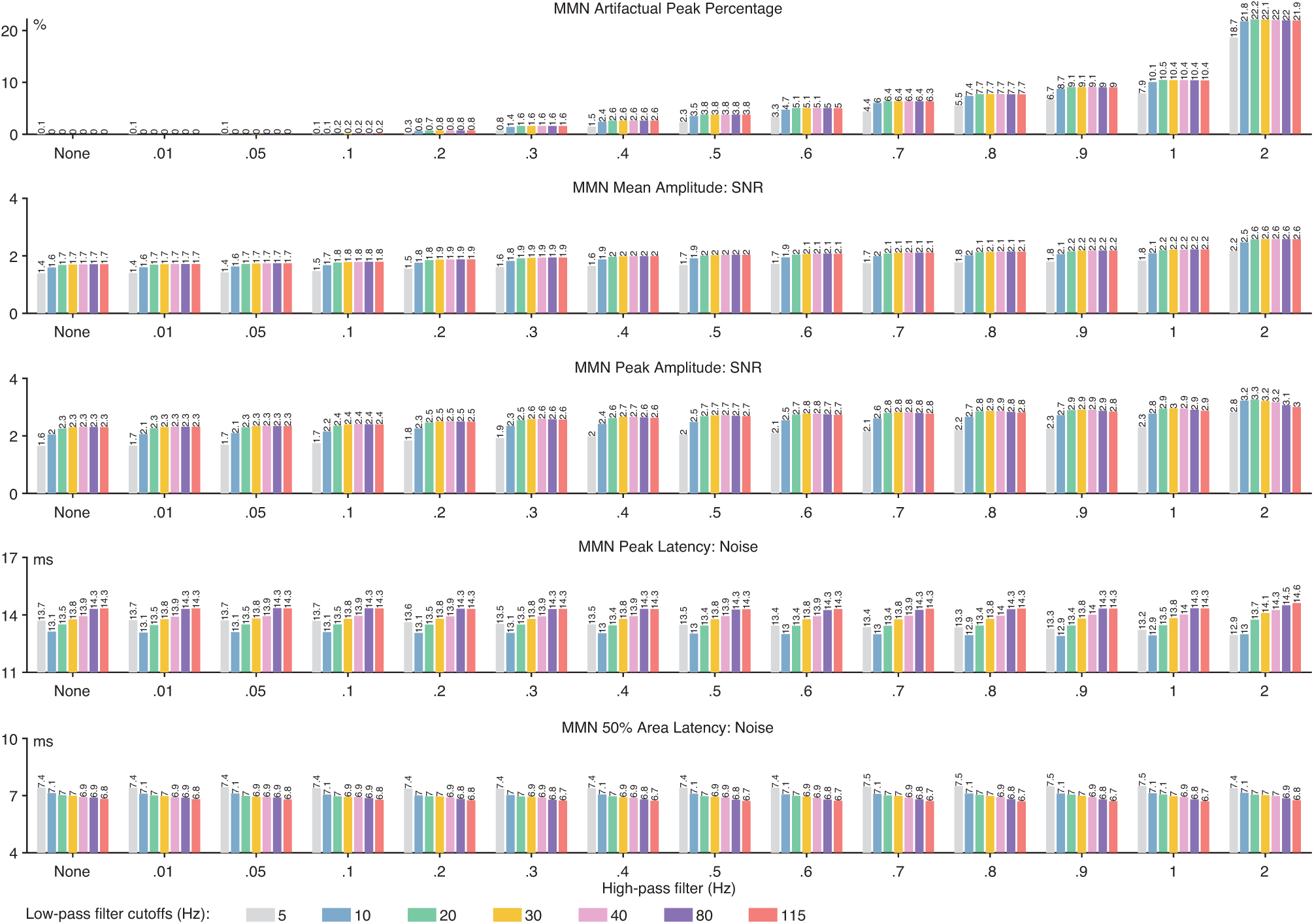
MMN artifactual peak percentage and data quality metrics. Data quality metrics are provided for each of the four scoring methods (mean amplitude, peak amplitude, peak latency and 50% area latency) and for each combination of high-pass filter cutoff frequency (0, 0.01, 0.05, 0.1, 0.2, 0.3, 0.4, 0.5, 0.6, 0.7, 0.8, 0.9, 1, and 2 Hz) and low-pass filter cutoff frequency (5, 10, 20, 30, 40, 80, and 115 Hz). The SNR values are unitless

**Figure S7:**
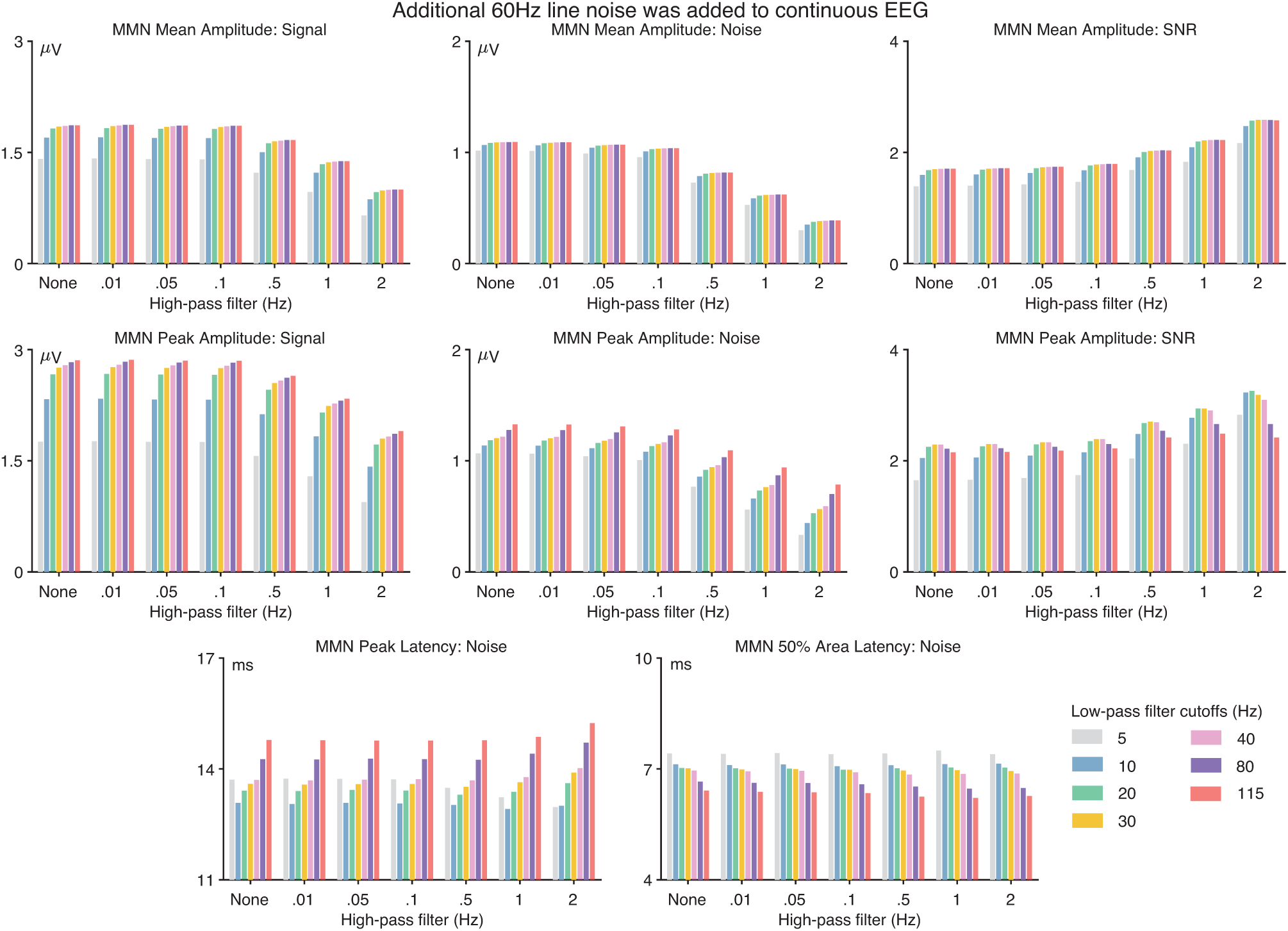
Data quality metrics for the MMN component when 60 Hz line noise (20 µV peak-to-peak amplitude) had been added to the continuous EEG prior to filtering. The format is identical to that of Figure 2 in the main document.

**Figure S8:**
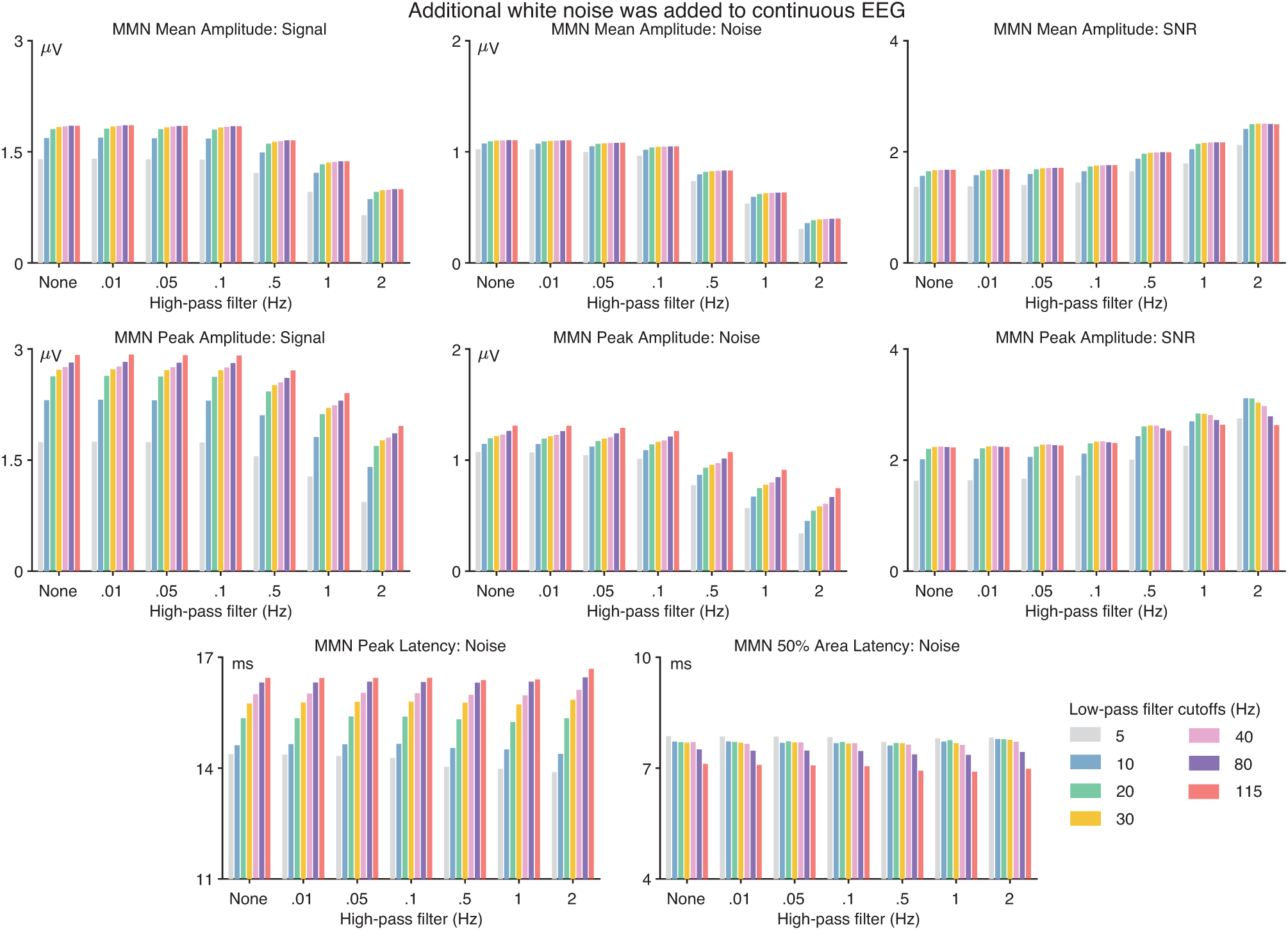
Data quality metrics for the MMN component when white noise (standard deviation = 7.07 µV) had been added to the continuous EEG prior to filtering. The format is identical to that of Figure 2 in the main document.

**Figure S9:**
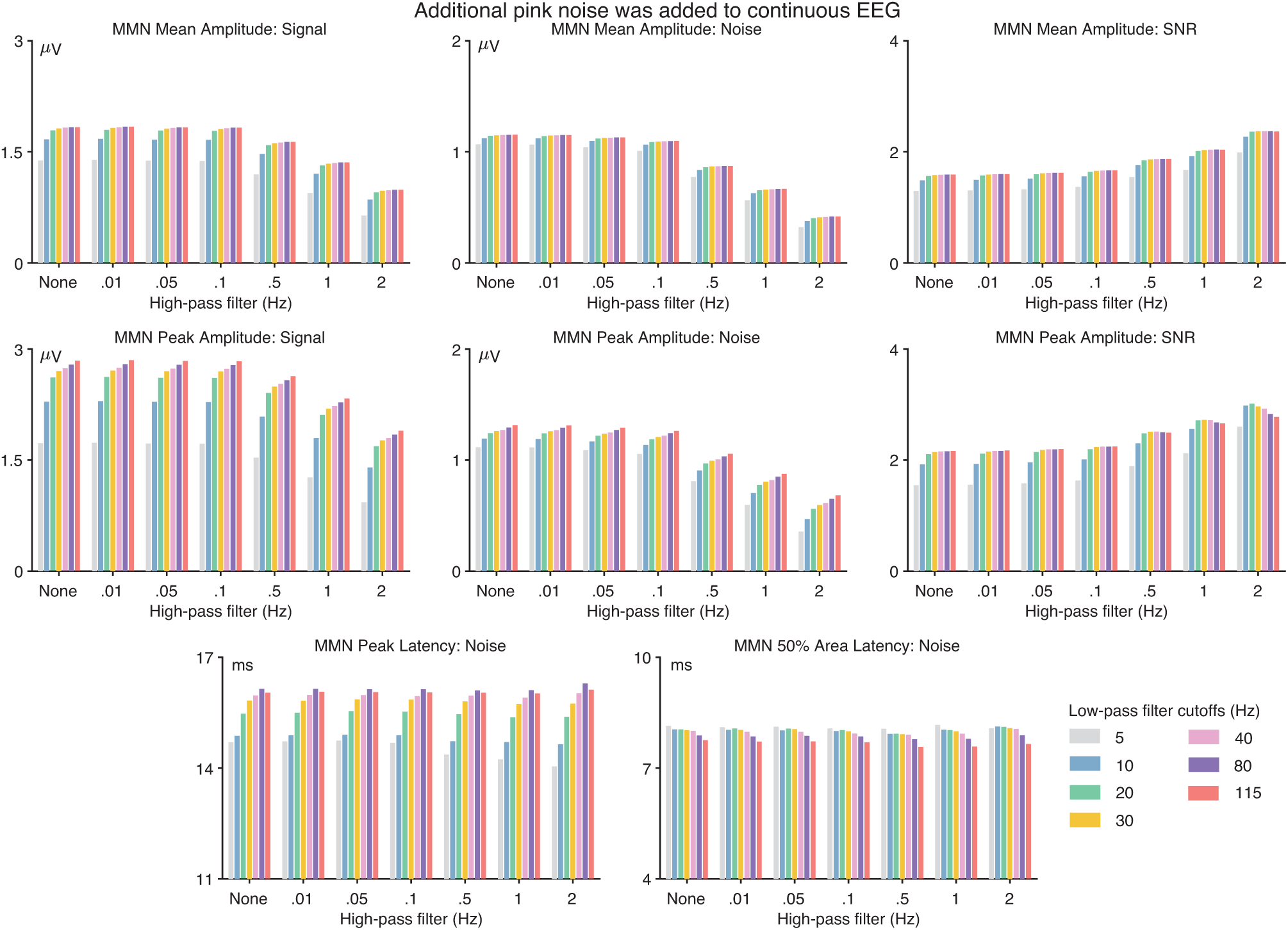
Data quality metrics for the MMN component when pink noise (standard deviation = 7.07 µV) had been added to the continuous EEG prior to filtering. The format is identical to that of Figure 2 in the main document.

**Figure S10:**
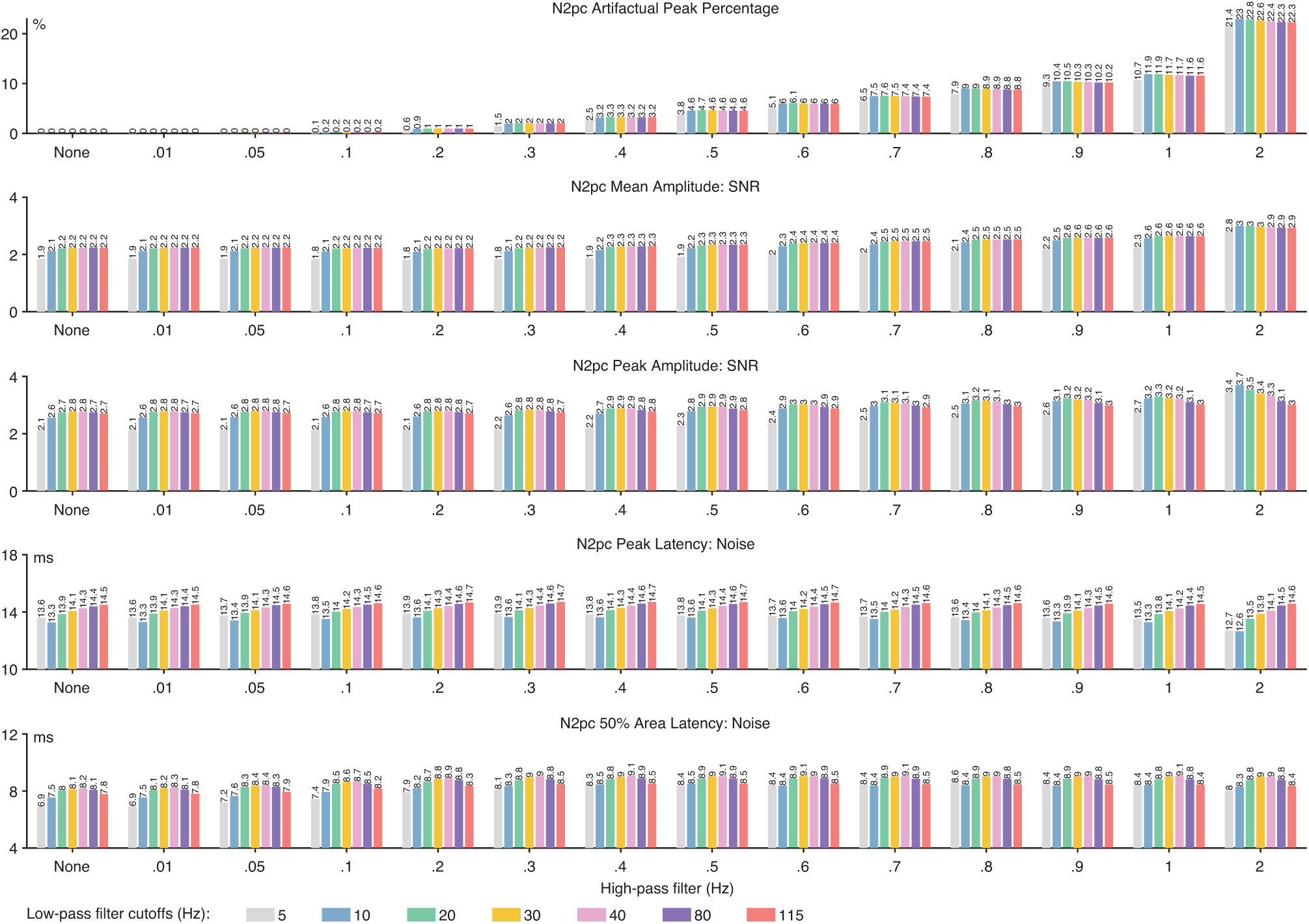
N2pc artifactual peak percentage and data quality metrics. Data quality metrics are provided for each of the four scoring methods (mean amplitude, peak amplitude, peak latency and 50% area latency) and for each combination of high-pass filter cutoff frequency (0, 0.01, 0.05, 0.1, 0.2, 0.3, 0.4, 0.5, 0.6, 0.7, 0.8, 0.9, 1, and 2 Hz) and low-pass filter cutoff frequency (5, 10, 20, 30, 40, 80, and 115 Hz). The SNR values are unitless.

**Figure S11:**
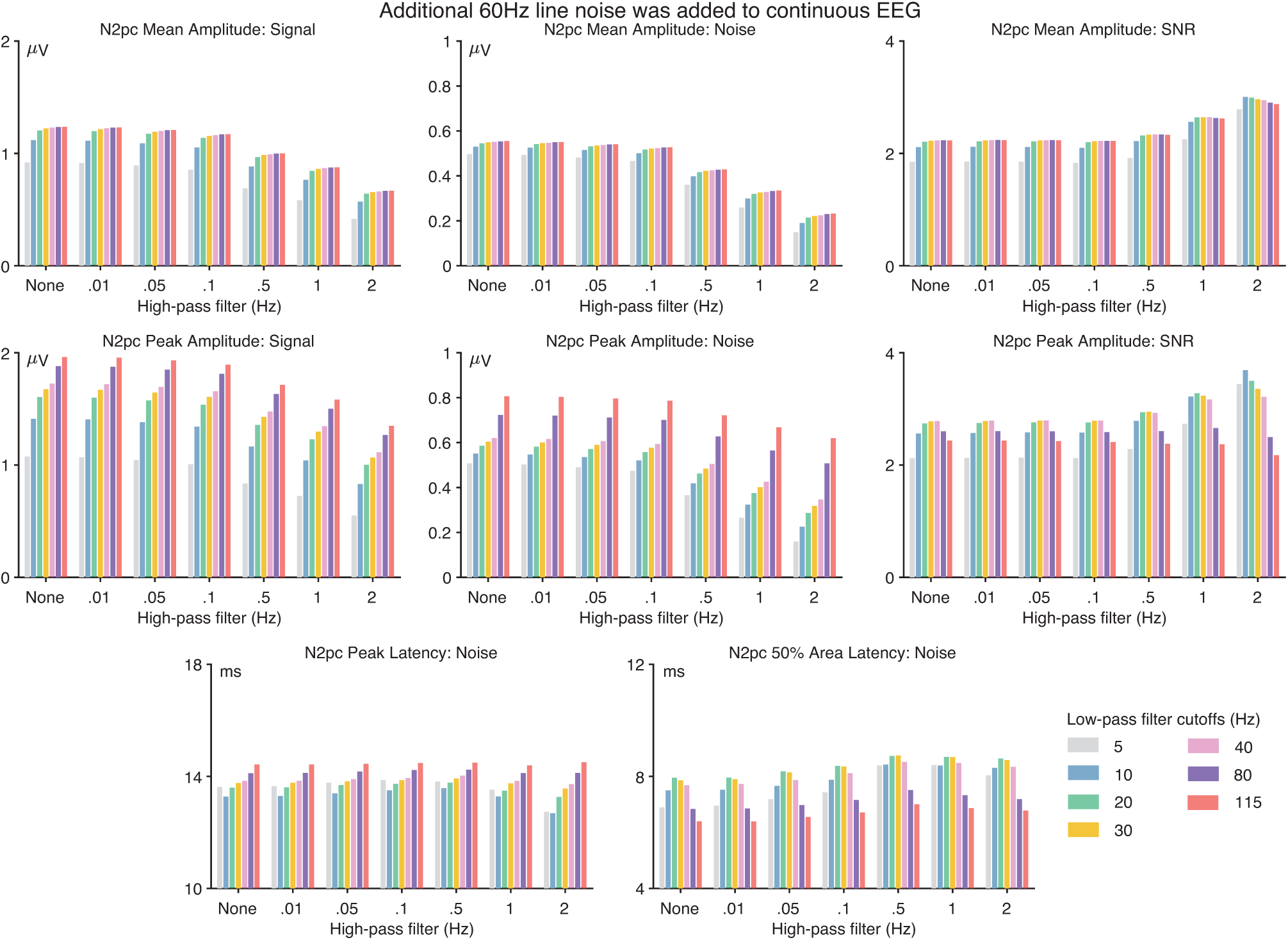
Data quality metrics for the N2pc component when 60 Hz line noise (20 µV peak-to-peak amplitude) had been added to the continuous EEG prior to filtering. The format is identical to that of Figure 2 in the main document.

**Figure S12:**
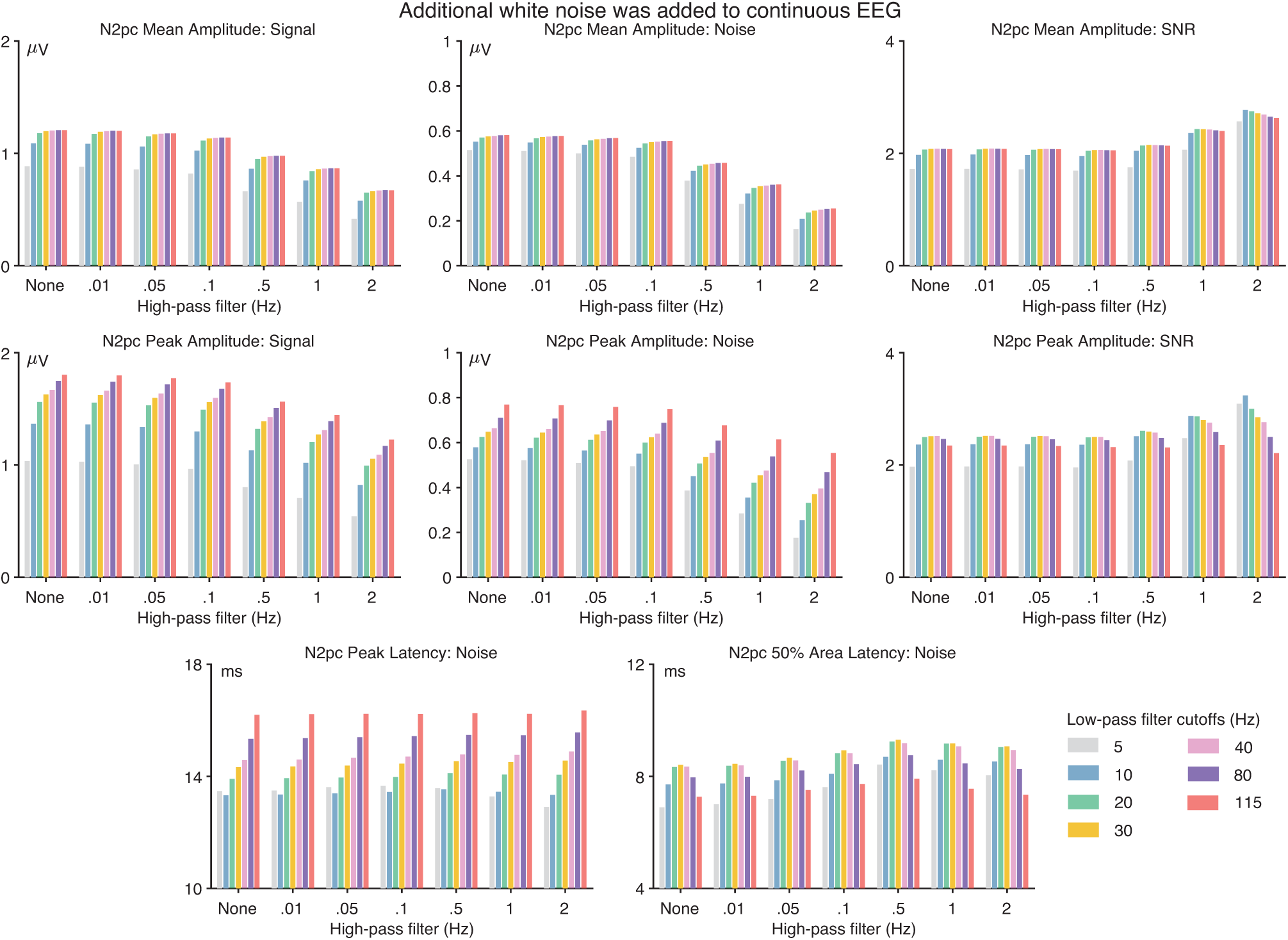
Data quality metrics for the N2pc component when white noise (standard deviation = 7.07 µV) had been added to the continuous EEG prior to filtering. The format is identical to that of Figure 2 in the main document.

**Figure S13:**
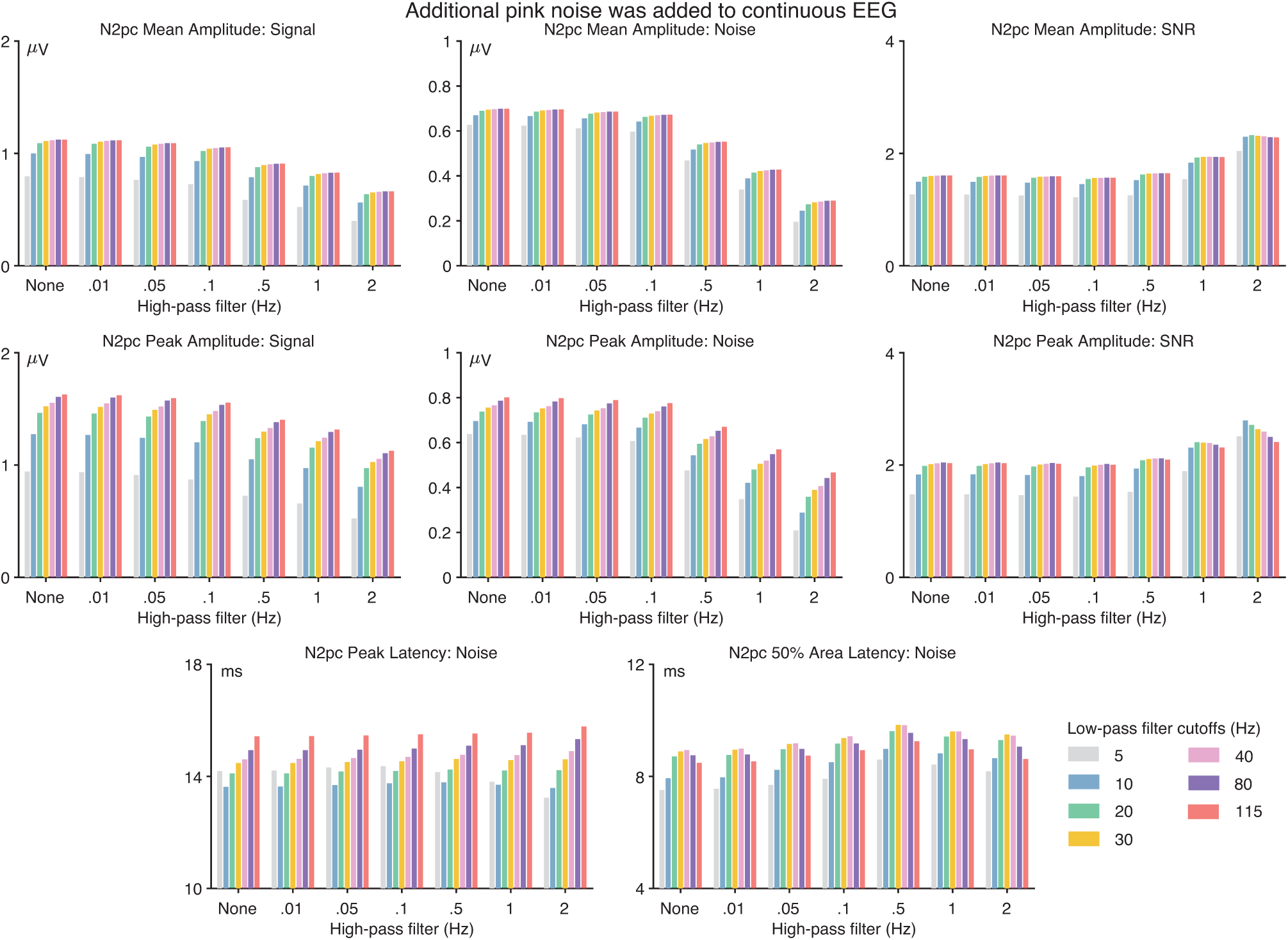
Data quality metrics for the N2pc component when pink noise (standard deviation = 7.07 µV) had been added to the continuous EEG prior to filtering. The format is identical to that of Figure 2 in the main document.

**Figure S14:**
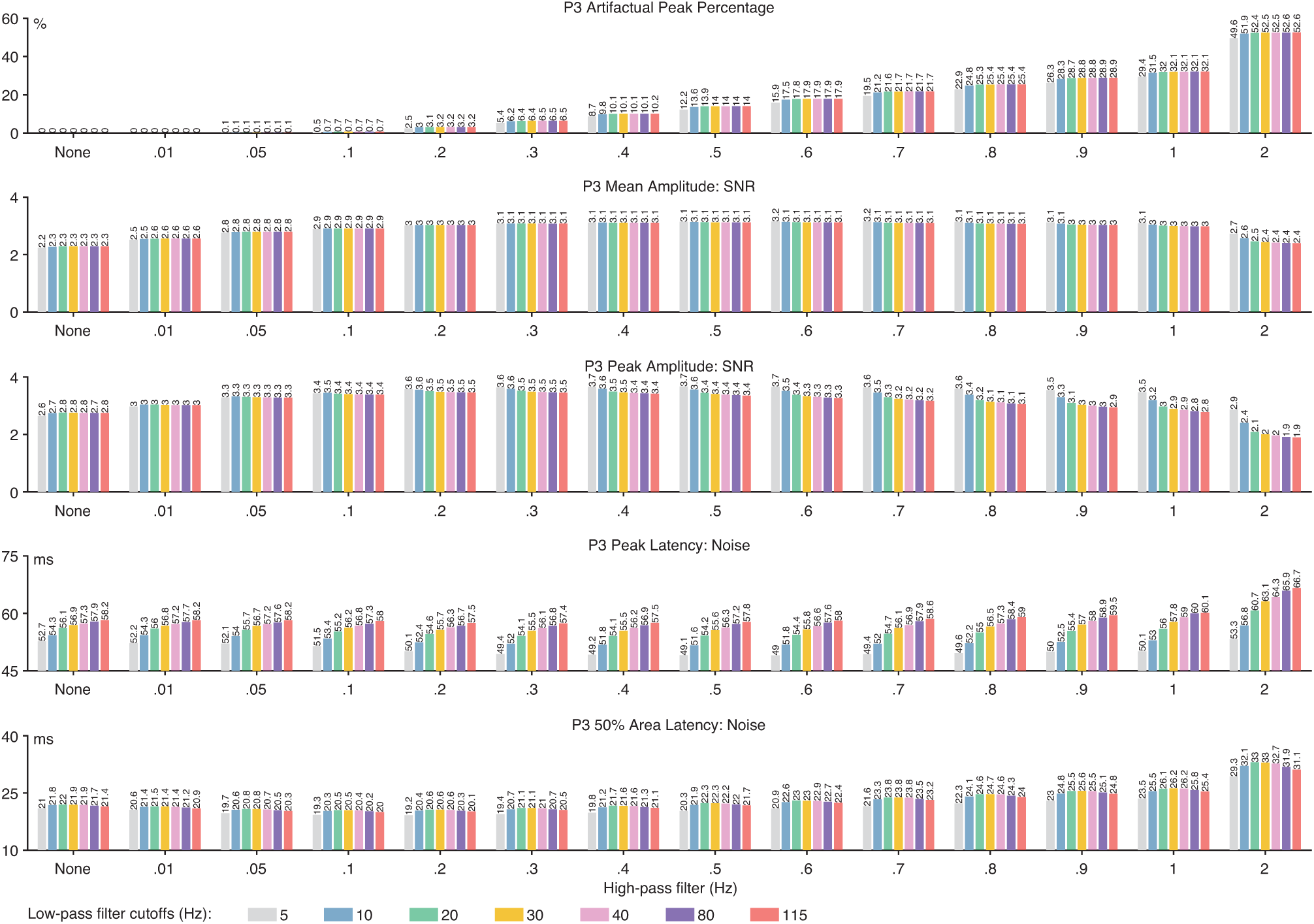
P3 artifactual peak percentage and data quality metrics. Data quality metrics are provided for each of the four scoring methods (mean amplitude, peak amplitude, peak latency and 50% area latency) and for each combination of high-pass filter cutoff frequency (0, 0.01, 0.05, 0.1, 0.2, 0.3, 0.4, 0.5, 0.6, 0.7, 0.8, 0.9, 1, and 2 Hz) and low-pass filter cutoff frequency (5, 10, 20, 30, 40, 80, and 115 Hz). The SNR values are unitless.

**Figure S15:**
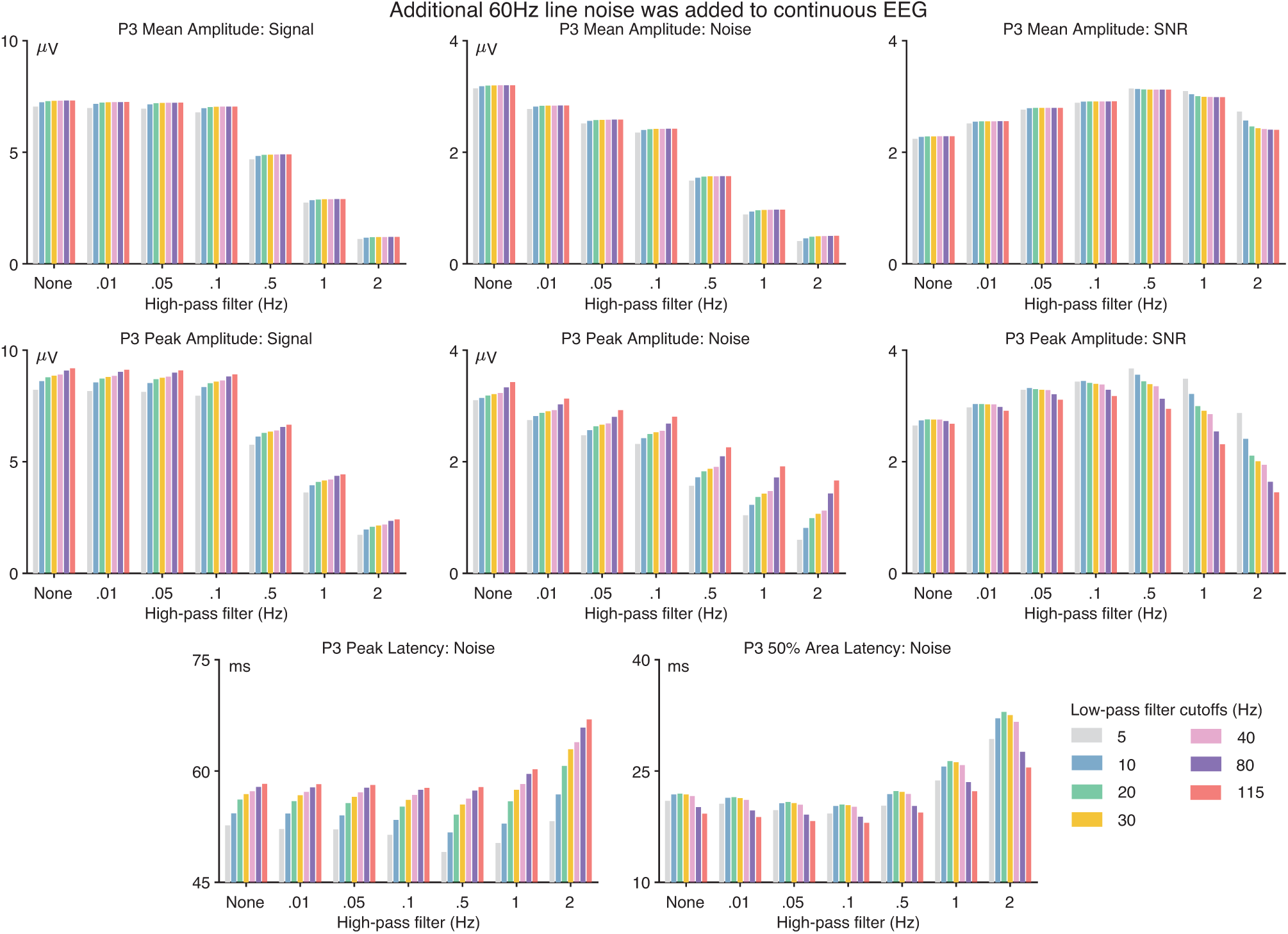
Data quality metrics for the P3 component when 60 Hz line noise (20 µV peak-to-peak amplitude) had been added to the continuous EEG prior to filtering. The format is identical to that of Figure 2 in the main document.

**Figure S16:**
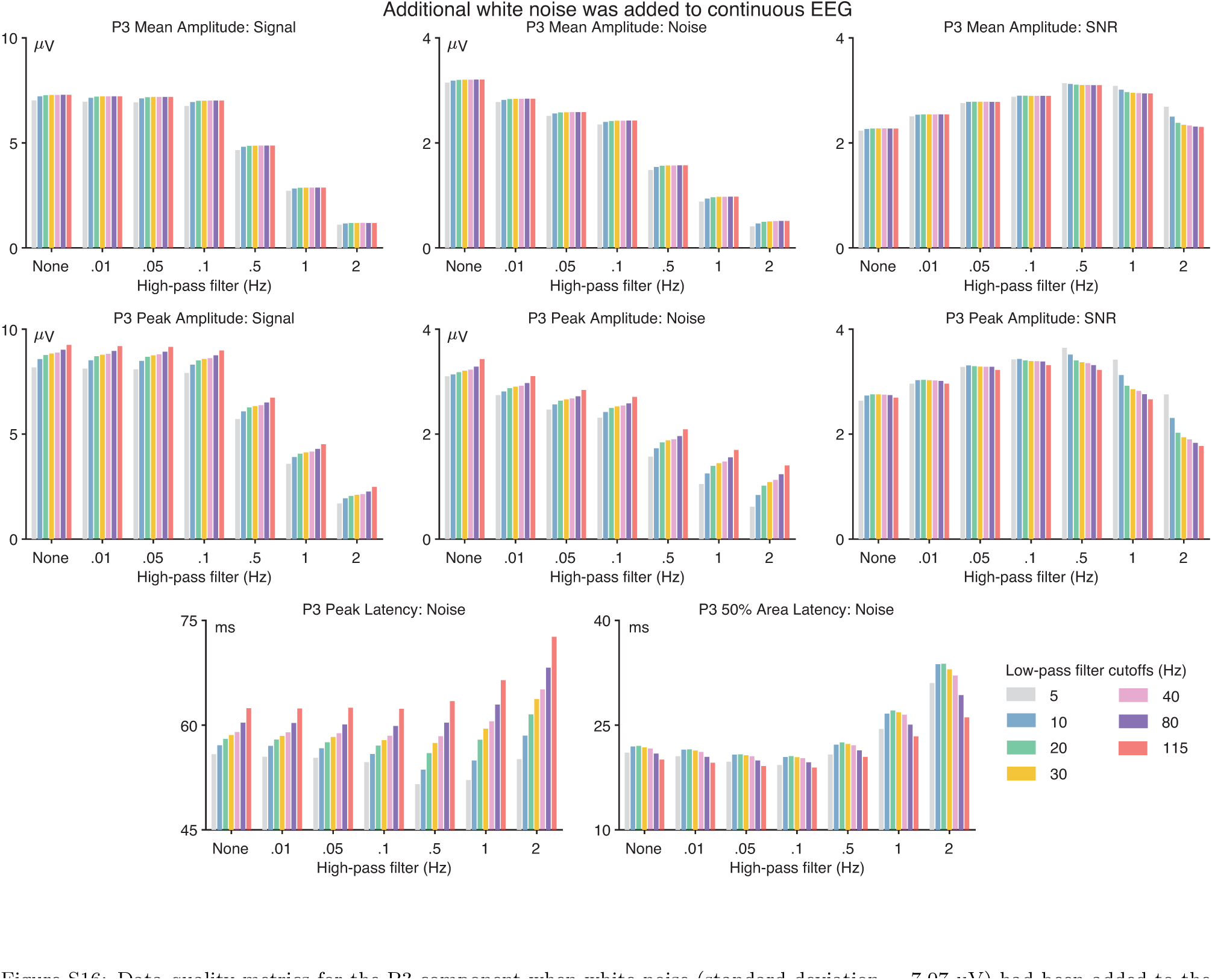
Data quality metrics for the P3 component when white noise (standard deviation = 7.07 µV) had been added to the continuous EEG prior to filtering. The format is identical to that of Figure 2 in the main document.

**Figure S17:**
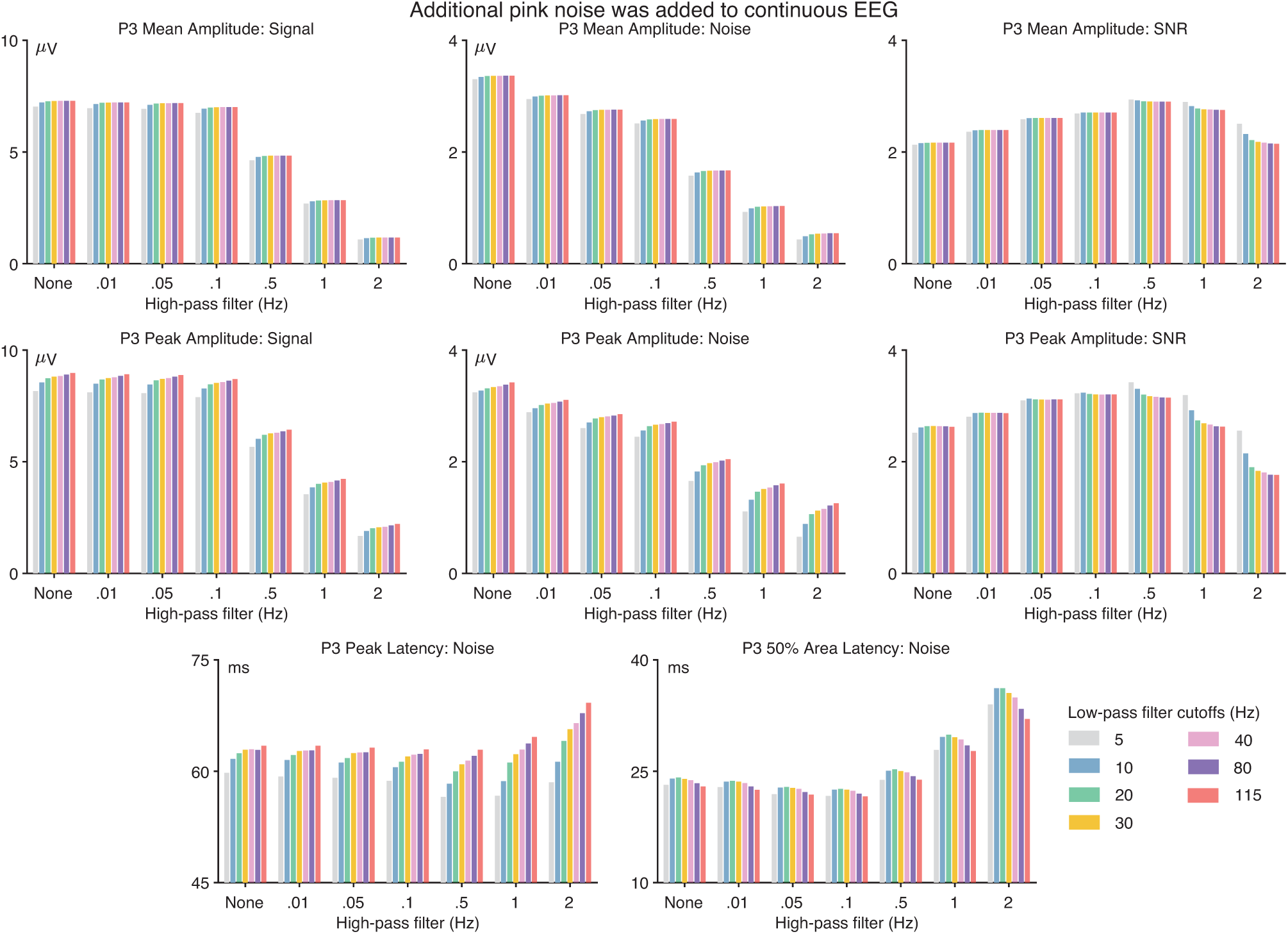
Data quality metrics for the P3 component when pink noise (standard deviation = 7.07 µV) had been added to the continuous EEG prior to filtering. The format is identical to that of Figure 2 in the main document.

**Figure S18:**
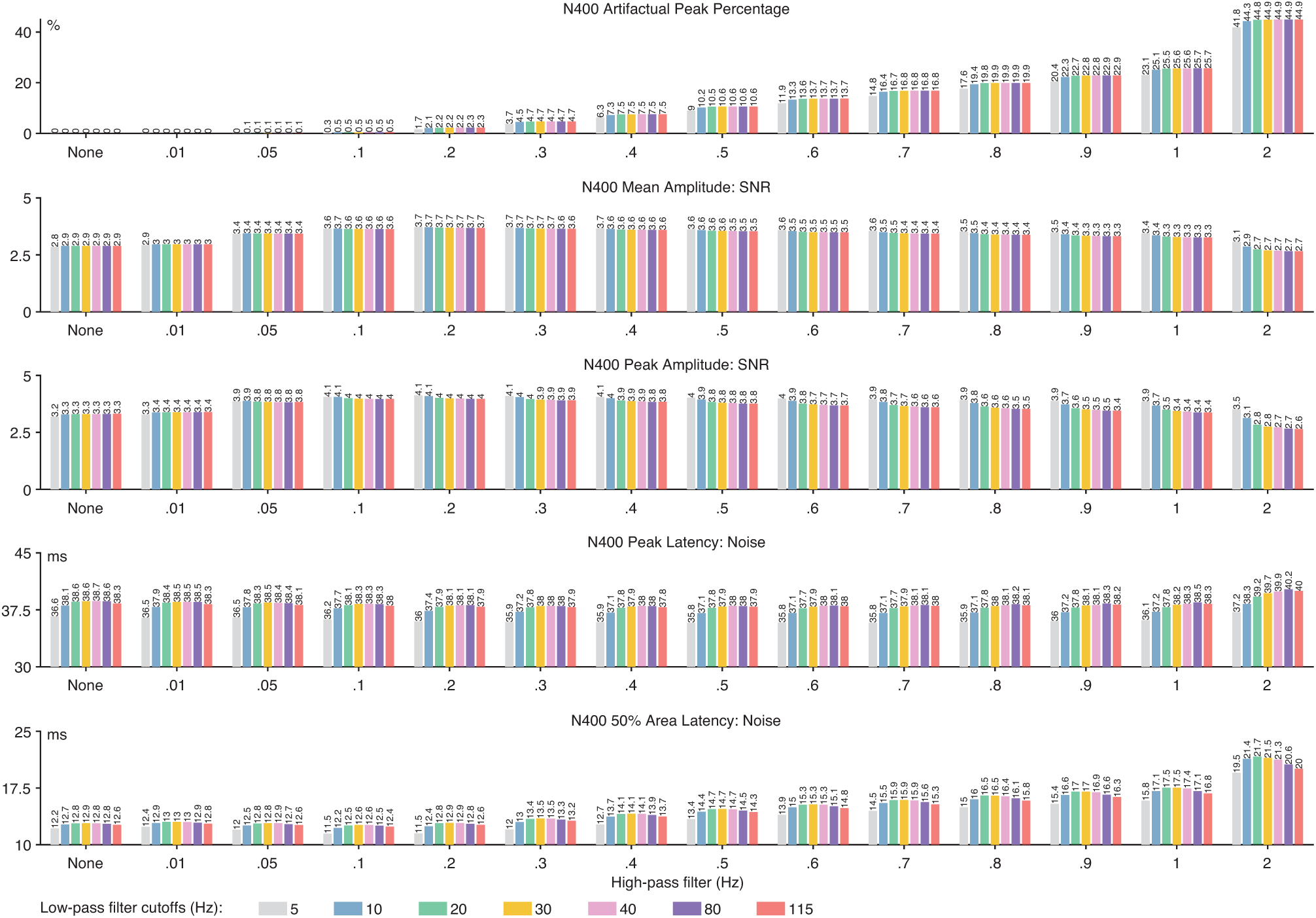
N400 artifactual peak percentage and data quality metrics. Data quality metrics are provided for each of the four scoring methods (mean amplitude, peak amplitude, peak latency and 50% area latency) and for each combination of high-pass filter cutoff frequency (0, 0.01, 0.05, 0.1, 0.2, 0.3, 0.4, 0.5, 0.6, 0.7, 0.8, 0.9, 1, and 2 Hz) and low-pass filter cutoff frequency (5, 10, 20, 30, 40, 80, and 115 Hz). The SNR values are unitless.

**Figure S19:**
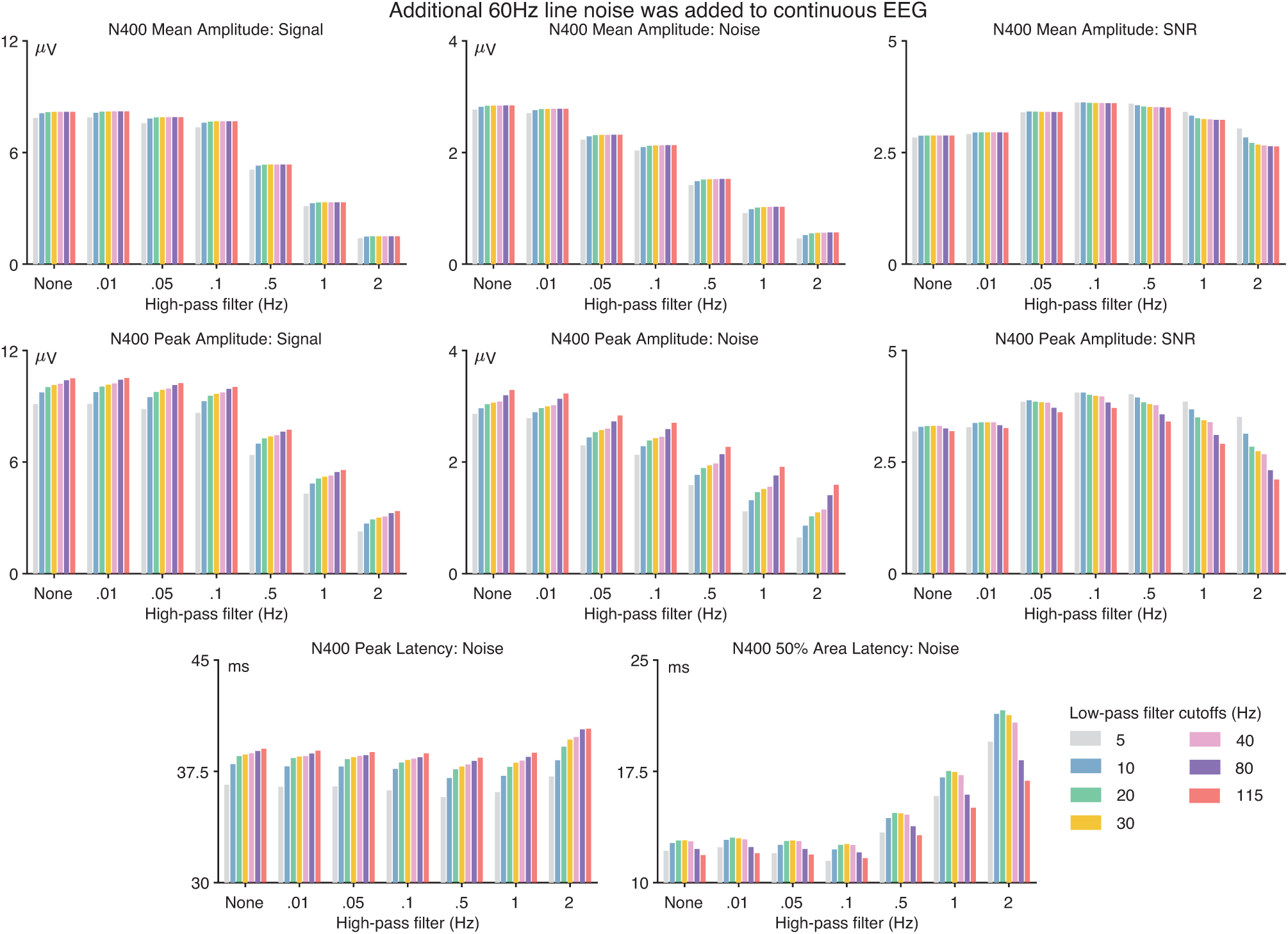
Data quality metrics for the N400 component when white noise (standard deviation = 7.07 µV) had been added to the continuous EEG prior to filtering. The format is identical to that of Figure 2 in the main document.

**Figure S20:**
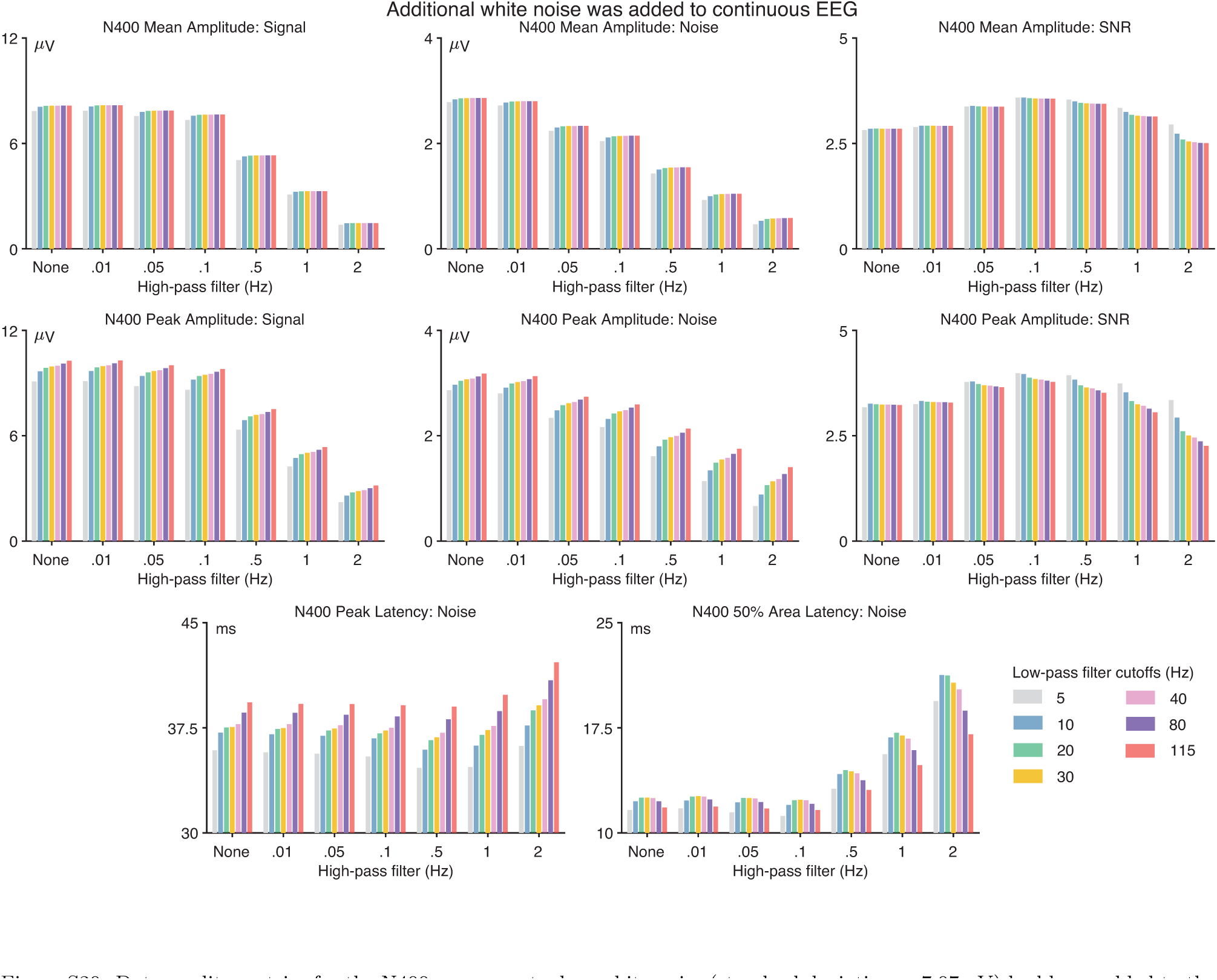
Data quality metrics for the N400 component when white noise (standard deviation = 7.07 µV) had been added to the continuous EEG prior to filtering.

**Figure S21:**
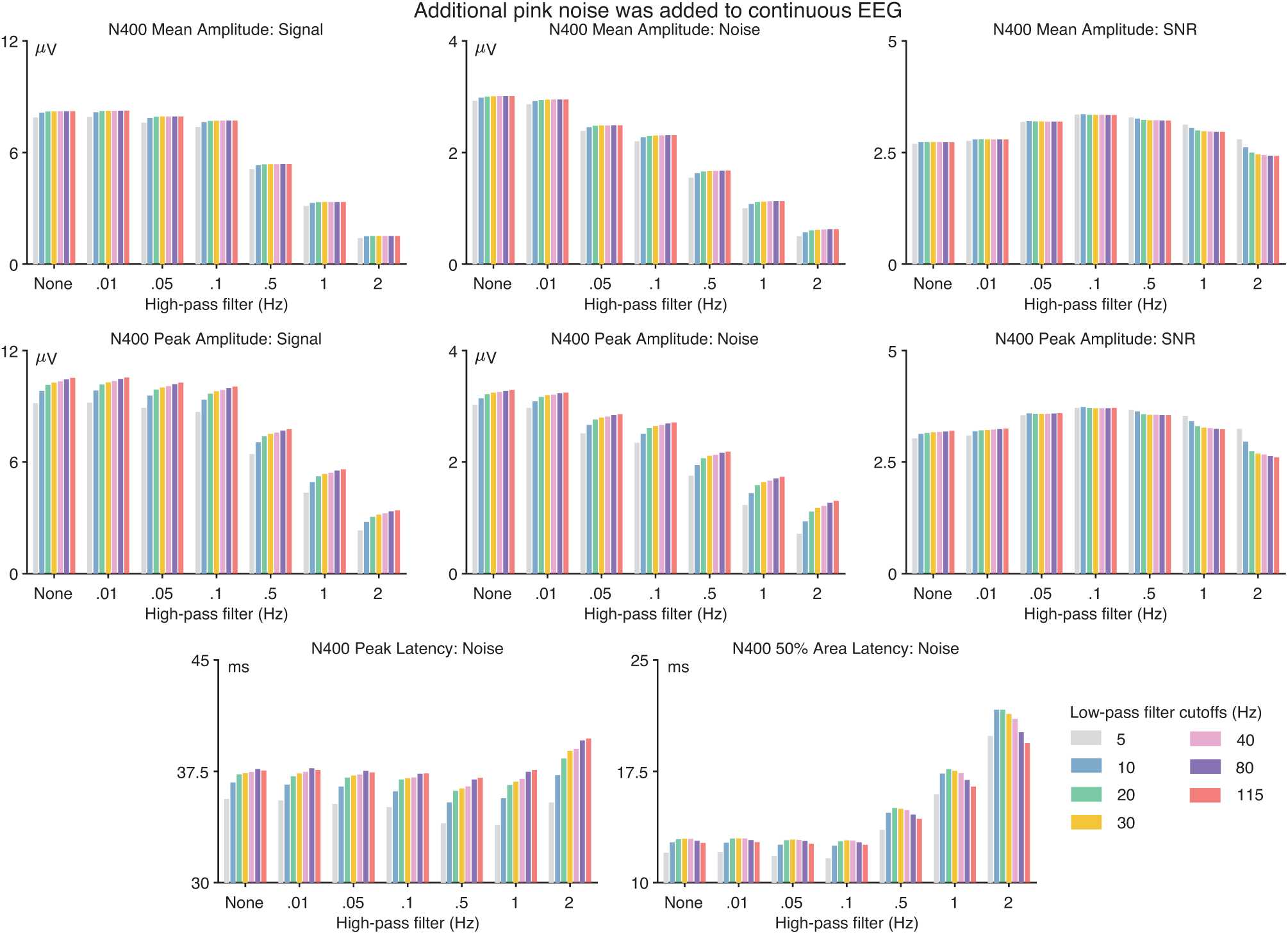
Data quality metrics for the N400 component when pink noise (standard deviation = 7.07 µV) had been added to the continuous EEG prior to filtering. The format is identical to that of Figure 2 in the main document.

**Figure S22:**
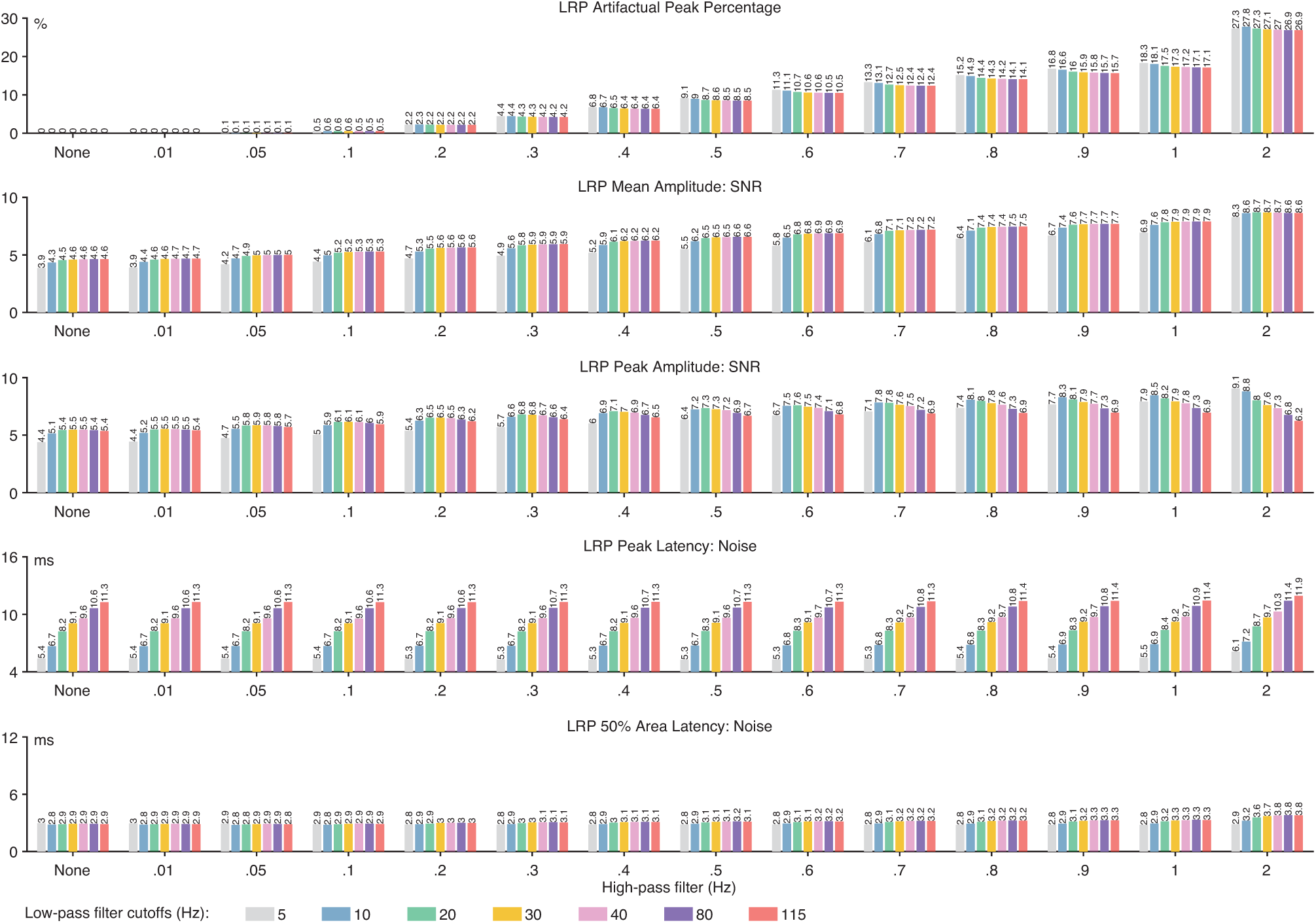
LRP artifactual peak percentage and data quality metrics. Data quality metrics are provided for each of the four scoring methods (mean amplitude, peak amplitude, peak latency and 50% area latency) and for each combination of high-pass filter cutoff frequency (0, 0.01, 0.05, 0.1, 0.2, 0.3, 0.4, 0.5, 0.6, 0.7, 0.8, 0.9, 1, and 2 Hz) and low-pass filter cutoff frequency (5, 10, 20, 30, 40, 80, and 115 Hz). The SNR values unitless.

**Figure S23:**
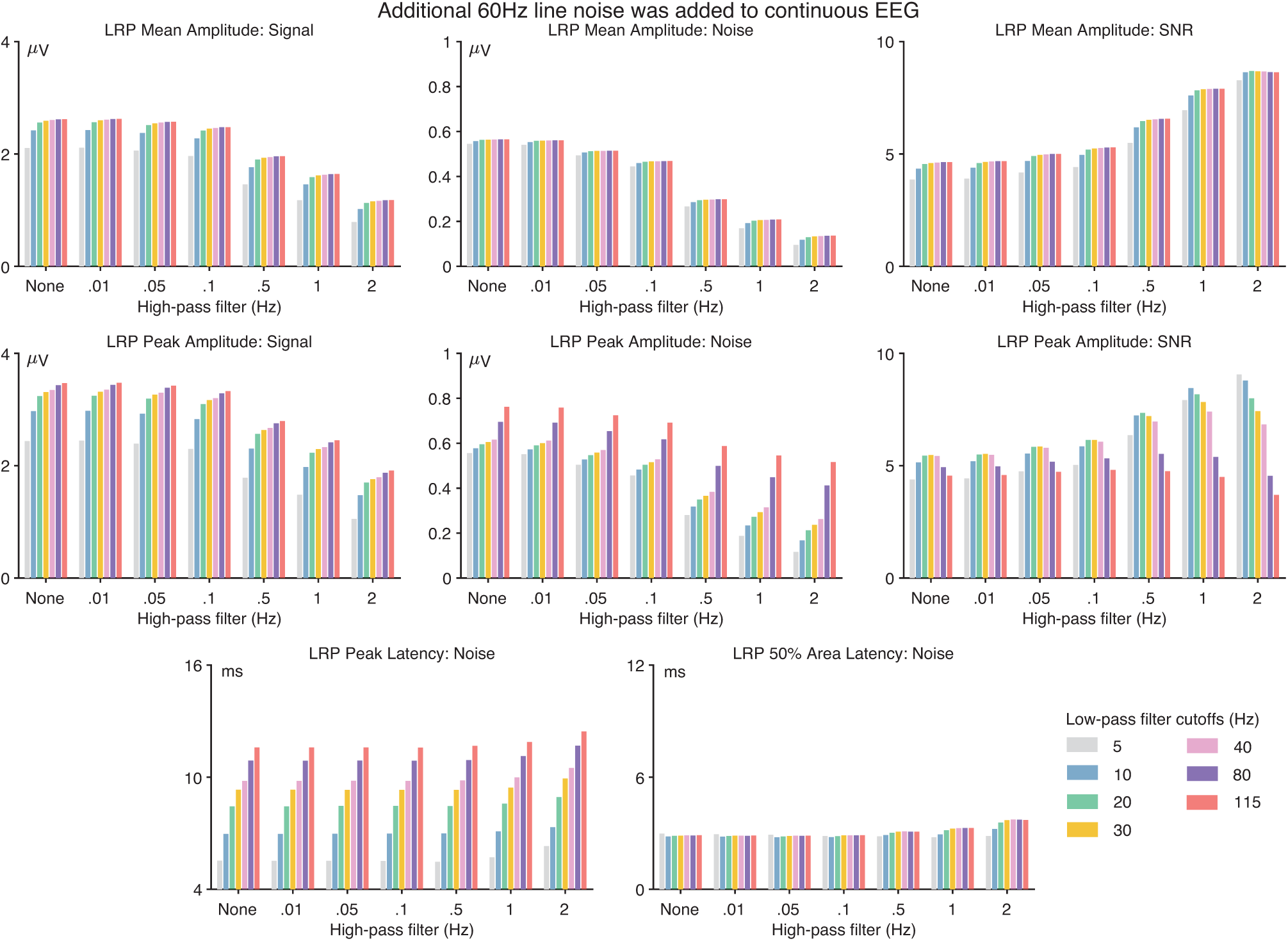
Data quality metrics for the LRP component when 60 Hz line noise (20 µV peak-to-peak amplitude) had been added to the continuous EEG prior to filtering. The format is identical to that of Figure 2 in the main document.

**Figure S24:**
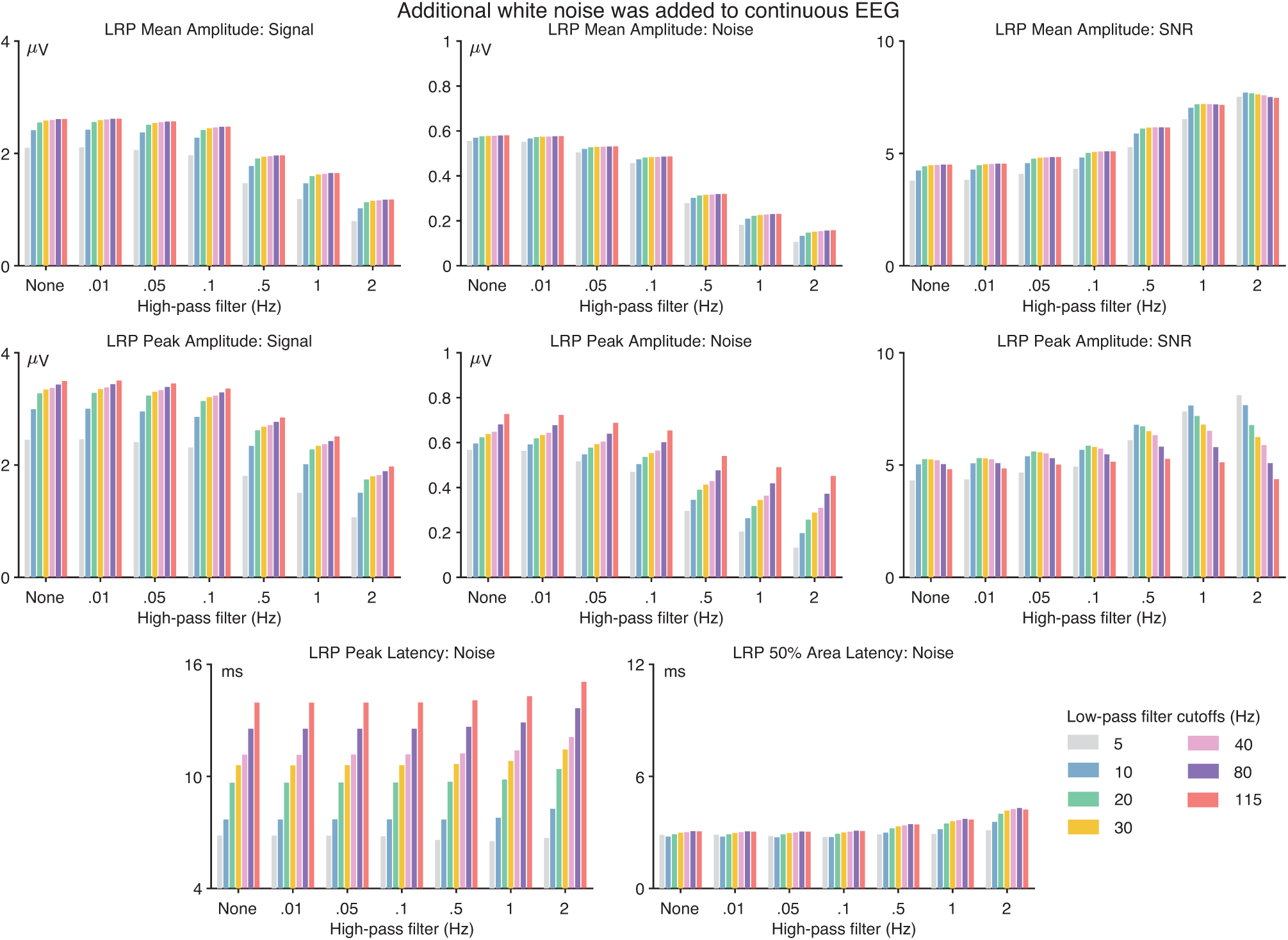
Data quality metrics for the LRP component when white noise (standard deviation = 7.07 µV) had been added to the continuous EEG prior to filtering. The format is identical to that of Figure 2 in the main document.

**Figure S25:**
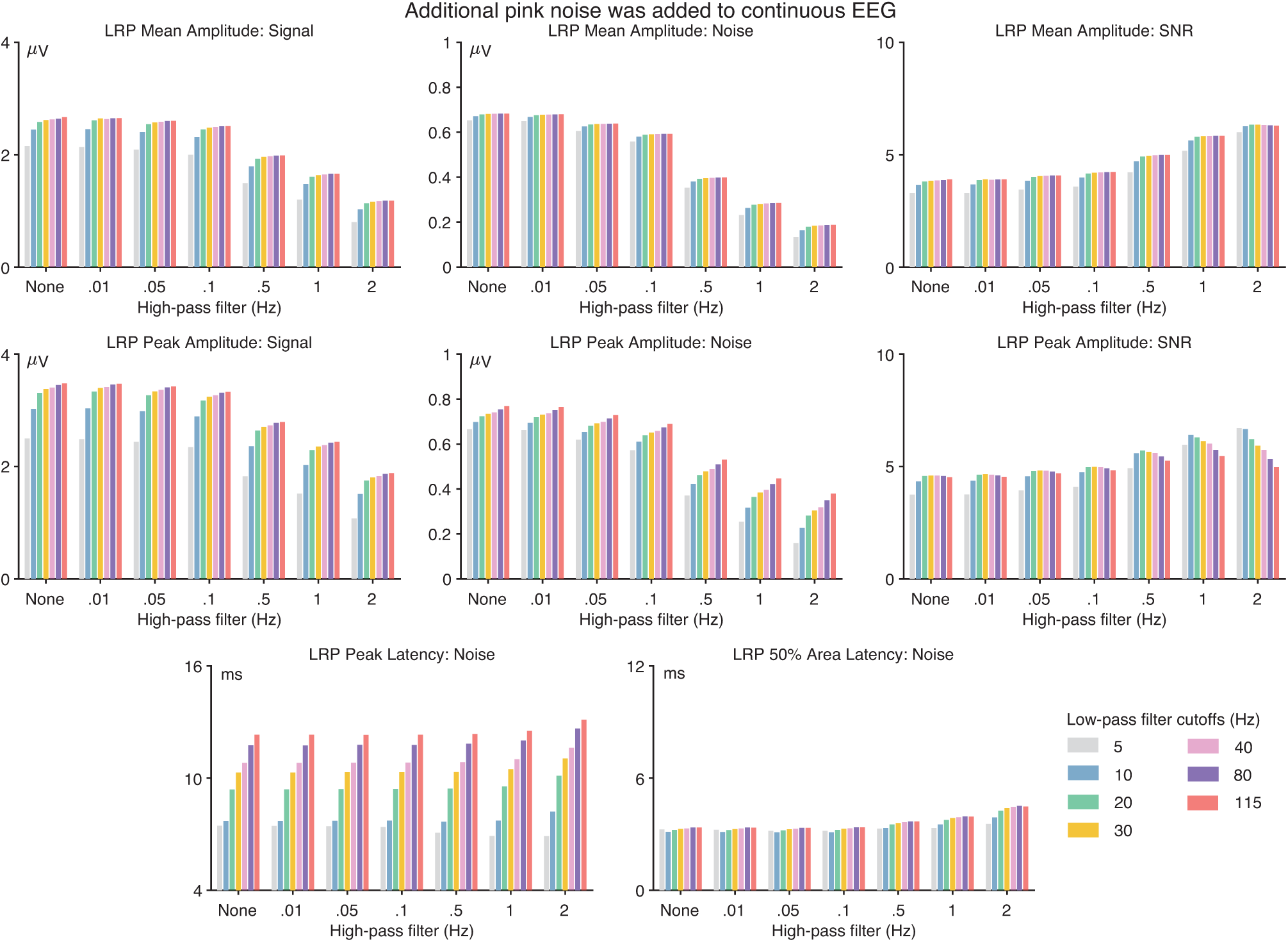
Data quality metrics for the LRP component when pink noise (standard deviation = 7.07 µV) had been added to the continuous EEG prior to filtering. The format is identical to that of Figure 2 in the main document.

**Figure S26:**
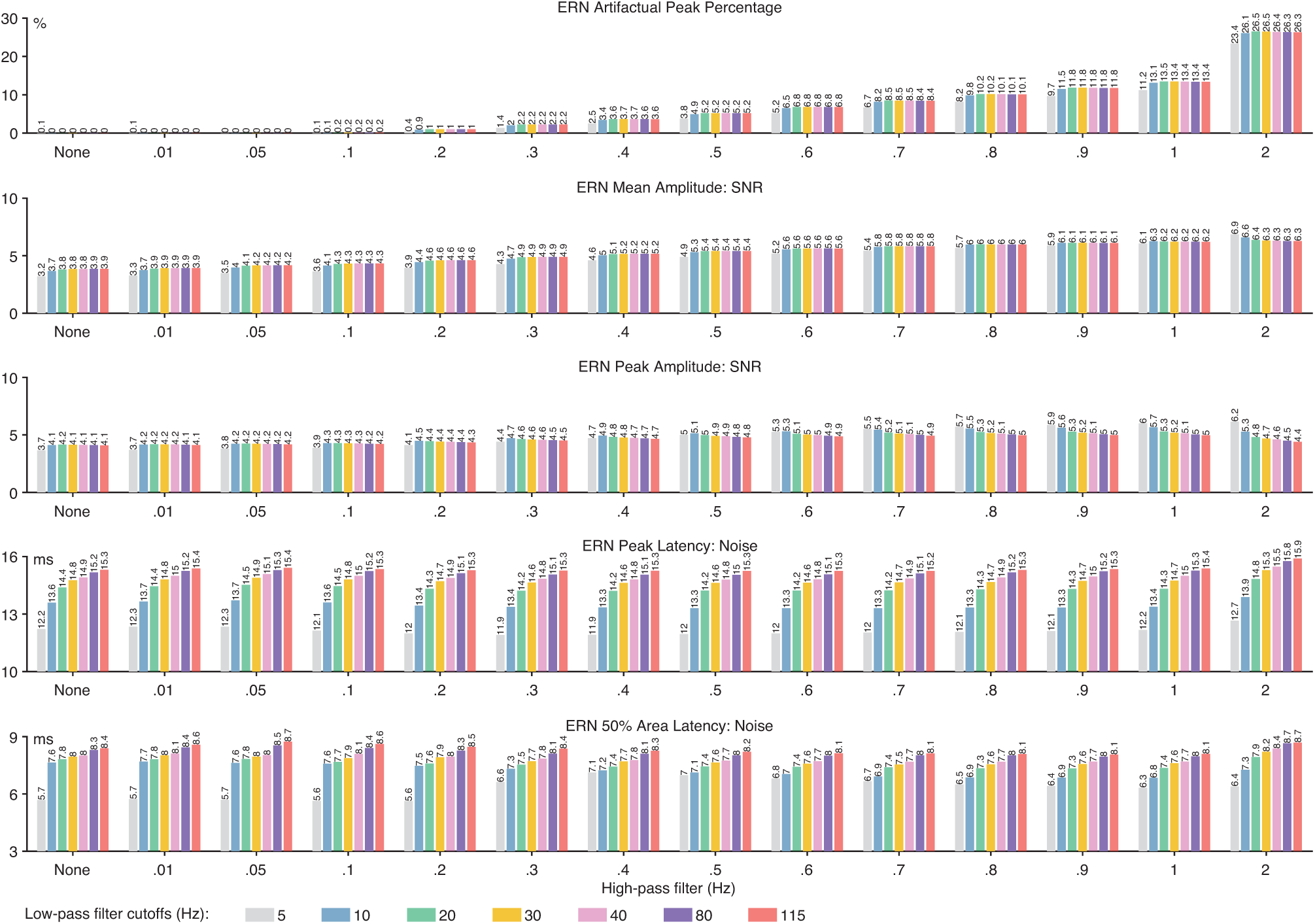
ERN artifactual peak percentage and data quality metrics. Data quality metrics are provided for each of the four scoring methods (mean amplitude, peak amplitude, peak latency and 50% area latency) and for each combination of high-pass filter cutoff frequency (0, 0.01, 0.05, 0.1, 0.2, 0.3, 0.4, 0.5, 0.6, 0.7, 0.8, 0.9, 1, and 2 Hz) and low-pass filter cutoff frequency (5, 10, 20, 30, 40, 80, and 115 Hz). The SNR values are unitless.

**Figure S27:**
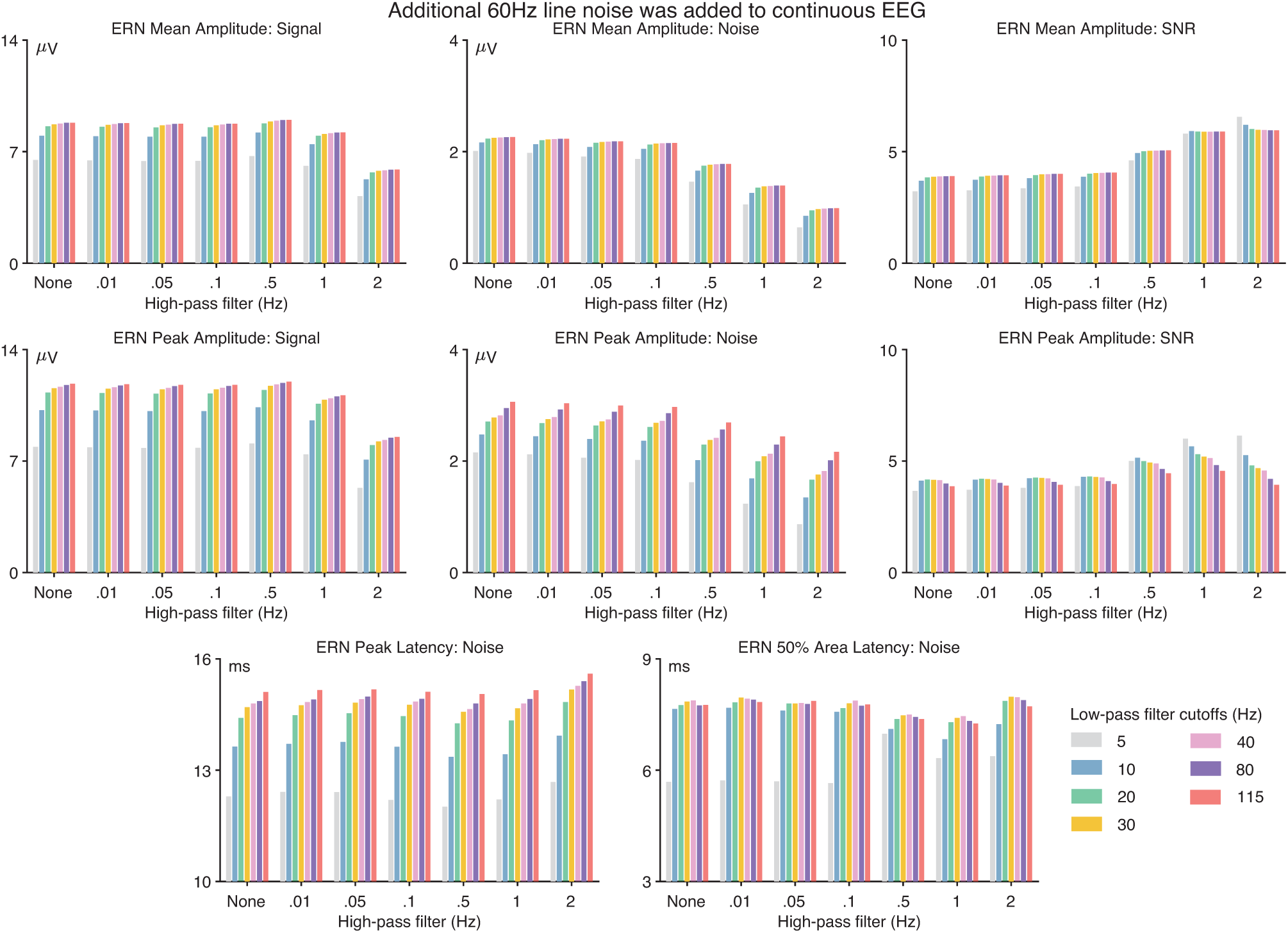
Data quality metrics for the ERN component when 60 Hz line noise (20 µV peak-to-peak amplitude) had been added to the continuous EEG prior to filtering. The format is identical to that of Figure 2 in the main document.

**Figure S28:**
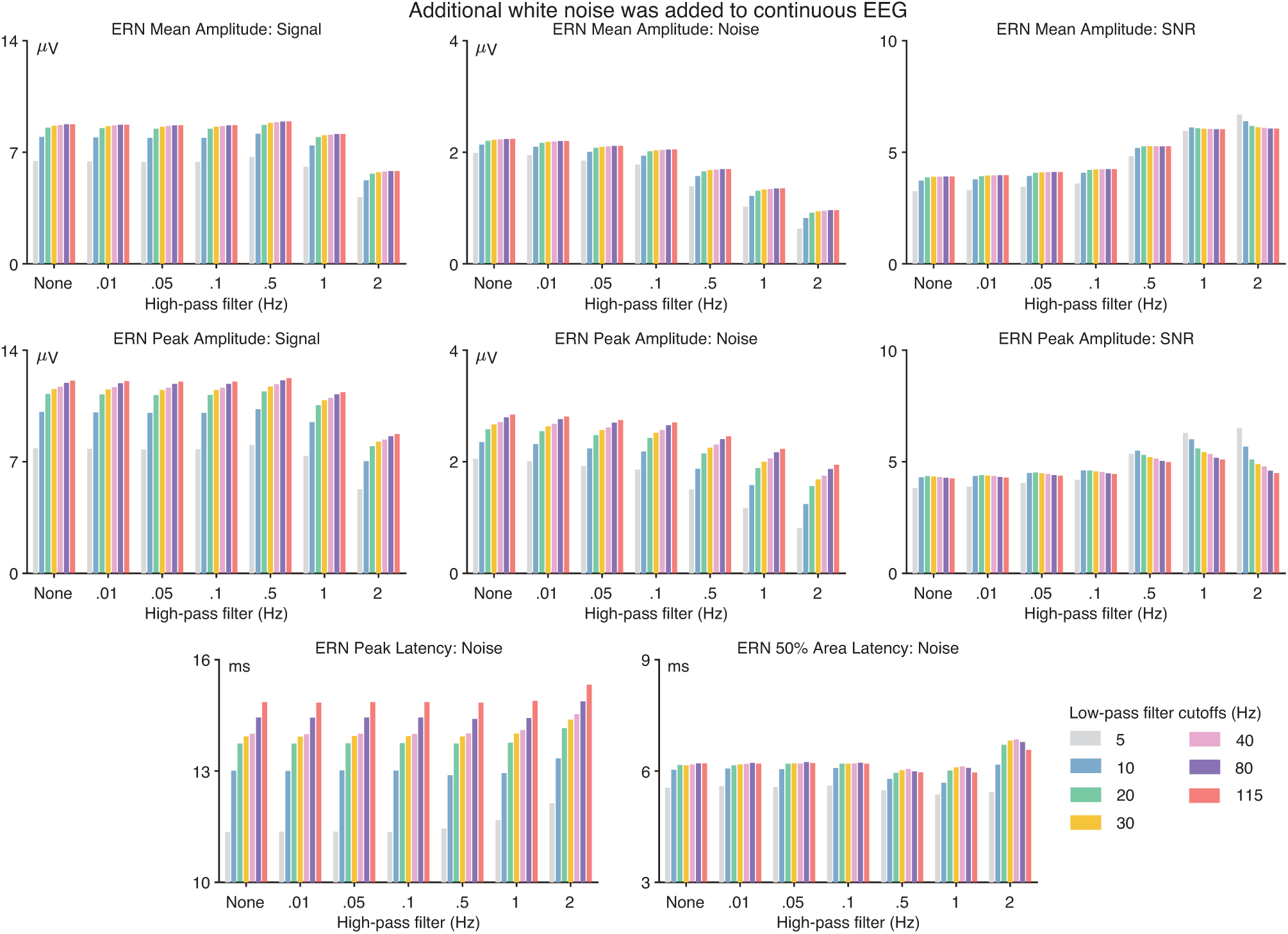
Data quality metrics for the ERN component when white noise (standard deviation = 7.07 µV) had been added to the continuous EEG prior to filtering. The format is identical to that of Figure 2 in the main document.

**Figure S29:**
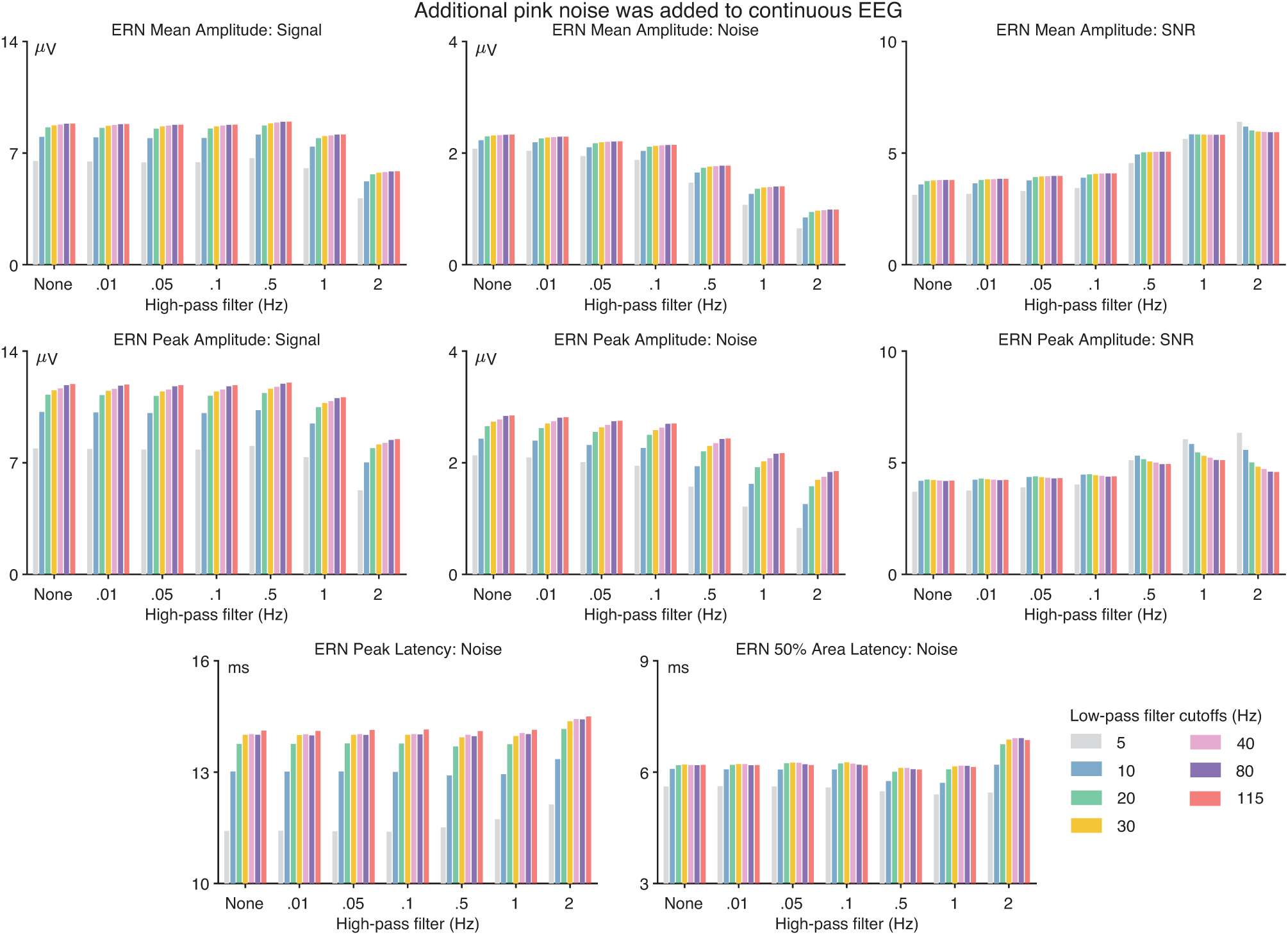
Data quality metrics for the ERN component when pink noise (standard deviation = 7.07 µV) had been added to the continuous EEG prior to filtering. The format is identical to that of Figure 2 in the main document.

## Footnotes

1 Although the RMS(SME) was virtually identical for a 10 Hz or 20 Hz cutoff, the 20 Hz cutoff produces less temporal smearing of the waveform, so 20 Hz may be slightly preferable to 10 Hz.

## Notes

### Competing Interest Statement

The authors have declared no competing interest.

## References

1. Acunzo, D. J., MacKenzie, G., & van Rossum, M. C. (2012). Systematic biases in early ERP and ERF components as a result of high-pass filtering. Journal of Neuroscience Methods, 209, 212–218. doi:10.1016/j.jneumeth.2012.06.011.

2. Clayson, P. E., Baldwin, S. A., & Larson, M. J. (2013). How does noise affect amplitude and latency measurement of event-related potentials (ERPs)? a methodological critique and simulation study. Psychophysiology, 50, 174–186. doi:10.1111/psyp.12001.

3. Debnath, R., Buzzell, G. A., Morales, S., Bowers, M. E., Leach, S. C., & Fox, N. A. (2020). The maryland analysis of developmental EEG (MADE) pipeline. Psychophysiology, 57, e13580. doi:10.1111/psyp.13580.

4. Delorme, A., & Makeig, S. (2004). EEGLAB: an open source toolbox for analysis of single-trial EEG dynamics including independent component analysis. Journal of Neuroscience Methods, 134, 9–21. doi:10.1016/j.jneumeth.2003.10.009.

5. van Driel, J., Olivers, C. N., & Fahrenfort, J. J. (2021). High-pass filtering artifacts in multivariate classification of neural time series data. Journal of Neuroscience Methods, 352, 109080. doi:10.1016/j.jneumeth.2021.109080.

6. Duncan, C. C., Barry, R. J., Connolly, J. F., Fischer, C., Michie, P. T., Näätänen, R., Polich, J., Reinvang, I., & Van Petten, C. (2009). Event-related potentials in clinical research: guidelines for eliciting, recording, and quantifying mismatch negativity, P300, and N400. Clinical Neurophysiology, 120, 1883–1908. doi:10.1016/j.clinph.2009.07.045.

7. Gabard-Durnam, L. J., Mendez Leal, A. S., Wilkinson, C. L., & Levin, A. R. (2018). The harvard automated processing pipeline for electroencephalography (HAPPE): standardized processing software for developmental and high-artifact data. Frontiers in Neuroscience, 12, 97. doi:10.3389/fnins.2018.00097.

8. Hamming, R. W. (1998). Digital filters. Courier Corporation.

9. Kam, J. W., Griffin, S., Shen, A., Patel, S., Hinrichs, H., Heinze, H.-J., Deouell, L. Y., & Knight, R. T. (2019). Systematic comparison between a wireless EEG system with dry electrodes and a wired EEG system with wet electrodes. NeuroImage, 184, 119–129. doi:/10.1016/j.neuroimage.2018.09.012.

10. Kappenman, E. S., Farrens, J. L., Zhang, W., Stewart, A. X., & Luck, S. J. (2021). ERP CORE: An open resource for human event-related potential research. NeuroImage, 225, 117465. doi:10.1016/j.neuroimage.2020.117465.

11. Klug, M., & Gramann, K. (2021). Identifying key factors for improving ICA-based decomposition of EEG data in mobile and stationary experiments. European Journal of Neuroscience, 54, 8406–8420. doi:10.1111/ejn.14992.

12. Li, G.-L., Wu, J.-T., Xia, Y.-H., He, Q.-G., & Jin, H.-G. (2020). Review of semi-dry electrodes for EEG recording. Journal of Neural Engineering, 17, 051004. doi:10.1088/1741-2552/abbd50.

13. Lopez-Calderon, J., & Luck, S. J. (2014). ERPLAB: an open-source toolbox for the analysis of event-related potentials. Frontiers in Human Neuroscience, 8, 213. doi:10.3389/fnhum.2014.002.

14. Luck, S. (2022). Applied event-related potential data analysis. LibreTexts.[Google Scholar], .

15. Luck, S. J. (2005). An introduction to the event-related potential technique. MIT press.

16. Luck, S. J. (2014). An introduction to the event-related potential technique. MIT press.

17. Luck, S. J., Stewart, A. X., Simmons, A. M., & Rhemtulla, M. (2021). Standardized measurement error: A universal metric of data quality for averaged event-related potentials. Psychophysiology, 58, e13793. doi:10.1111/psyp.13793.

18. Rousselet, G. A. (2012). Does filtering preclude us from studying erp time-courses? Frontiers in Psychology, 3, 131. doi:10.3389/fpsyg.2011.00365.

19. Schindler, S., & Bublatzky, F. (2020). Attention and emotion: An integrative review of emotional face processing as a function of attention. Cortex, 130, 362–386. doi:10.1016/j.cortex.2020.06.010.

20. Shad, E. H. T., Molinas, M., & Ytterdal, T. (2020). Impedance and noise of passive and active dry EEG electrodes: a review. IEEE Sensors Journal, 20, 14565–14577. doi:10.1109/JSEN.2020.3012394.

21. Tanner, D., Morgan-Short, K., & Luck, S. J. (2015). How inappropriate high-pass filters can produce artifactual effects and incorrect conclusions in ERP studies of language and cognition. Psychophysiology, 52, 997–1009. doi:10.1016/j.jneumeth.2016.01.002.

22. Tanner, D., Norton, J. J., Morgan-Short, K., & Luck, S. J. (2016). On high-pass filter artifacts (they’re real) and baseline correction (it’sa good idea) in ERP/ERMF analysis. Journal of Neuroscience Methods, 266, 166–170. doi:10.1111/psyp.12437.

23. VanRullen, R. (2011). Four common conceptual fallacies in mapping the time course of recognition. Frontiers in Psychology, 2, 365. doi:10.3389/fpsyg.2011.00365.

24. Yeung, N., Bogacz, R., Holroyd, C. B., Nieuwenhuis, S., & Cohen, J. D. (2007). Theta phase resetting and the error-related negativity. Psychophysiology, 44, 39–49. doi:10.1111/j.1469-8986.2006.00482.x.

25. Zhang, G., Garrett, D. R., & Luck, S. J. (2023). Optimal filters for erp research i: A general approach for selecting filter settings. bioRxiv, (pp. 2023–05). doi:10.1101/2023.05.25.542359.

26. Zhang, G., & Luck, S. J. (2023). Variations in ERP data quality across paradigms, participants, and scoring procedures. Psychophysiology, 60, e14264. doi:10.1111/psyp.14264.

